# Brain-epigenome wide association study (BEWAS) on the effects of two emerging psychedelics: ketamine & MDMA

**DOI:** 10.1101/2025.07.03.663007

**Authors:** Moira G. Semple, Sarah E. Mennenga, Ryan Smith, Varun B. Dwaraka, Baruch Rael Cahn, Joseph Tafur, David M. Rabin, Berra Yazar-Klosinski, Candace R. Lewis

**Affiliations:** School of Life Sciences, Arizona State University; Tempe, AZ, USA; TruDiagnostic, Lexington, KY 40503, USA; Department of Psychiatry and Behavioral Sciences, University of Southern California, Los Angeles, CA, United States; Modern Spirit, Phoenix, AZ, United States; The Board of Medicine, Pittsburgh, PA, United States; Independent Researcher, Watson, CA; Neurogenomics Division, Translational Genomics Research Institute (TGen); Phoenix, AZ, USA

## Abstract

Psychedelic compounds such as ketamine and MDMA have shown therapeutic promise for mood and trauma-related disorders, yet their molecular mechanisms remain unclear. This study applied a Brain-Epigenome-Wide Association Study (BEWAS) to assess DNA methylation changes in brain-enriched genes following treatment. Pre- and post-treatment blood (ketamine, *N* = 20) and saliva (MDMA, *N* = 16) samples from clinical trial participants were analyzed. Ketamine altered methylation at 1,210 CpG sites; MDMA affected 2,074 CpG sites. Functional enrichment analyses revealed changes in genes involved in neuroplasticity, immune regulation, and mental processes. Overlapping effects were observed in genes such as *PTPRN2* and *SHANK2*, suggesting shared epigenetic mechanisms in driving increased neuroplasticity. These findings highlight psychedelics’ capacity to induce coordinated, lasting molecular changes relevant to neuroimmune function and psychiatric health.

## INTRODUCTION

The incidence of mental health disorders is increasing globally(*1*, *2*), and treatment rates remain low, resulting in an urgent need for new mental health interventions(*3*). Scientific and clinical interest in psychedelics is high(*4*) following recent clinical trials that support their broad therapeutic potential (e.g.,(*5*, *6*)). The clinical benefits of psychedelics are widely attributed to pro-neuroplastic(*7*) and neuroimmune(*8*) effects; however, the mechanism(s) by which they affect mental health remain unclear. Ketamine and 3,4-methylenedioxymethamphetamine (MDMA) are two prominent psychedelics under investigation for treatment of similar mood- and anxiety-related symptoms that act via distinct and complex pharmacological mechanisms.

Ketamine has been shown to rapidly reduce symptoms of Major Depressive Disorder (MDD)(*9*, *10*), and meta-analysis supports its potential as a treatment for Post-Traumatic Stress Disorder (PTSD)(*6*). Ketamine produces dissociation at high doses, or euphoria and mild sensory alterations at lower doses, via antagonism of N-methyl-D-aspartate (NMDA) receptors(*10*), which have been widely studied for their role in synaptic plasticity(*11*). Ketamine’s NMDA effects alter GABAergic and glutamatergic neurotransmission(*12*), which has been linked to enhanced neuroplasticity and improvements in mood(*13*). However, ketamine affects many other key brain systems as well, including dopaminergic(*14*), opioidergic(*15*), serotonergic(*16*), stress(*17*), and immune pathways(*18*). Thus, clinical benefits of ketamine could arise through effects on virtually any neurocircuit and may involve regulation of multiple systems.

MDMA has also been shown to regularize negative affect circuit activity(*19*) and effectively treat PTSD symptoms when combined with psychotherapy, outperforming psychotherapy alone(*20*, *21*). MDMA produces empathogenic, sensory-altering, and mood-enhancing psychedelic effects, primarily via actions at the serotonin transporter that enhance serotonergic signaling. However, MDMA is also a potent releaser/reuptake inhibitor of norepinephrine and dopamine(*22*), likely affecting mood, motivation, and reward processes(*23*). MDMA also has anti-inflammatory properties(*24*), promotes release of the empathogenic neuropeptide oxytocin(*25*), and is a potent activator of the hypothalamic pituitary adrenal stress system(*26*). Thus, like ketamine, MDMA’s therapeutic effects are likely complex, and may involve convergence of effects across multiple pathways.

The longevity of symptom reduction following only a few psychedelic treatments suggests that exposure to these drugs produces lasting neurobiological change, such as by affecting molecular processes that regulate gene accessibility for transcription. Changes in DNA methylation, the addition/removal of methyl groups at cytosine guanine dinucleotide (CpG) sites on the DNA(*27*), have been repeatedly linked to mental health and psychiatric treatment success. In fact, DNA methylation from peripheral samples (e.g., blood or saliva) is increasingly utilized as a biomarker for psychiatric treatment response(*28*), as it offers an effective and non-invasive way to predict treatment outcomes(*29*). Ketamine administration has been shown to induce RNA changes in brain(*30*) and DNA methylation changes in blood (preprint(*31*)). In a small convenience sample, our group recently observed that changes in methylation at CpG sites on cortisol-regulating genes may predict PTSD symptom reduction following treatment with MDMA assisted therapy, providing preliminary support for further investigation(*32*).

We now expand these findings by conducting the first comprehensive evaluation of ketamine and MDMA effects on DNA methylation, using an analytical method that we termed a *Brain-Epigenome-Wide Association Study (BEWAS)*. This approach combines the strengths of Candidate Gene and Epigenome-Wide Association Study (EWAS) methods. Candidate Gene approaches, where pre-selected genes are studied based on their biological relevance to a trait or disease, are cost-effective, require smaller datasets, and are suitable for hypothesis-driven research, but they are limited by existing knowledge and can omit important genes. In contrast, EWAS takes an unbiased, hypothesis-free approach by examining the entire epigenome to identify associations with traits or diseases. While EWAS is a powerful tool for discovery, it requires large datasets and incurs high statistical penalties to combat alpha inflation. The BEWAS approach reduces the required number of statistical tests by focusing on methylation at or near genes known to be highly expressed in the brain, thereby limiting alpha inflation without requiring hypothesis-driven gene candidate selection.

## RESULTS

### Participant Demographics

Ketamine and MDMA participants were similar in age (ketamine: M[SD]=40.24[12.12]; MDMA: M[SD]=43.49[11.83]), however there was a greater representation of females in the ketamine cohort (ketamine: 25% male; MDMA 56% male). See Figure 1 for the study analyses work flow.

**Fig. 1.**
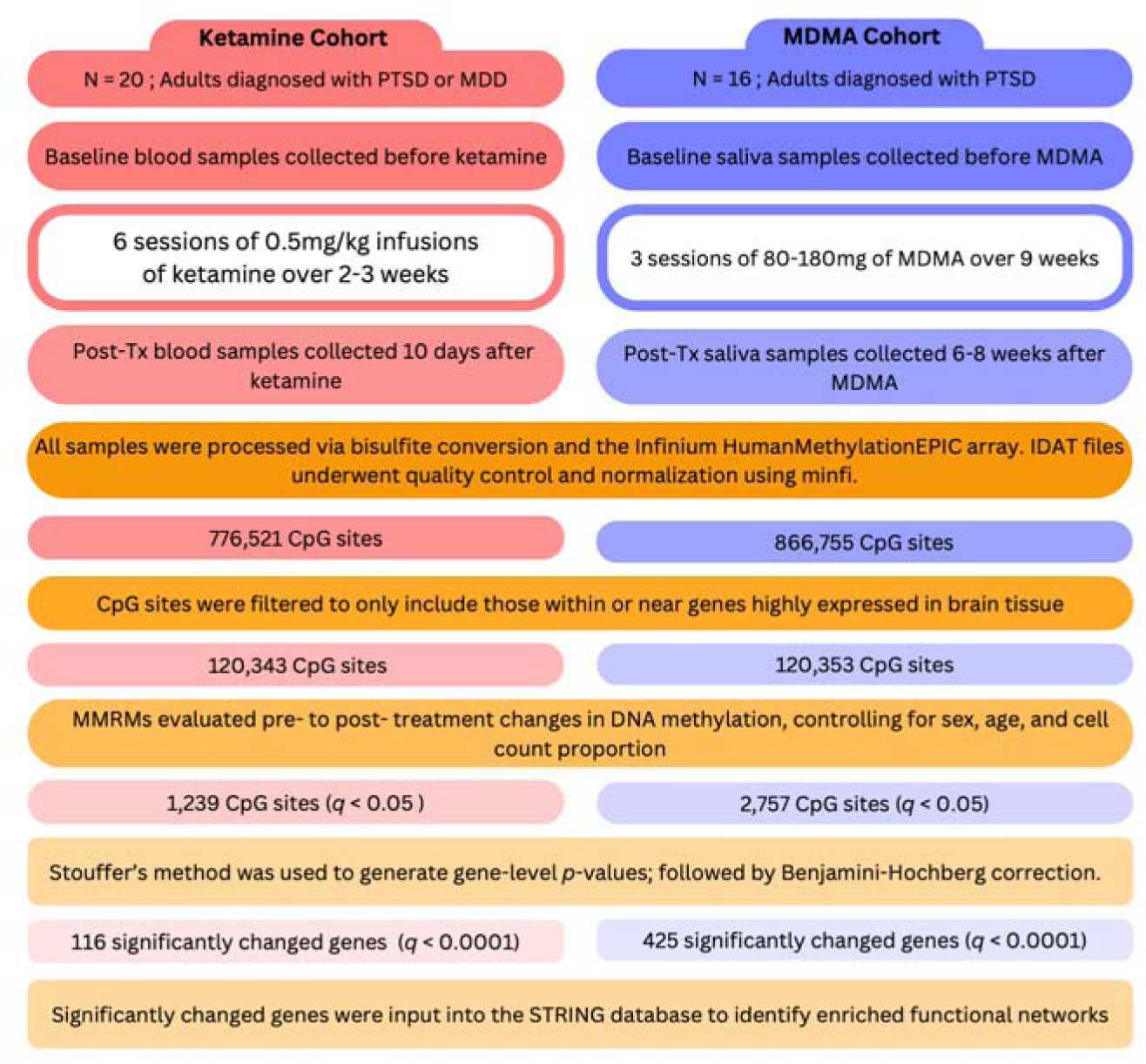
Summary of study methods across two cohorts. Flowchart depicting methodology for drug administration, biological sample collection, DNA isolation, as well as differential DNA methylation analysis across two cohorts. Type II ANCOVA was used in this study as the MMRM (Mixed model for repeated measures) which was followed by the Benjamini-Hochberg correction.

### Ketamine - Tests of CpG and gene level effects

Of the 1,210 brain-enriched CpG sites (within or near 788 genes) that demonstrated methylation change from pre- to post-ketamine treatment, 72.3% showed increased methylation and 27.7% showed decreased methylation (Figure 2A) and 82.3% had an absolute β value greater than 0.01 (Supplemental Figure 1A). CpG sites that changed with ketamine treatment were located within, upstream, and downstream of promotors, with the greatest number in Open Sea (Table 1; complete list in Supplemental Table 1). FDR-corrected Stouffer-Lipták-Kechris gene-level analyses revealed 790 genes that were significantly altered from pre- to post-ketamine treatment (*q* < 0.01), and 126 genes at a stricter threshold (*q* < 0.0001; Supplemental Table 2).

**Fig. 2.**
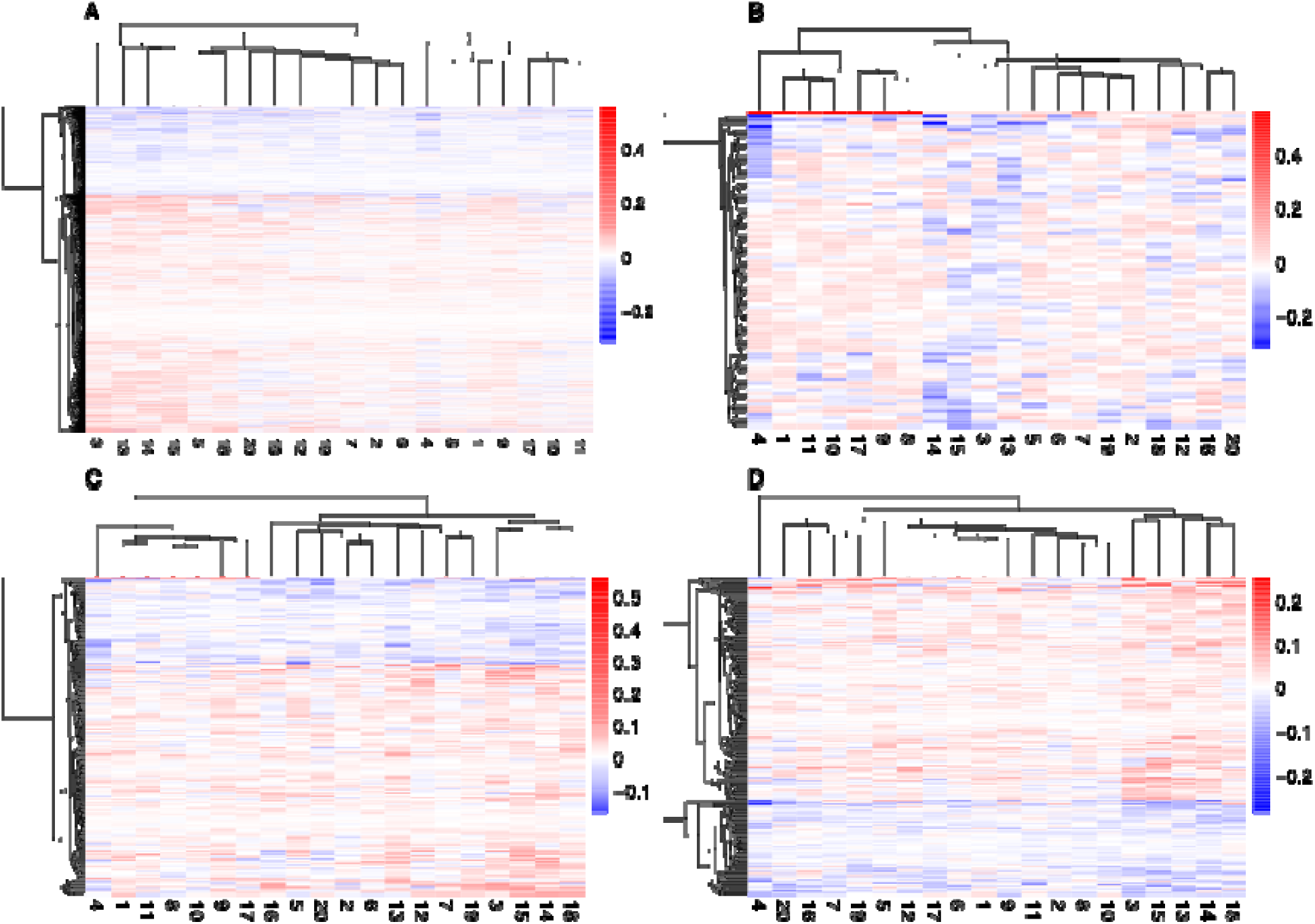
Ketamine induced change of DNA methylation across individuals and across mental health related systems. **(A)** Columns represent individuals (N = 20), rows represent CpG sites altered after ketamine. treatment (N = 1,210; *p* < 0.01). Cell color depicts the change if methylation from pre- to post-ketamine treatment. Negative values depict a decrease in methylation (blue) and positive values depict an increase in methylation (red). Heatmaps were generated using the package “pheatmap” in R version 4.4.1. with hierarchical clustering along each axis. **(B**) Neurotransmission/plasticity N = 89 CpG sites. **(C)** Immune system; N = 197 CpG sites. **(D)** Mental processes; N = 218 CpG sites.

**Table 1.**
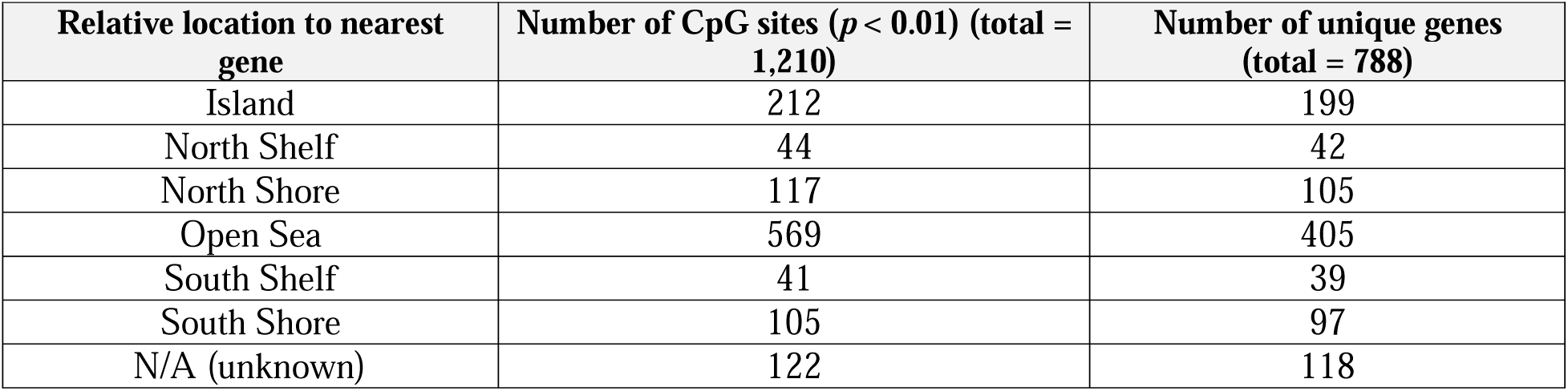
Location and count of CpG sites altered by ketamine.

| Relative location to nearest gene | Number of CpG sites ($p < 0.01$) (total = 1,210) | Number of unique genes (total = 788) |
| --- | --- | --- |
| Island | 212 | 199 |
| North Shelf | 44 | 42 |
| North Shore | 117 | 105 |
| Open Sea | 569 | 405 |
| South Shelf | 41 | 39 |
| South Shore | 105 | 97 |
| N/A (unknown) | 122 | 118 |

### Ketamine - Tests of neuroplasticity, immune function, and mental processes hypotheses

Of the genes that changed from pre- to post-ketamine treatment (*q* < 0.0001), 24 (19%) were related to neuroplasticity (Supplemental Table 3), and we identified 24 functional enrichment networks (Supplemental Table 4), of which six (25%) were related to immune function and five (20.8%) were related to mental processes (Table 2; Figure 2B-D). Number of genes, signal amplitude, and strength of select significant functional enrichments are displayed in Figure 3.

**Fig. 3.**
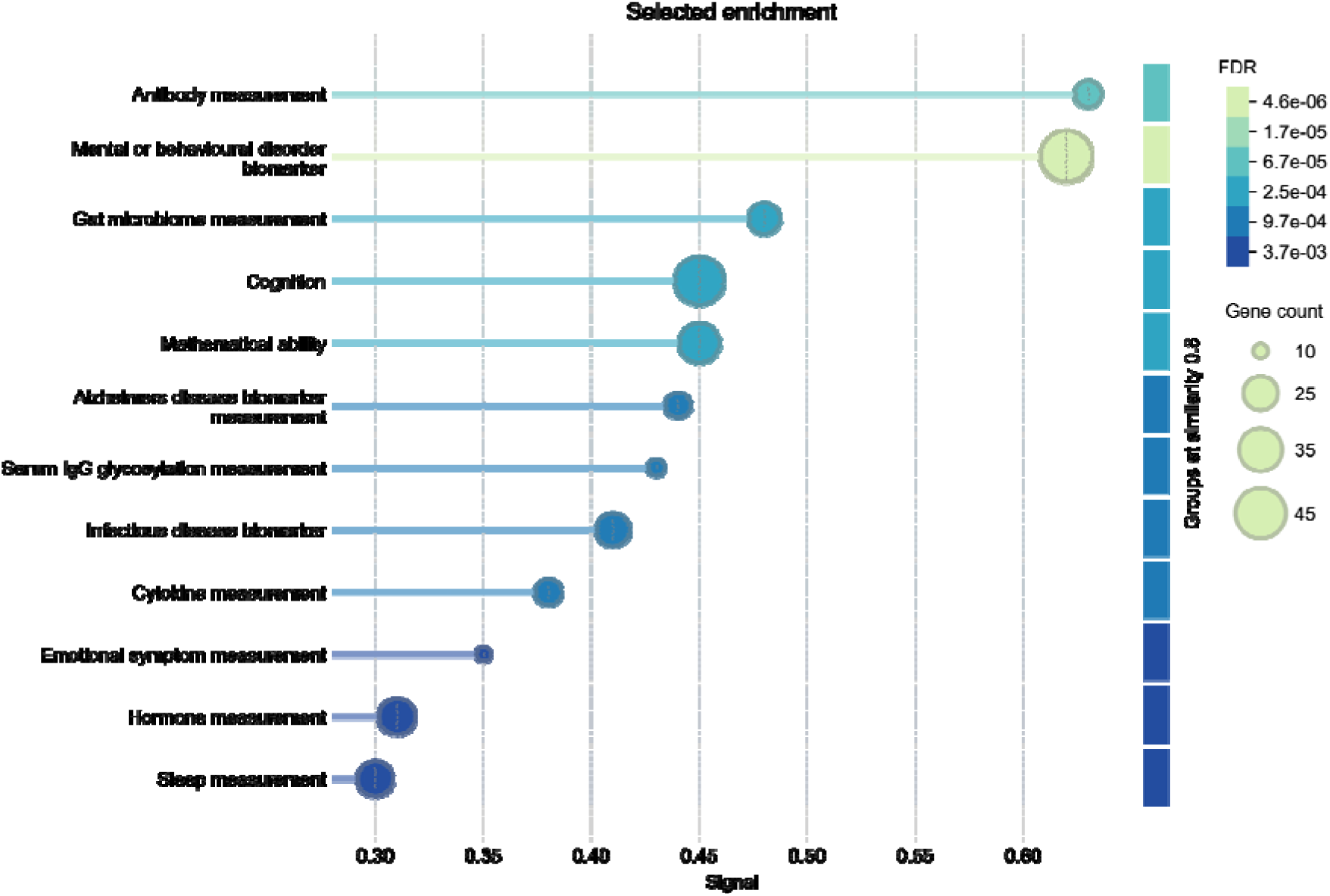
Ketamine significantly alters the DNA methylation of genes associated with various functional enrichment networks. Out of the 24 functional enrichments detected from the list of genes significantly altered post-ketamine, 12 were chosen based on their association with the hypothesized mechanisms. For the full list of functional enrichment networks and the associated genes within each network see Supp Table 3.

**Table 2.** Number of ketamine-affected genes associated with neuroplasticity, immune function, and mental processes (*q* < 0.0001)

| Hypothesis | | Gene count ($q < 0.0001$) |
| --- | --- | --- |
| Neuroplasticity | Synaptic Plasticity | 24 |
|  | Neurotrophins/Receptors |  |
|  | Neurotransmission |  |
|  | **Functional Enrichment Networks** | **Gene count ($q < 0.0001$)** |
| Immune Function | Antibody measurement | 18 |
|  | Gut microbiome measurement | 22 |
|  | Infectious disease biomarker | 24 |
|  | Serum IgG glycosylation measurement | 12 |
|  | Cytokine measurement | 18 |
|  | Disease | 26 |
| Mental Processes | Mental or behavioral disorder biomarker | 46 |
|  | Cognition | 42 |
|  | Mathematical ability | 31 |
|  | Alzheimer's disease biomarker measurement | 17 |
|  | Emotional symptom measurement | 11 |

**Table 3.** Location and count of CpG sites altered by MDMA.

| Relative location to nearest gene | Count of significant CpG sites ($p < 0.01$) (total = 2,074) | Number of unique genes (total = 1,027) |
| --- | --- | --- |
| Island | 253 | 223 |
| North Shelf | 104 | 92 |
| North Shore | 197 | 164 |
| Open Sea | 1,233 | 699 |
| South Shelf | 67 | 61 |
| South Shore | 164 | 149 |
| N/A (unknown CpG site relative location) | 56 | 51 |

### MDMA - Tests of CpG and gene level effects

Of the 2,074 brain-enriched CpG sites (within or near 1,027 genes) that demonstrated methylation change from pre- to post-MDMA treatment, 48.3% had increased methylation and 51.7% had decreased methylation (Figure 4A), and 91.4% had an absolute β value greater than 0.01 (Supplemental Figure 1B). CpG sites that changed with MDMA treatment were also located within, upstream, and downstream of promotors, and the greatest number was again in Open Sea (Table 3; complete list in Supplemental Table 5). FDR-corrected Stouffer-Lipták-Kechris gene-level analyses revealed 1,027 genes that were significantly altered from pre- to post-MDMA treatment (*q* < 0.01), and 344 genes at a stricter threshold (*q* < 0.0001; Supplemental Table 6).

**Fig. 4.**
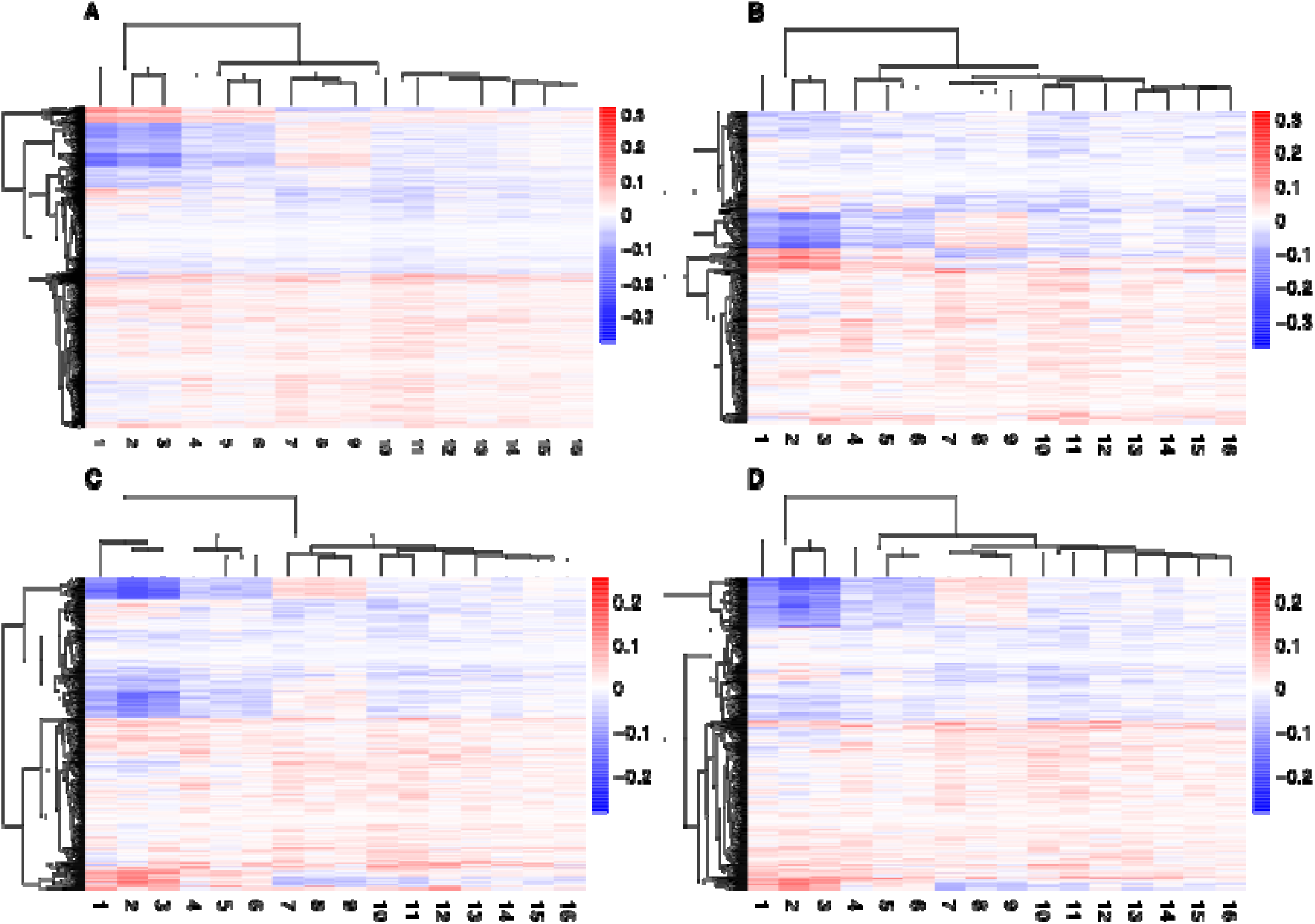
MDMA-induced change of DNA methylation across individuals and across mental health related system. **(A)** Columns represent individuals (N = 16), rows represent CpG sites altered after ketamine treatment (N = 2,074; *p* < 0.01). Cell color depicts the change if methylation from pre- to post-ketamine treatment. Negative values depict a decrease in methylation (blue) and positive values depict an increase in methylation (red). Heatmaps were generated using the package “pheatmap” in R version 4.4.1. with hierarchical clustering along each axis. **(B**) Neurotransmission/plasticity N = 440 CpG sites. **(C)** Immune system; N = 596 CpG sites. **(D)** Mental processes; N = 599 CpG sites.

### MDMA - Tests of neuroplasticity, immune function, and mental processes hypotheses

Of the genes that changed from pre- to post-MDMA treatment (*q* < 0.0001), 69 (20%) were related to neuroplasticity/neurotransmission (Supplemental Table 7), and STRING(*33*) identified 46 significant functional enrichment networks (Supplemental Table 8), of which six (13%) were related to immune function, and five (10.8%) were related to mental processes (Table 4; Figure 4B-D). Number of genes, signal amplitude, and strength of select significant functional enrichments are displayed in Figure 5.

**Fig. 5.**
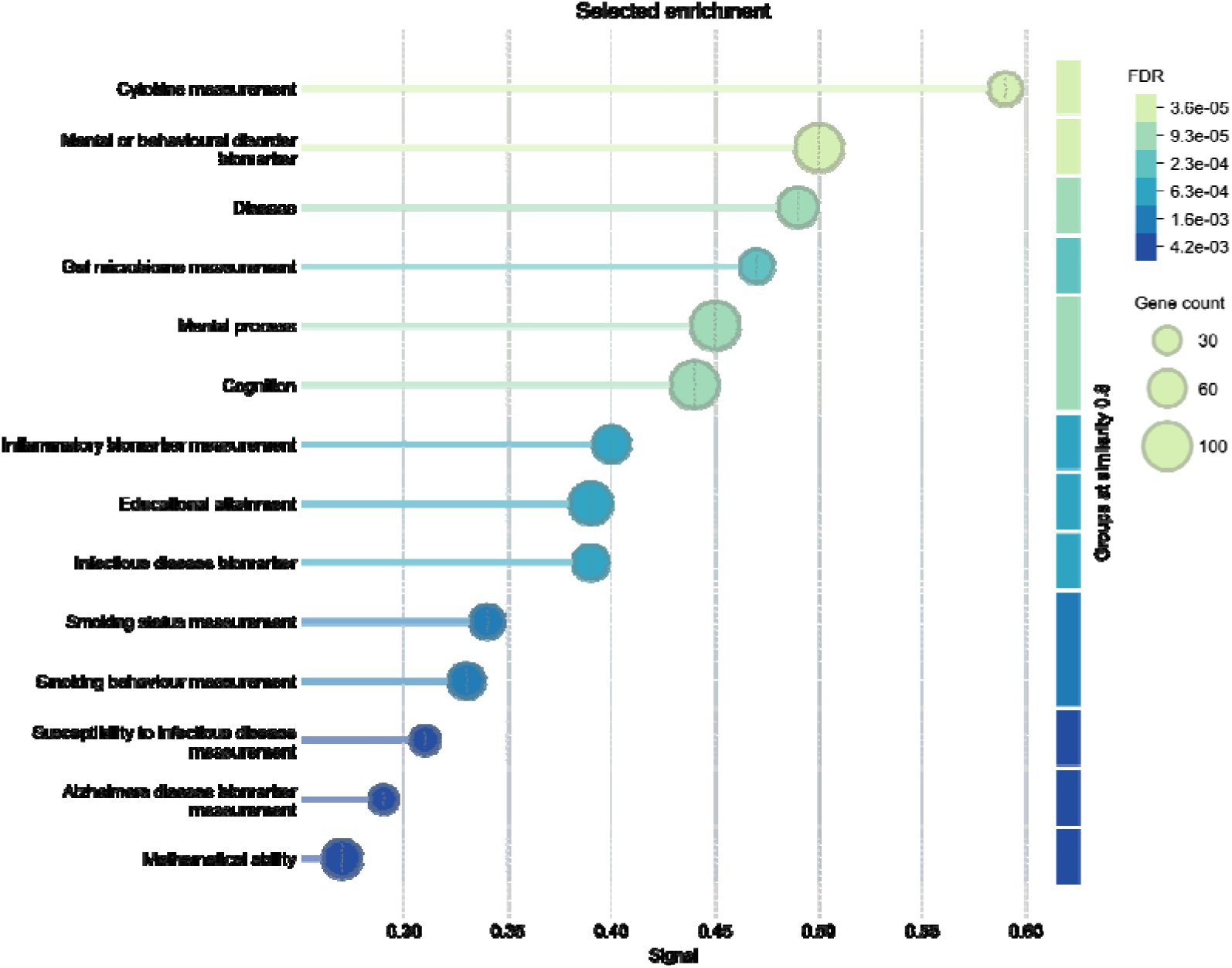
MDMA significantly alters the DNA methylation of genes associated with various functional enrichment networks. Out of the 46 functional enrichments detected from the list of genes significantly altered post-MDMA, 14 were chosen based on their association with the hypothesized mechanisms. For the full list of functional enrichment networks and the associated genes within each network see Supp Table 6.

**Table 4.** Number of MDMA-affected genes associated with neuroplasticity, immune function, and mental processes (*q* < 0.0001)

| Hypothesis | | Gene count ($q < 0.0001$) |
| --- | --- | --- |
| Neuroplasticity | Synaptic Plasticity | 69 |
|  | Neurotrophins/Receptors |  |
|  | Neurotransmission |  |
|  | **Functional Enrichment Networks** | **Gene count ($q < 0.0001$)** |
| Immune Function | Cytokine measurement | 44 |
|  | Disease | 65 |
|  | Gut microbiome measurement | 46 |
|  | Inflammatory biomarker measurement | 54 |
|  | Infectious disease biomarker | 51 |
|  | Susceptibility to infectious disease biomarker | 33 |
| Mental Processes | Mental or behavioral disorder biomarker | 94 |
|  | Mental process | 97 |
|  | Cognition | 95 |
|  | Educational attainment | 73 |
|  | Smoking status measurement | 46 |
|  | Smoking behavior measurement | 51 |
|  | Alzheimer's disease biomarker measurement | 31 |
|  | Mathematical ability | 61 |

### Ketamine and MDMA Overlap

There was substantial overlap in DNA methylation changes associated with ketamine and MDMA treatment; however, MDMA alone was associated with a larger number of differentially methylated CpG sites than either ketamine alone or both drugs (Figure 6). Of the 126 ketamine-affected and 344 MDMA-affected genes, we identified 12 genes with 10 or more CpG sites that were significantly affected by either ketamine or MDMA, including at least three significantly altered sites per drug (Table 5). Lastly, both ketamine and MDMA influenced a substantial number of overlapping functional networks (Figure 7).

**Fig. 6.**
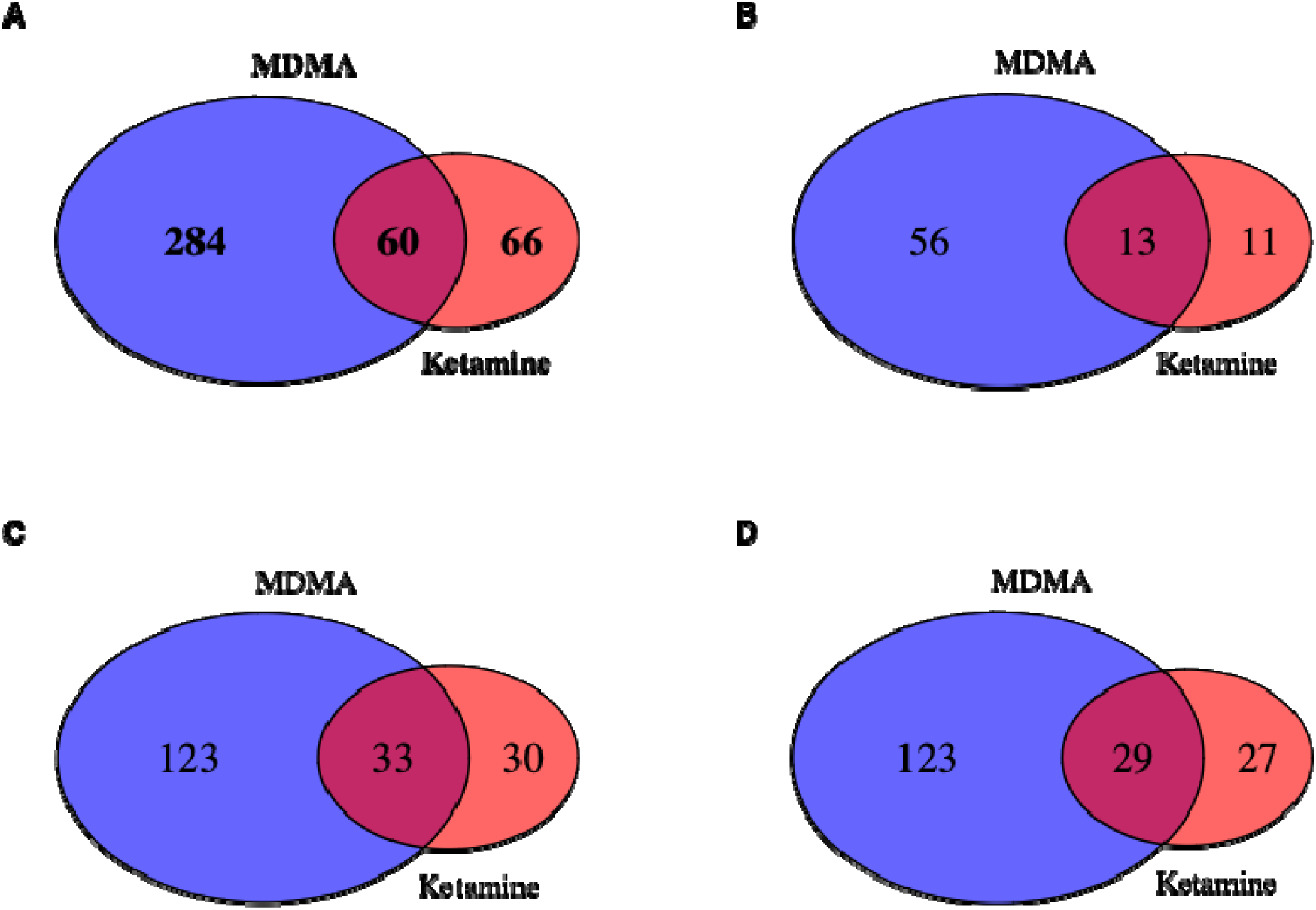
Ketamine and MDMA alter DNA methylation across the same and distinct genes. Stouffer-Lipták-Kechris adjustment followed by FDR correction was used to generate gene level results for ketamine and MDMA (*q <* 0.0001). The package “VennDiagram” in R version 4.4.1 was used to generate figures. **(A)** Overlapping significance across all genes (q < 0.0001). **(B)** The list of genes associated with neurotransmission/plasticity was compiled from numerous sources. For the **(C)** immune system and **(D)** mental process diagrams, the lists of genes were obtained using the STRING network results, by combining the genes in each network associated with either immune health or mental process/behavioral phenotypes.

**Fig 7.**
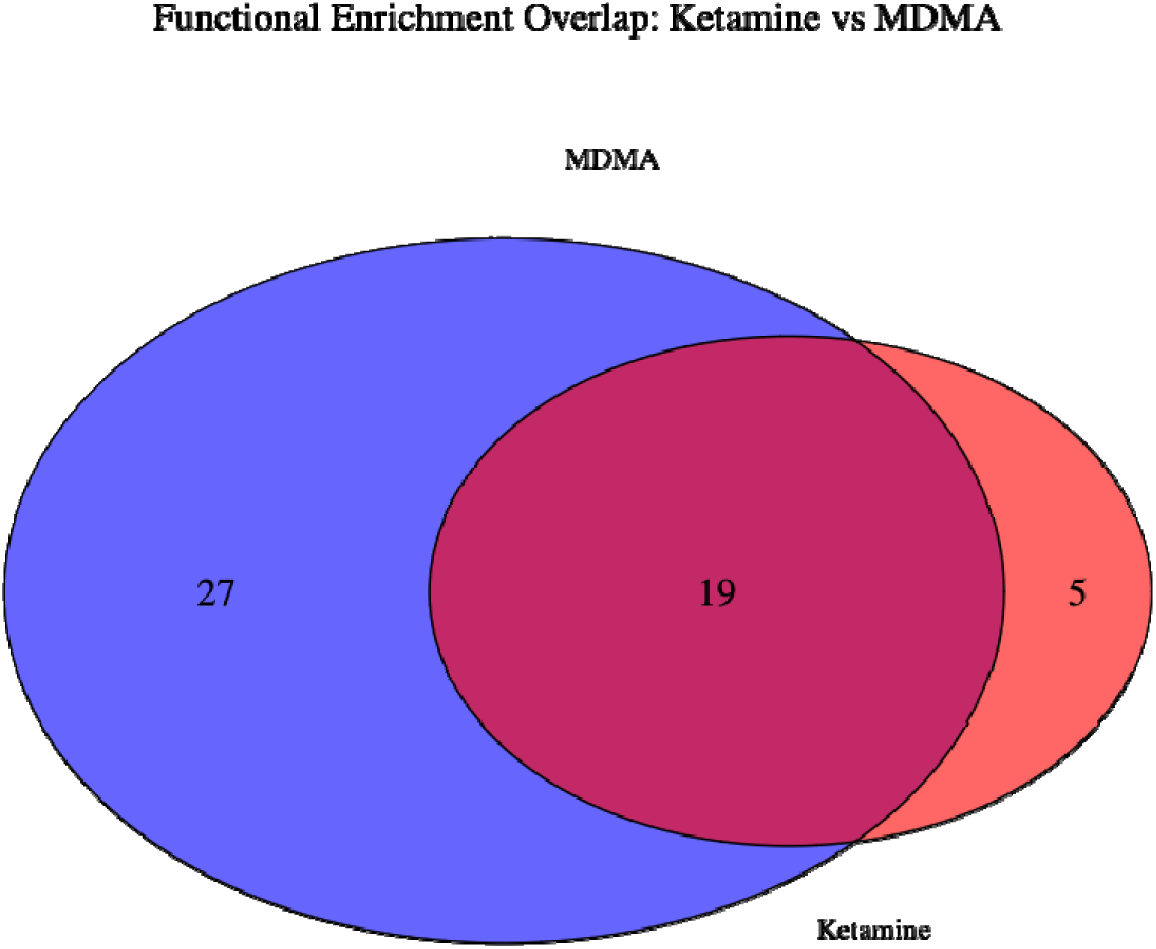
Overlap of functional enrichments following ketamine and MDMA exposure. *Shared enrichments (n =19):* Mental or behavioural disorder biomarker, Body mass index, Temporal measurement, Body weights and measures, Anthropometric measurement, Cognition, Protein measurement, Self reported educational attainment, Mathematical ability, Gut microbiome measurement, Population measurement, Alzheimers disease biomarker measurement, Infectious disease biomarker, Phenotypic abnormality, Cytokine measurement, Disease, Hormone measurement, Repeat, Alternative splicing. *Unique to ketamine (n=5):* Antibody measurement, Blood toxic metal measurement, Serum IgG glycosylation measurement, Emotional symptom measurement, Sleep measurement. *Unique to MDMA (n = 27):* Cardiovascular measurement, Vital signs, Cardiovascular disease biomarker measurement, Heart function measurement, Mental process, Blood pressure, Respiratory disease biomarker, Metabolite measurement, Pulmonary function measurement, Body height, Educational attainment, Inflammatory biomarker measurement, Lipid or lipoprotein measurement, Complete blood cell count, Electrocardiography, Smoking status measurement, Smoking behaviour measurement, Lipid measurement, Eye measurement, FEV/FEC ratio, Systolic blood pressure, Susceptibility to infectious disease measurement, Abnormality of the cardiovascular system, Testosterone measurement, Erythrocyte count, Brain measurement, Metal-binding.

**Table 5.** Genes with High Recurrence in both Ketamine and MDMA CpG Site Data*.

| Gene Name | Count of significant CpG sites from ketamine cohort | Count of significant CpG sites from MDMA cohort |
| --- | --- | --- |
| *PTPRN2* | 20 | 27 |
| *SHANK2* | 6 | 12 |
| *NTM* | 6 | 9 |
| *ABR* | 5 | 10 |
| *SALL3* | 5 | 7 |
| *CBFA2T3* | 4 | 8 |
| *ADARB2* | 5 | 6 |
| *KAZN* | 5 | 5 |
| *PRKAR1B* | 5 | 5 |
| *SORCS2* | 5 | 5 |
| *BCL11B* | 4 | 6 |
| *GNG7* | 4 | 6 |

## DISCUSSION

Our findings reveal large-scale epigenetic modifications related to neuroplasticity, immune function, and mental health processes following treatment with two promising new psychedelics. These results suggest that the therapeutic effects of ketamine and MDMA may be mediated by epigenetic mechanisms, and support the growing evidence that psychedelic-assisted treatments exert their effects through complex, multi-system interactions(*34–36*).

The broad epigenetic changes we found are in agreement with large-scale shifts in gene expression that have been previously reported with repeated ketamine treatment(*37*, *38*), and align with known pharmacological effects of ketamine and MDMA. Here, ketamine exposure was associated with DNA methylation changes in 126 genes, many of which are linked to NMDA receptor function and glutamate signaling (e.g., *GRIN1*, *GRIN2B*, *GRIN2D*, *GRIN3A*, as well as *GRIA1*, *GRIA3*, *GRIK2*, *GRIK3*, *GRID1*, *GRID2*, and *SLC1A3*; see Supplemental Table 3). MDMA treatment was associated with changes in more than twice as many genes as ketamine overall, including some classically associated with serotonergic signaling (e.g., *HTR1B* and *HTR2A*; see Supplemental Table 6). These differences may reflect the presence of psychotherapeutic engagement in the MDMA-assisted therapy model, in contrast to the primarily pharmacological ketamine intervention. Additionally, the greater clinical heterogeneity in the ketamine sample (MDD + PTSD) may have contributed to increased baseline epigenetic variability and limited the detection of consistent methylation changes.

We found ketamine-related methylation changes in several genes that are widely studied for their role in neuroplasticity, including *NEGR1,* a neuronal growth regulator, and *RASGRF1*, a regulator of the MAPK/ERK pathway known for its role in synaptic plasticity, memory formation, and neuronal excitability that has been implicated in the response to ketamine treatment(*39*). These findings add to an increasing body of evidence suggesting that ketamine’s antidepressant effects involve changes in synaptic plasticity(*40*, *41*). We also found MDMA-induced methylation changes in plasticity-related genes, including several involved in synapse formation (e.g. *SHANK2, NRXN1, PRKCZ*), synaptic vesicle trafficking (e.g. *SYN3, CADPS2, SNCA, PRKCA*), and glutamatergic signaling (e.g. *GRIK1, GRID1, SLC1A2 and SLC1A3*; see Supplemental Table 6), further bolstering the general neuroplasticity hypothesis of psychedelic treatment. Notably, DNA methylation at *SHANK2* has been previously associated with PTSD in large-scale epigenome-wide association discovery and validation cohorts(*42*), further supporting its potential relevance as a treatment target.

Ketamine and MDMA exposure were also both associated with altered methylation of genes related to immune function. Functional enrichment analysis revealed ketamine- and MDMA-related change in networks associated with cytokine regulation, gut microbiome, and disease susceptibility (Tables 2 and 4; Supplemental Tables 4 and 8). Prior evidence of ketamine’s and MDMA’s immunomodulatory effects are in agreement with the current findings(*8*, *18*, *43*), and changes in inflammatory markers have been widely implicated in mood disorders(*44*), strengthening the hypothesis that immune regulation is an additional key part of the therapeutic response to psychedelics.

Both drugs were also associated with alterations in genes that have been implicated in mental health-related phenotypes, including cognition and mental/behavioral disorders (Tables 2 and 4). MDMA, but not ketamine, was also linked to several genes related to smoking behavior; an interesting finding in the context of ongoing trials of psilocybin, another serotonergic psychedelic, for smoking cessation(*45–47*). These broad DNA methylation changes may, in part, be a molecular basis for the transdiagnostic therapeutic efficacy of psychedelic drugs in general.

Despite their distinct pharmacological profiles, ketamine and MDMA exhibited overlapping effects on DNA methylation, particularly in genes linked to neuroplasticity. Genes that showed methylation changes following treatment with both drugs (*PTPRN2, NTM, SHANK2, ABR*) implicate vesicle dynamics and synaptic remodeling as potential shared neuroplastic mechanisms(*48*, *49*). While both substances influenced immune-related pathways, ketamine demonstrated a stronger effect on inflammatory biomarker regulation, whereas MDMA had greater involvement in cytokine modulation (Figures 3 and 5). These differences may reflect distinct mechanisms of action on immune-related pathways.

Several limitations to our approach must be acknowledged. Our relatively small sample size across both trials necessitates replication in larger cohorts to confirm these results. Also, while peripheral DNA methylation has been shown to reflect differences in brain structure(*50*–*52*), future studies should aim to validate these findings in animal models that permit collection of brain tissue. It is also important to note that methylation was assessed in different peripheral tissues across the two trials (blood for ketamine and saliva for MDMA), which may introduce tissue-specific variation. Nevertheless, the high degree of overlap observed in enriched pathways across these distinct tissues is notable and may point to robust, convergent biological processes engaged by both interventions. Additionally, while this study demonstrates associations between psychedelic treatment and DNA methylation changes, directionality and causality remain unclear. These findings do not clarify whether neuroplasticity- or neuroimmune-related DNA methylation changes lead to up- or down-regulated plasticity. This distinction is critical, as prior research suggests that high acute doses of MDMA may suppress neurogenesis (for review see(*7*)). Longitudinal studies tracking epigenetic changes alongside other neurobiological and clinical changes will be critical in determining whether these modifications directly contribute to symptom improvement. Lastly, while our BEWAS approach was designed to identify changes in methylation at genes that had relatively high expression in brain tissue, other approaches to honing epigenome-wide changes are also valid and warranted, especially given the known effects psychedelics have on peripheral tissues like the cardiovascular system(*53*) and gut microbiome(*54*). This approach could also be extended to study other psychedelics, such as psilocybin and LSD, to further elucidate and compare their mechanisms of action.

## MATERIALS AND METHODS

### Study Design

We analyzed DNA methylation data from participants enrolled in clinical trials for either ketamine or MDMA. All participants provided biospecimens (either blood or saliva) before and after their respective psychedelic treatment (Figure 1). We use a BEWAS approach to assess epigenetic change across genes that have relatively high expression in brain, using measurements from peripheral samples.

### Ketamine Participants

Twenty participants with moderate-severe MDD or PTSD were recruited for a study on the epigenetic effects of ketamine (for full study details(*31*)). Participants received 6 ketamine infusions (0.5mg/kg) over the course of 2-3 weeks. Peripheral whole blood samples were collected before and after treatment, using the standard lancet and capillary method. DNA was isolated and underwent bisulfite conversion using the EZ DNA Methylation Kit (Zymo Research).

### MDMA Participants

A convenience sample of sixteen men and women with severe PTSD were recruited as part of a phase 3 trial of MDMA for PTSD (for full study details(*20*)). Participants received three oral doses of MDMA (80-180mg) approximately four weeks apart, along with psychotherapy (Figure 1). Saliva samples were collected one week before the first MDMA treatment and six-eight weeks after the last MDMA treatment, with Oragene-DNA saliva kits. DNA was isolated with a DNA Genotek isolation kit (PT-L2P; Ottawa, Ontario, Canada) and underwent bisulfite conversion using the EZ DNA Methylation Kit (Zymo Research).

### DNA Methylation Analysis

DNA was processed on the Infinium HumanMethylationEPICv1 BeadChip with standard protocols, and raw image intensities were captured using the Illumina iScan System. Raw Intensity Data (IDAT) files were preprocessed utilizing the Minfi package in R (for more details(*31*)). Probes known to be cross-reactive, located on sex chromosomes, and near SNPs were removed. Quality control analyses included quantile normalization, checking for sex mismatches, and excluding low-intensity samples (*p* < 0.01)(*32*). Methylation values were normalized and missing CpG values for ketamine samples were imputed using the k-nearest neighbors (k-NN) algorithm. The R package EpiDISH (Epigenetic Dissection of Intra-Sample Heterogeneity, 3.8) Robust Partial Correlation (RPC) method was used to generate the estimated proportion of epithelial cells.

### CpG filtering for BEWAS analysis

We used publicly available data to identify 2,197 genes that have elevated expression in brain tissue, compared to other tissue types (The Human Protein Atlas, n.d.). We then restricted the processed DNA methylation data to sites on or near genes in this list, reducing the number of CpG sites for analysis by roughly 85%.

### Tests of CpG and gene level effects

To assess the effects of each intervention, we constructed mixed models for repeated measures (MMRMs) using the package lmerTest in R (version 4.4.1)(*55*), including pre- and post-treatment DNA methylation as a repeated measure and fixed effects of age, sex, and epithelial cell count. Psychedelic-affected CpG sites (*p* < 0.01) were then aggregated and assessed for gene-level significance using the Stouffer-Lipták-Kechris adjustment (autocorrelation-adjusted z-test)(*56*, *57*). Genes with a *q* < 0.0001 were considered significant, using the Benjamini-Hochberg correction for false discovery(*58*).

### Tests of neuroplasticity, immune function, and mental processes hypotheses

We also took a hypothesis-driven approach to assess the impact of psychedelics on brain-enriched genes known to be involved in neuroplasticity, immune function, and mental processes. To generate a neuroplasticity gene list, we cross-referenced data from commercial sources and the literature(*59*) with our BEWAS gene list (Supp. Tables 1 and 5). To assess effects on immune function and mental processes, we conducted functional enrichment analyses using STRING v12.0 bioinformatics platform (https://string-db.org/)(33), with a BEWAS gene list background. Enriched functional networks were considered significant with a *q* < 0.05.

## List of Supplementary Materials

Fig. S1A-B. Volcano plots of DNA methylation ANCOVA results showing –log (p-value) versus effect size for all CpG sites included in BEWAS for Ketamine and MDMA.

Table S1. Complete list of CpG sites included in ketamine BEWAS analysis with estimate and p-value results

Table S2. List of gene-level results post-Ketamine

Table S3. List of genes associated with neurotransmission/neuroplasticity post-ketamine

Table S4. Full Functional Enrichment Results post-ketamine

Table S5. Complete list of CpG sites included in MDMA BEWAS analysis with estimate and p-value results

Table S6. List of gene-level results post-MDMA

Table S7. List of genes associated with neurotransmission/neuroplasticity post-MDMA

Table S8. Full Functional Enrichment Results post-MDMA

Data file S1. DNA methylation of Ketamine cohort before and after treatment Data file S2. DNA methylation of MDMA cohort before and after treatment

## Funding

Modern Spirit; 501c3 non-profit

Board of Medicine; 501c3 non-profit

## Author contributions

Conceptualization: MGS, CRL

Methodology: MGS, CRL

Visualization: MGS, SEM, CRL

Funding acquisition: JT, RS, BY-K

Project administration: BRC, BY-K, RS, VBD, CRL, DR

Supervision: SEM, CRL

Writing – original draft: MGS, SEM, CRL

Writing – review & editing: RS, VBD, BRC, JT, BY-K, DR

## Data and materials availability

Data will be made available upon request from qualified individuals.

---

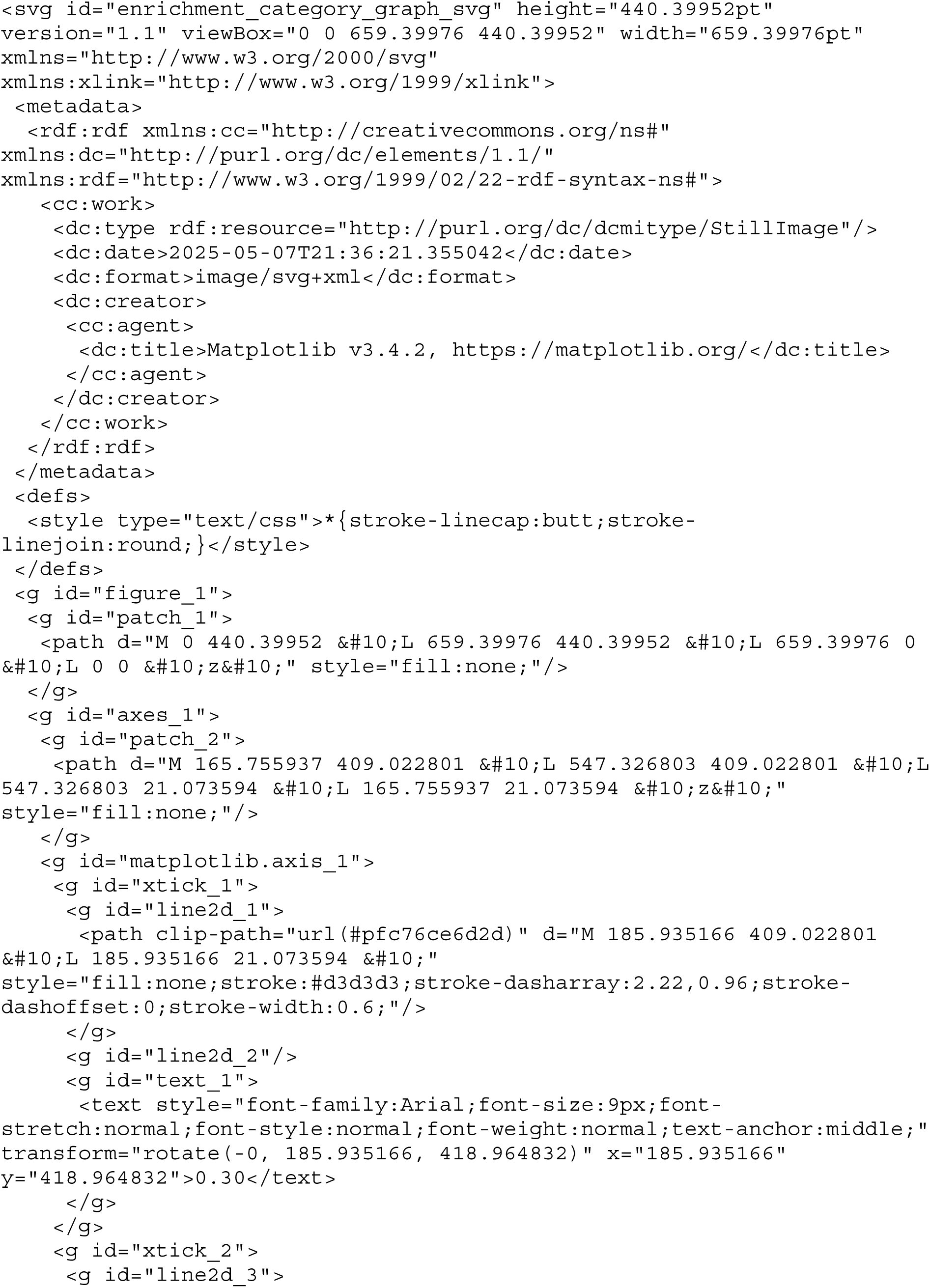

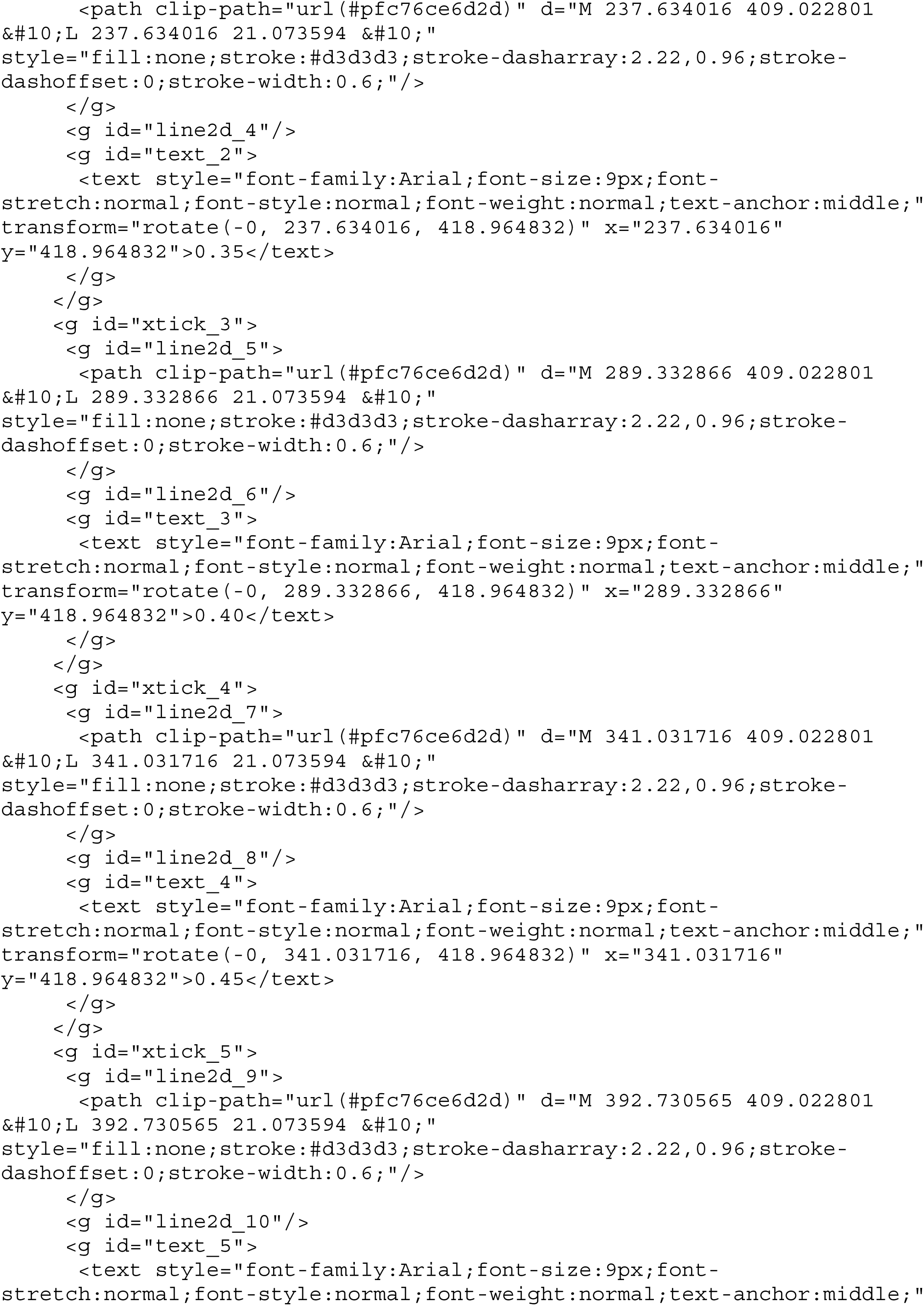

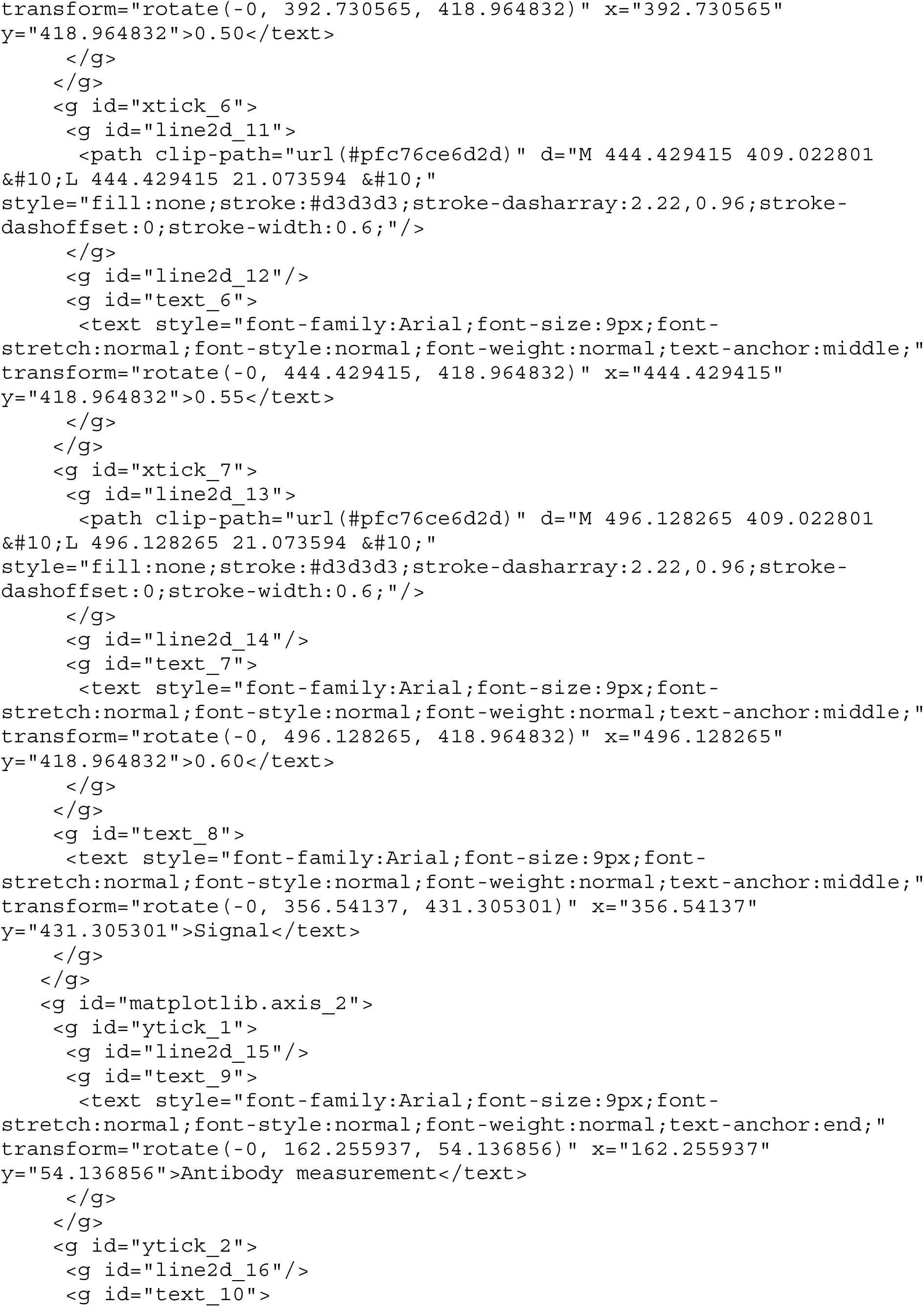

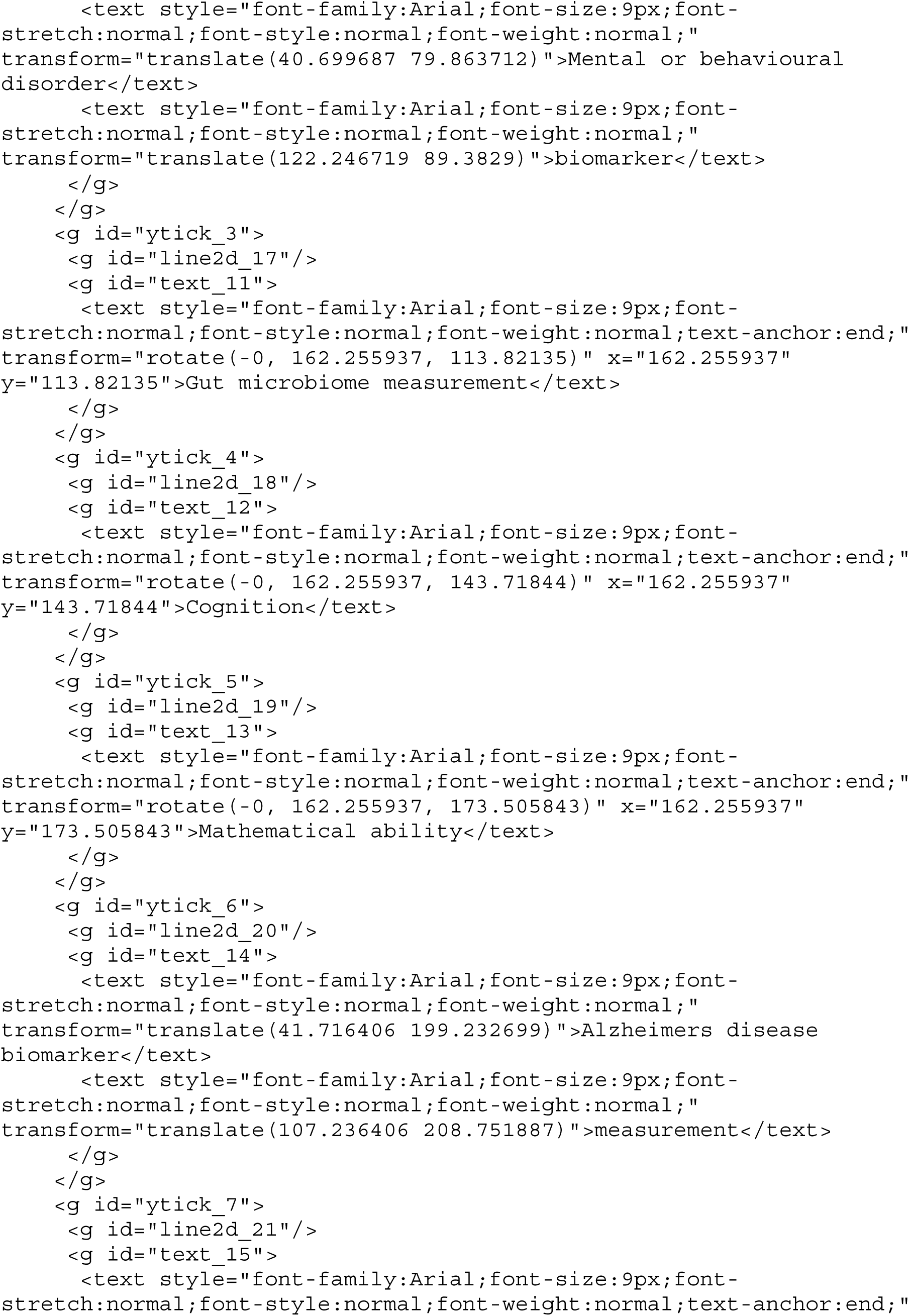

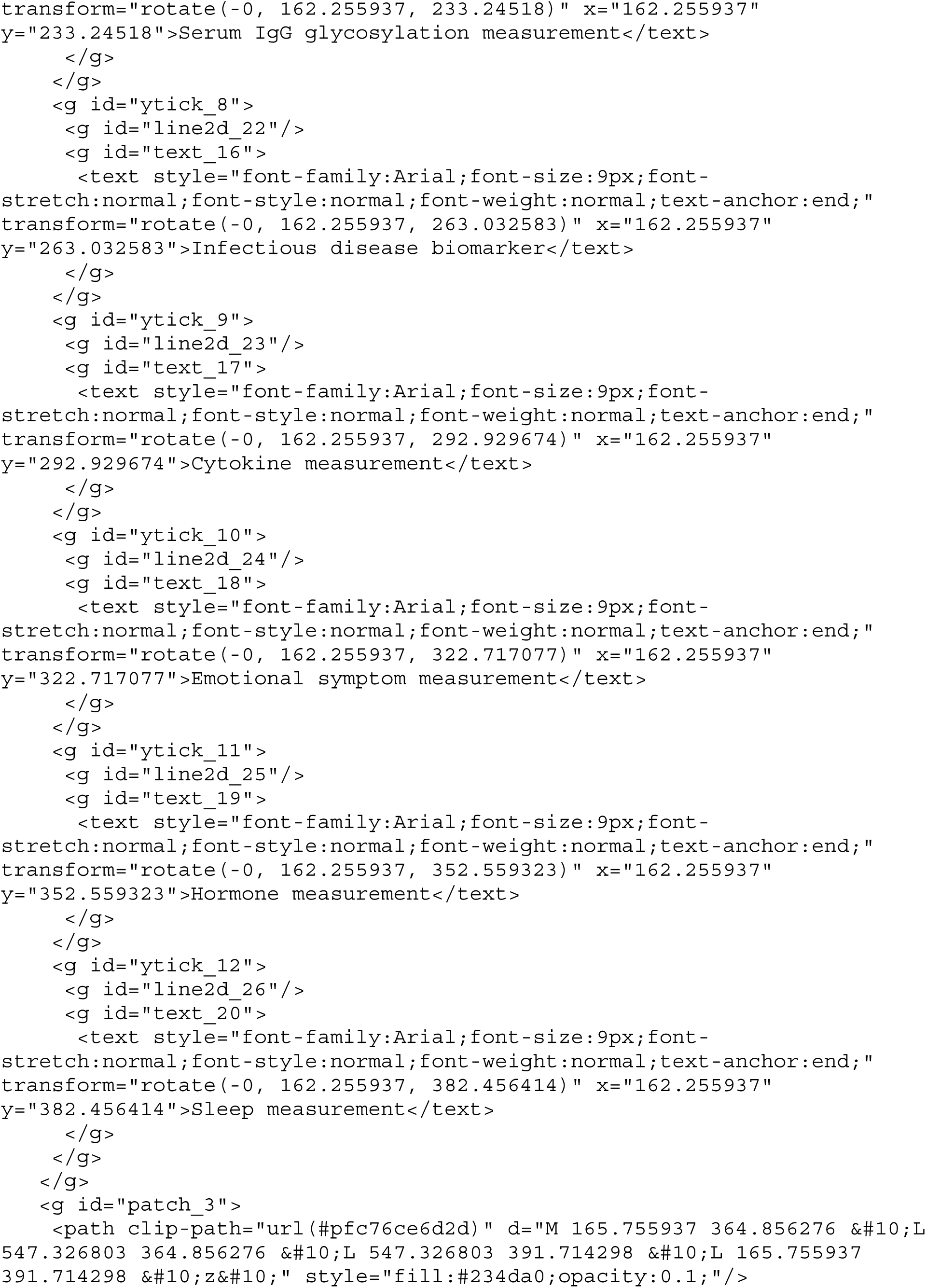

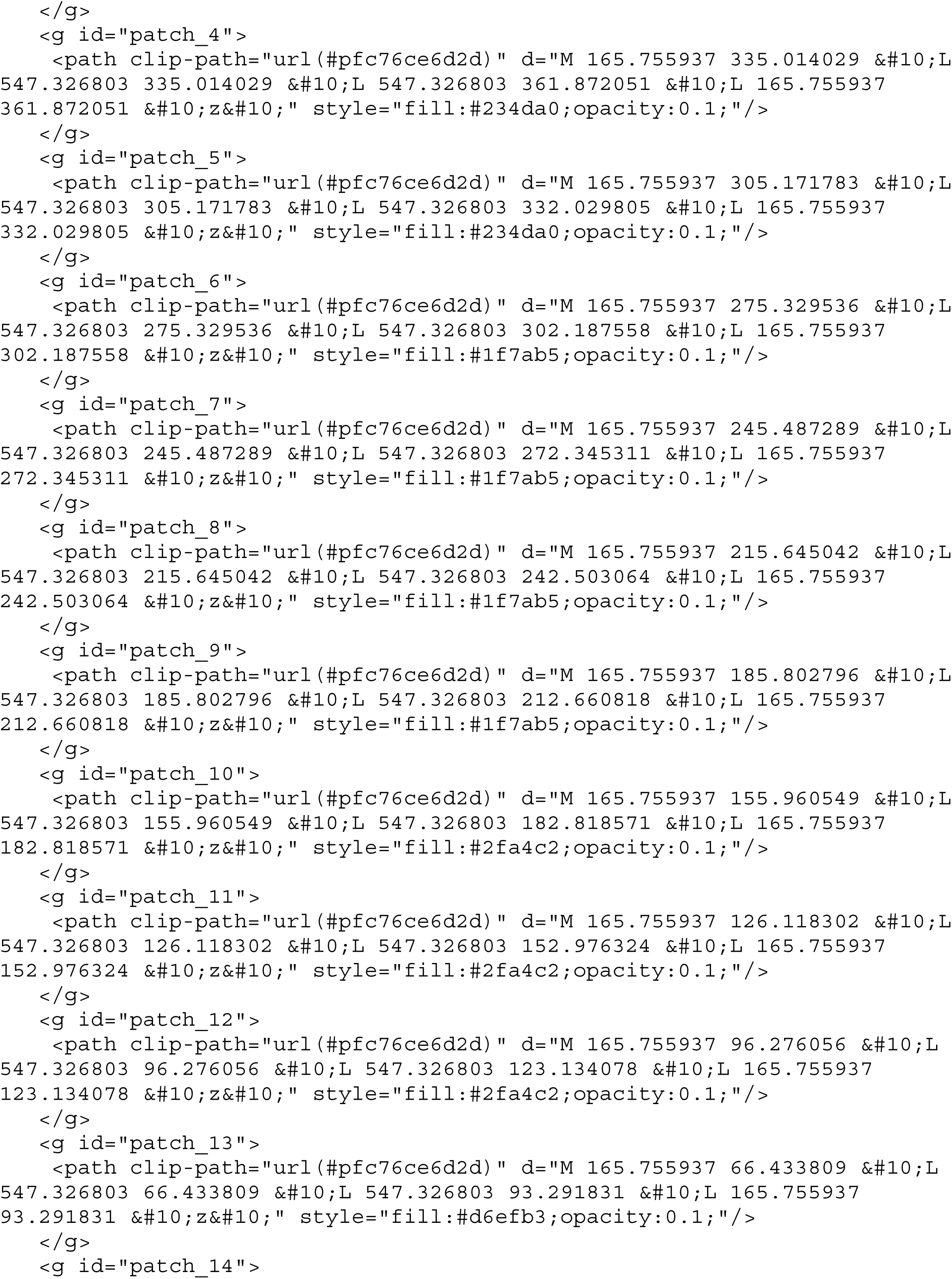

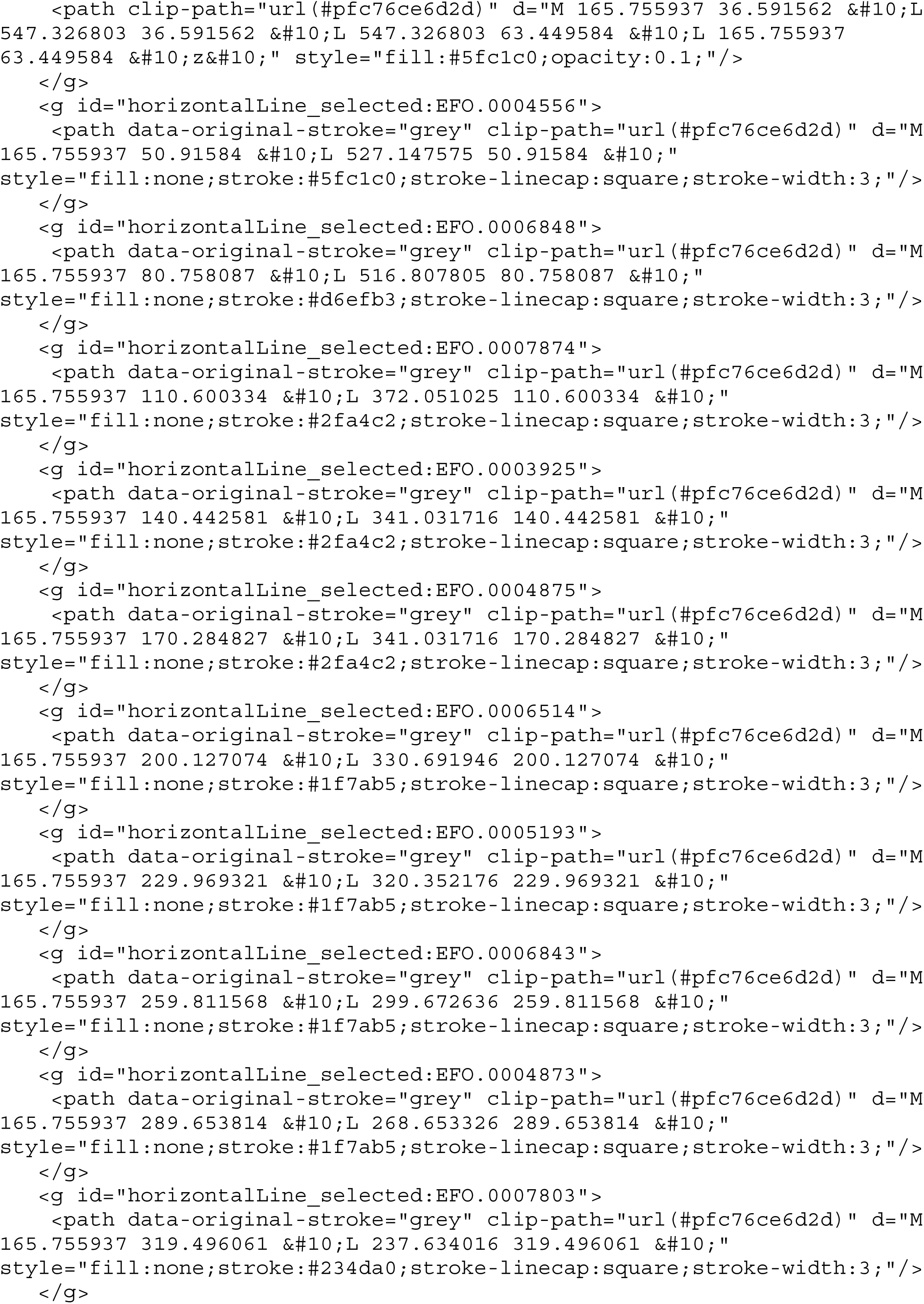

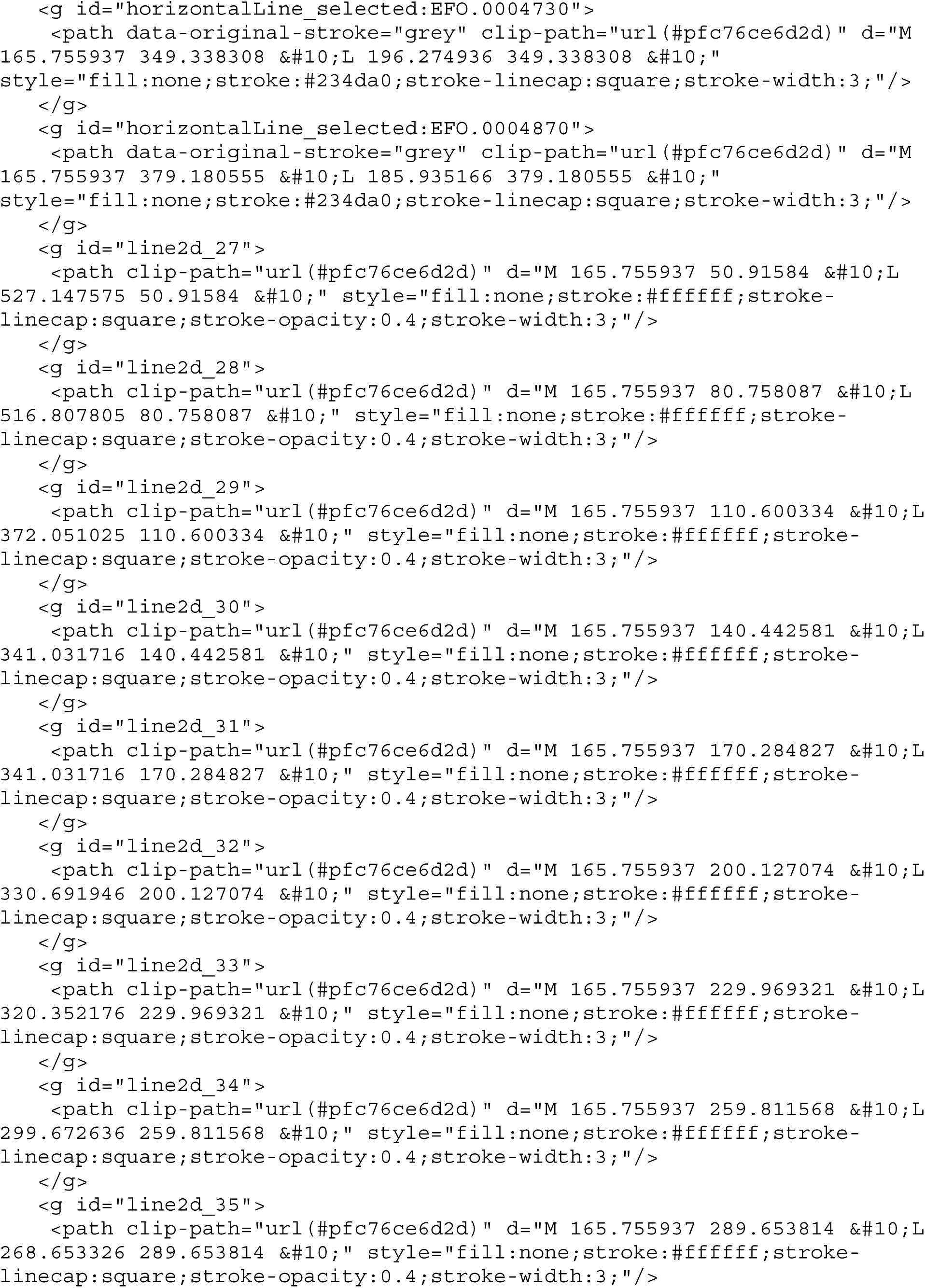

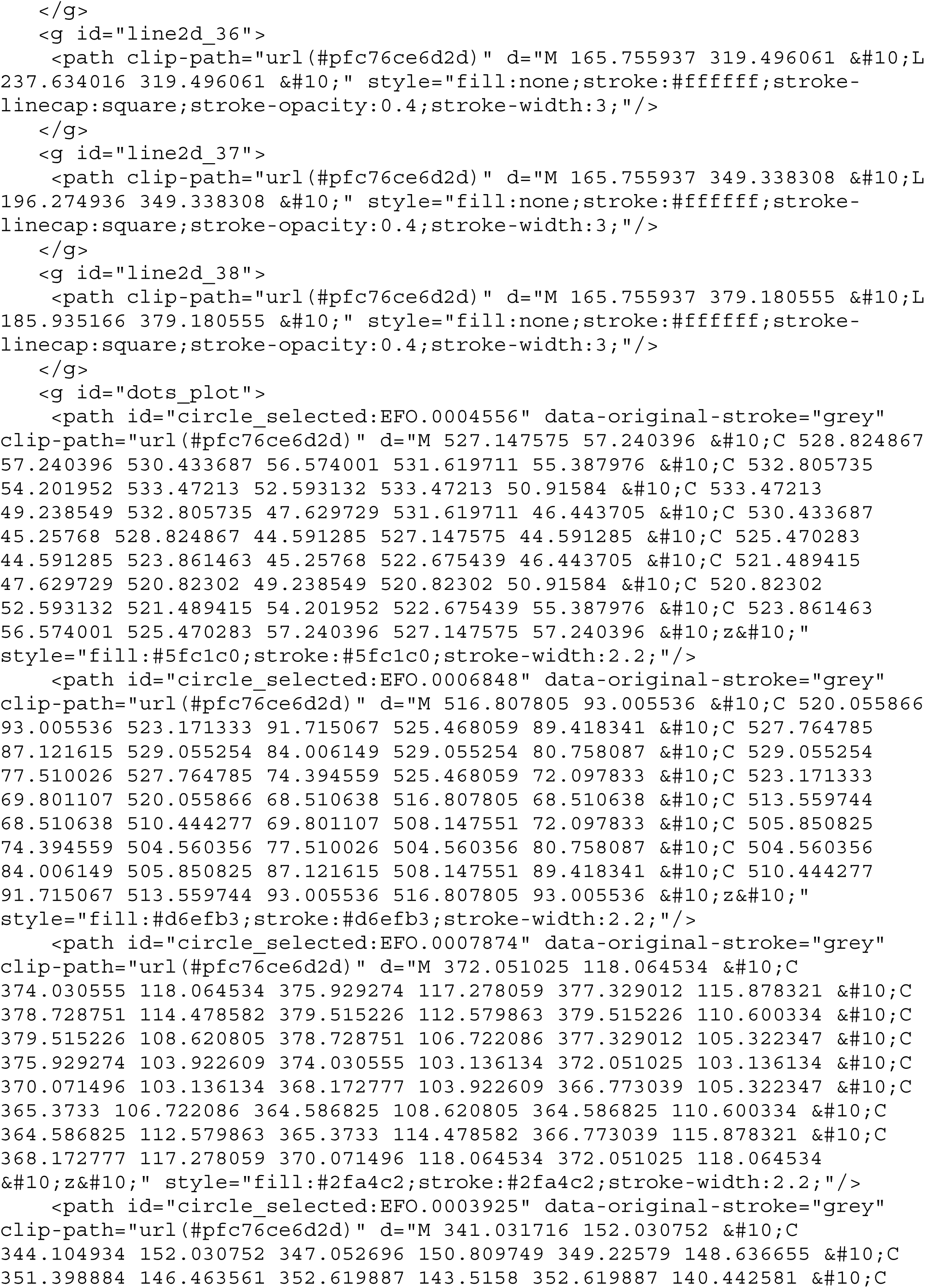

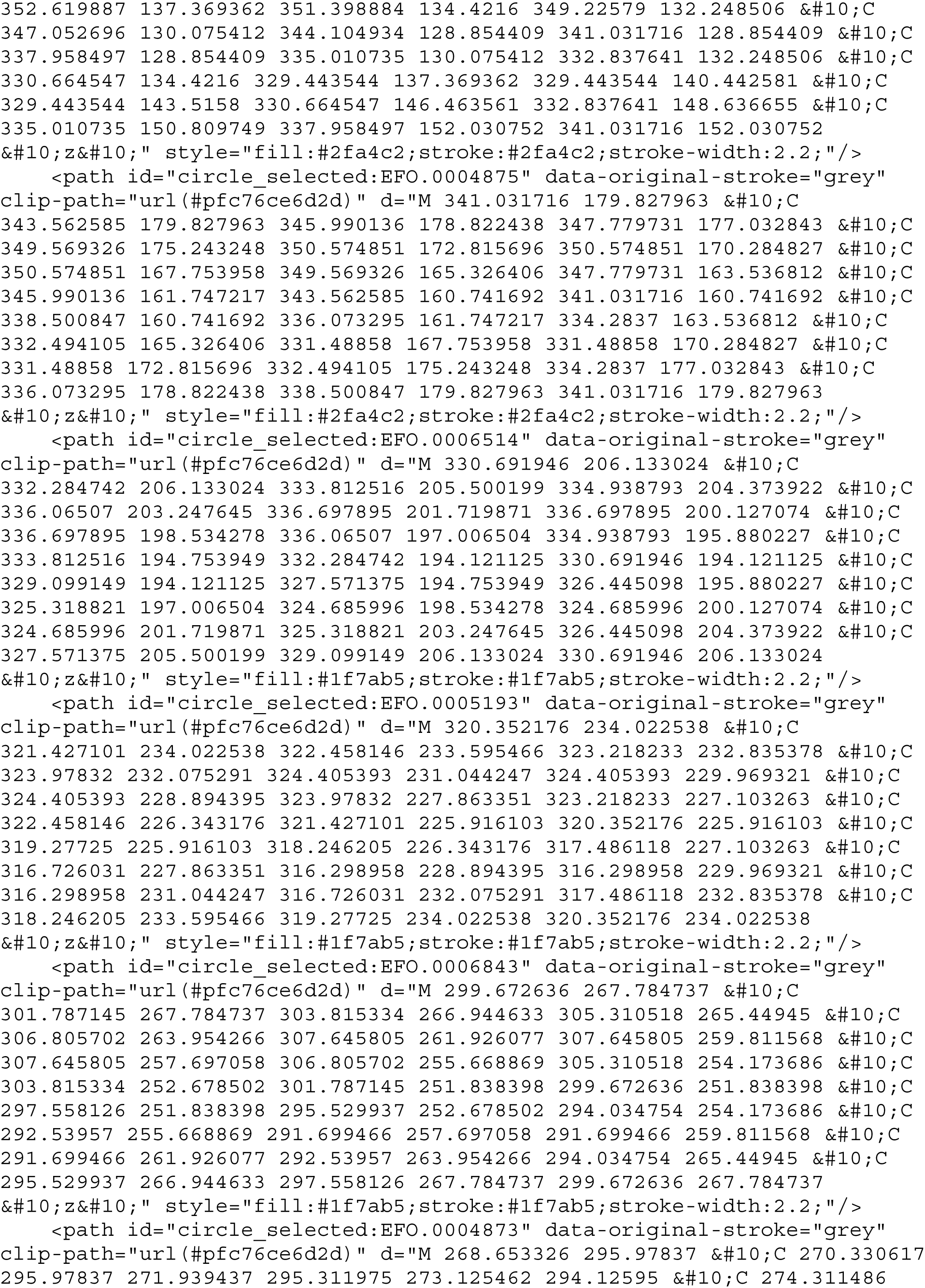

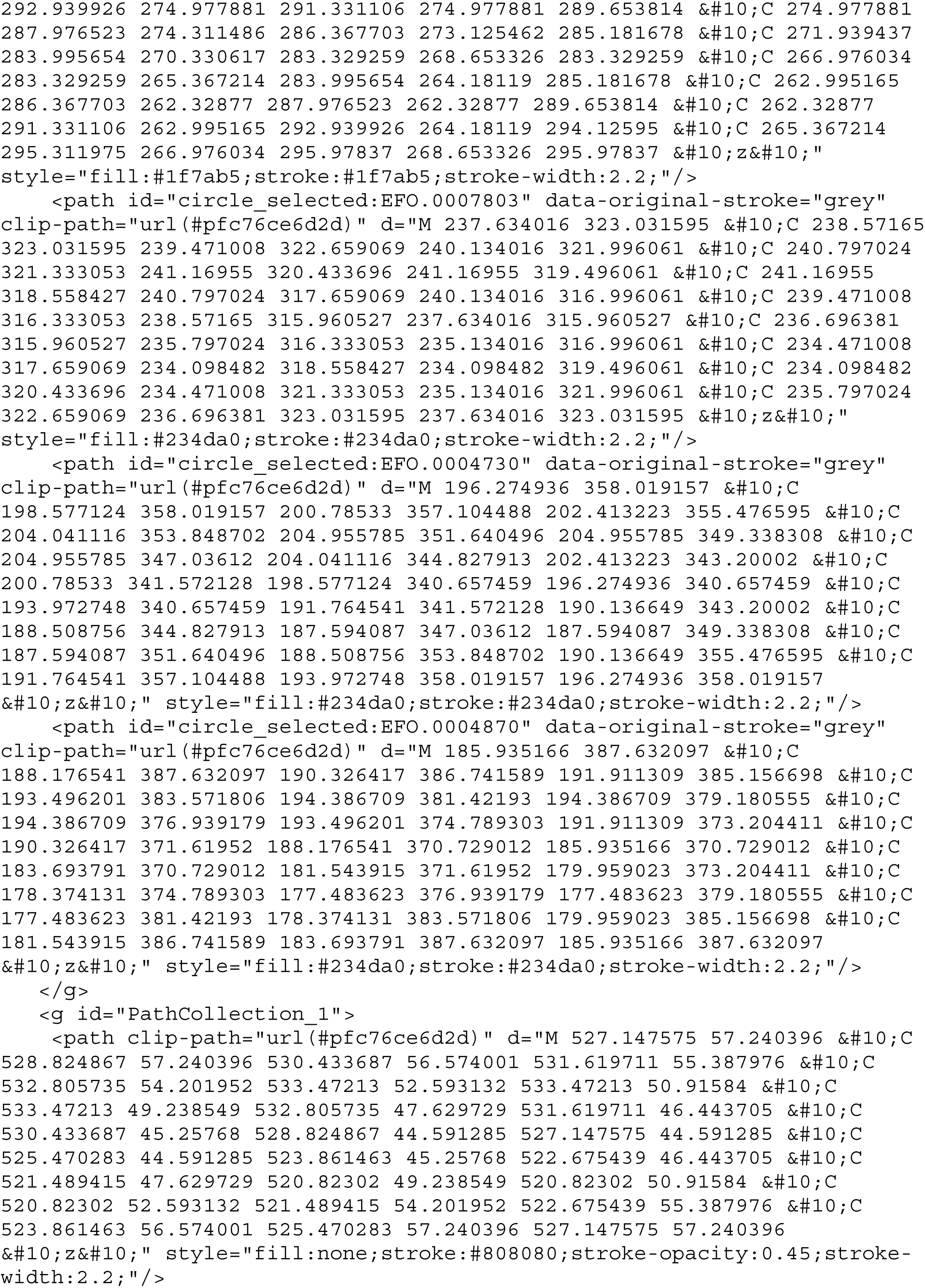

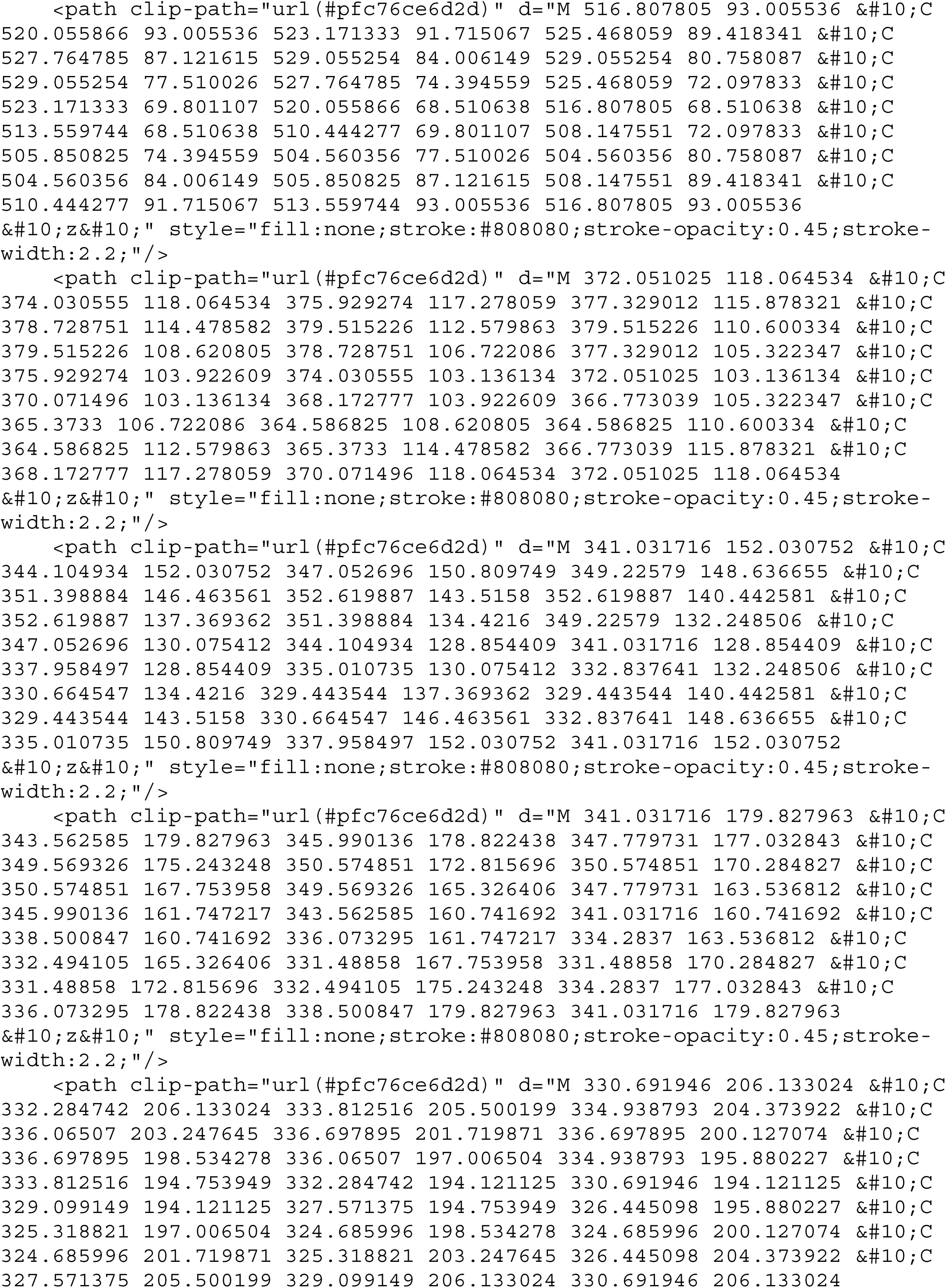

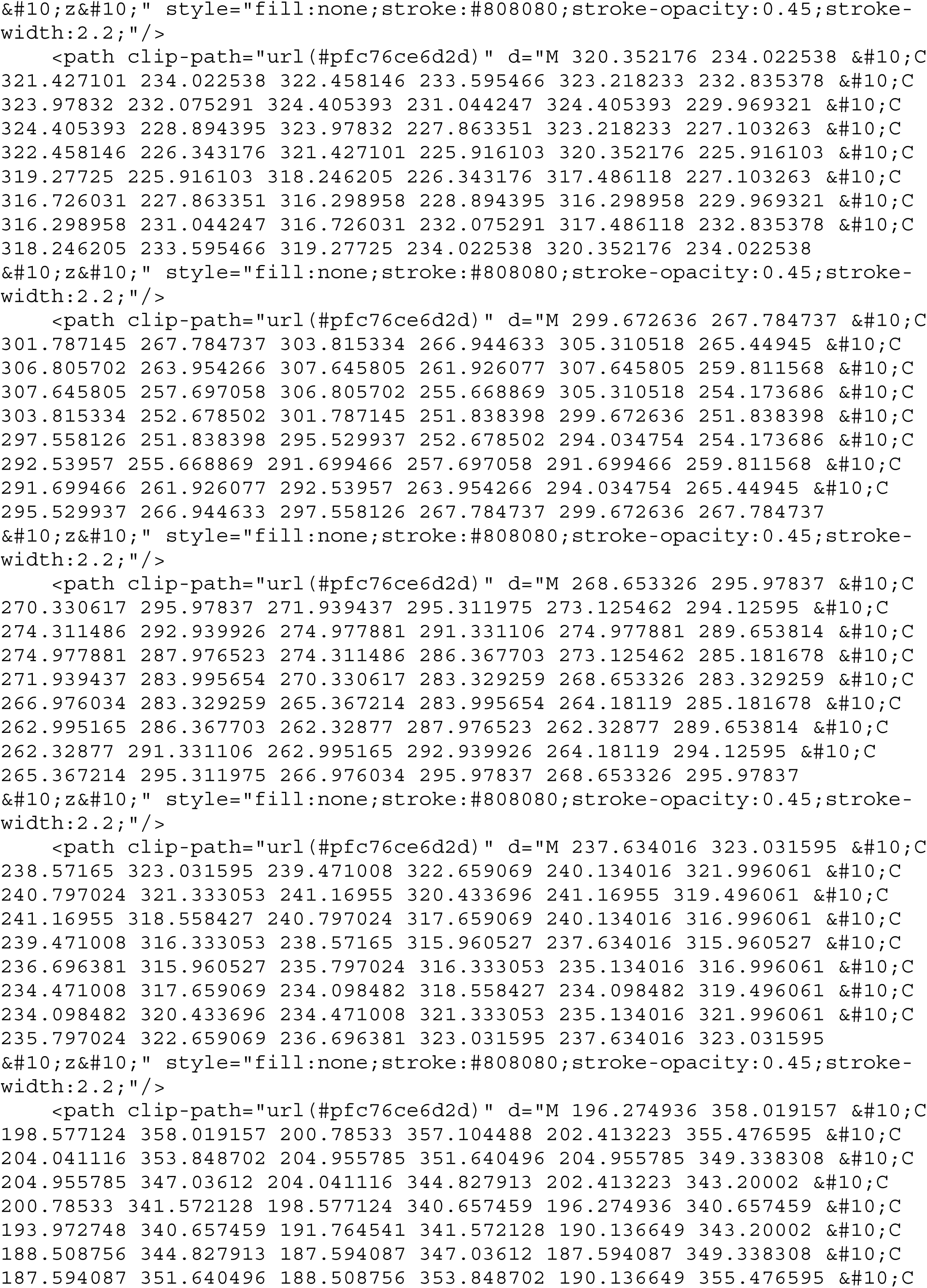

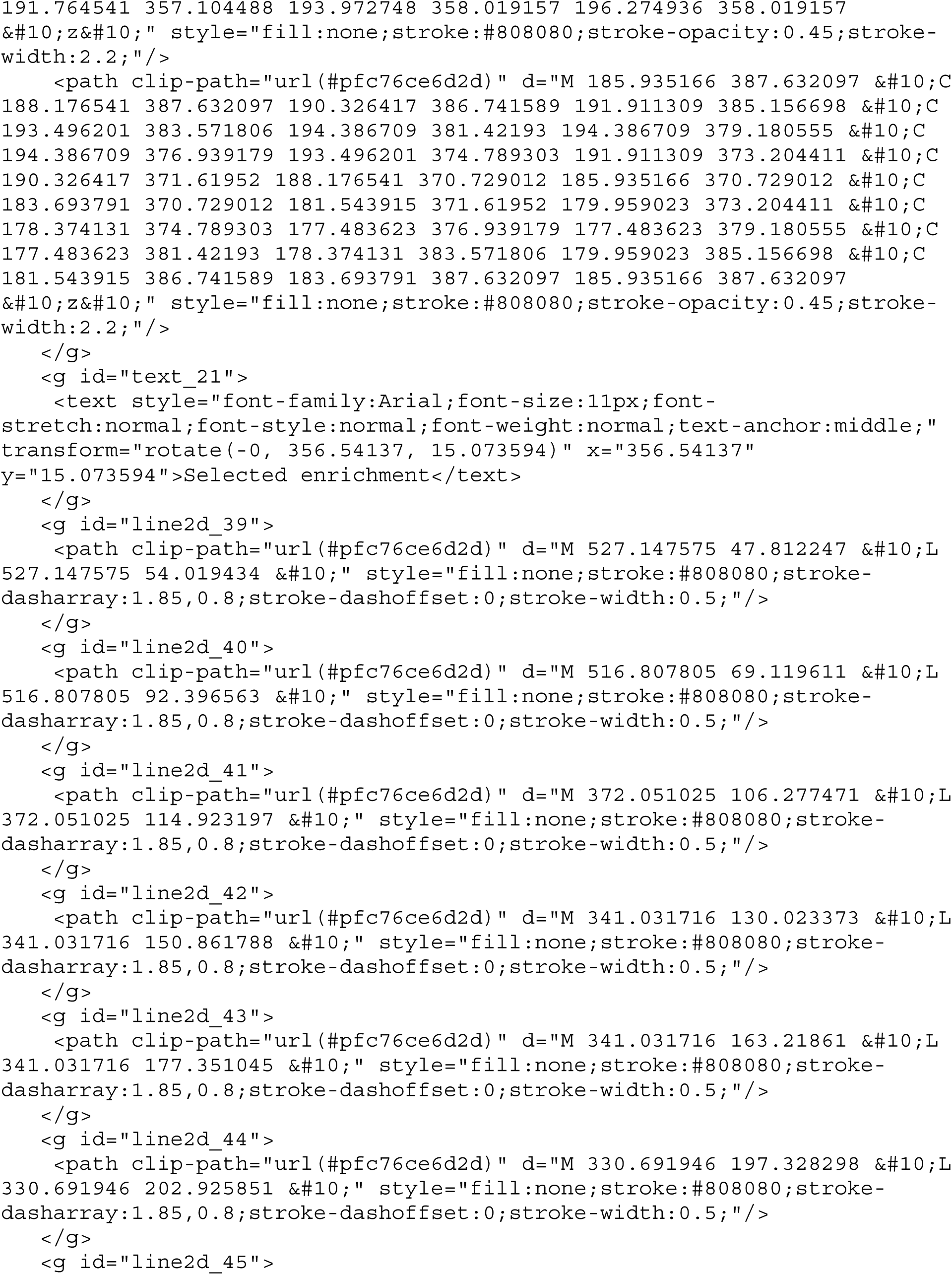

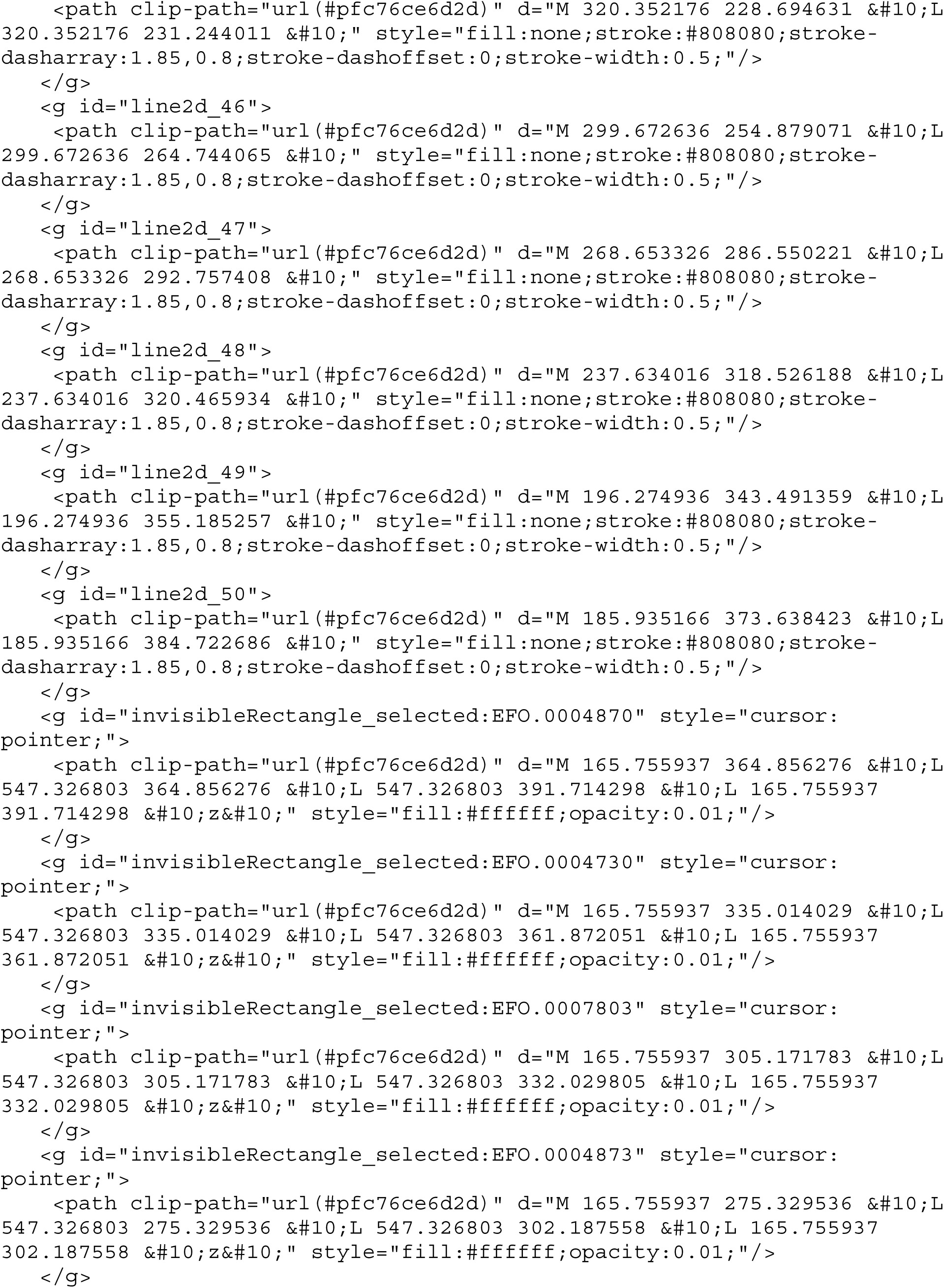

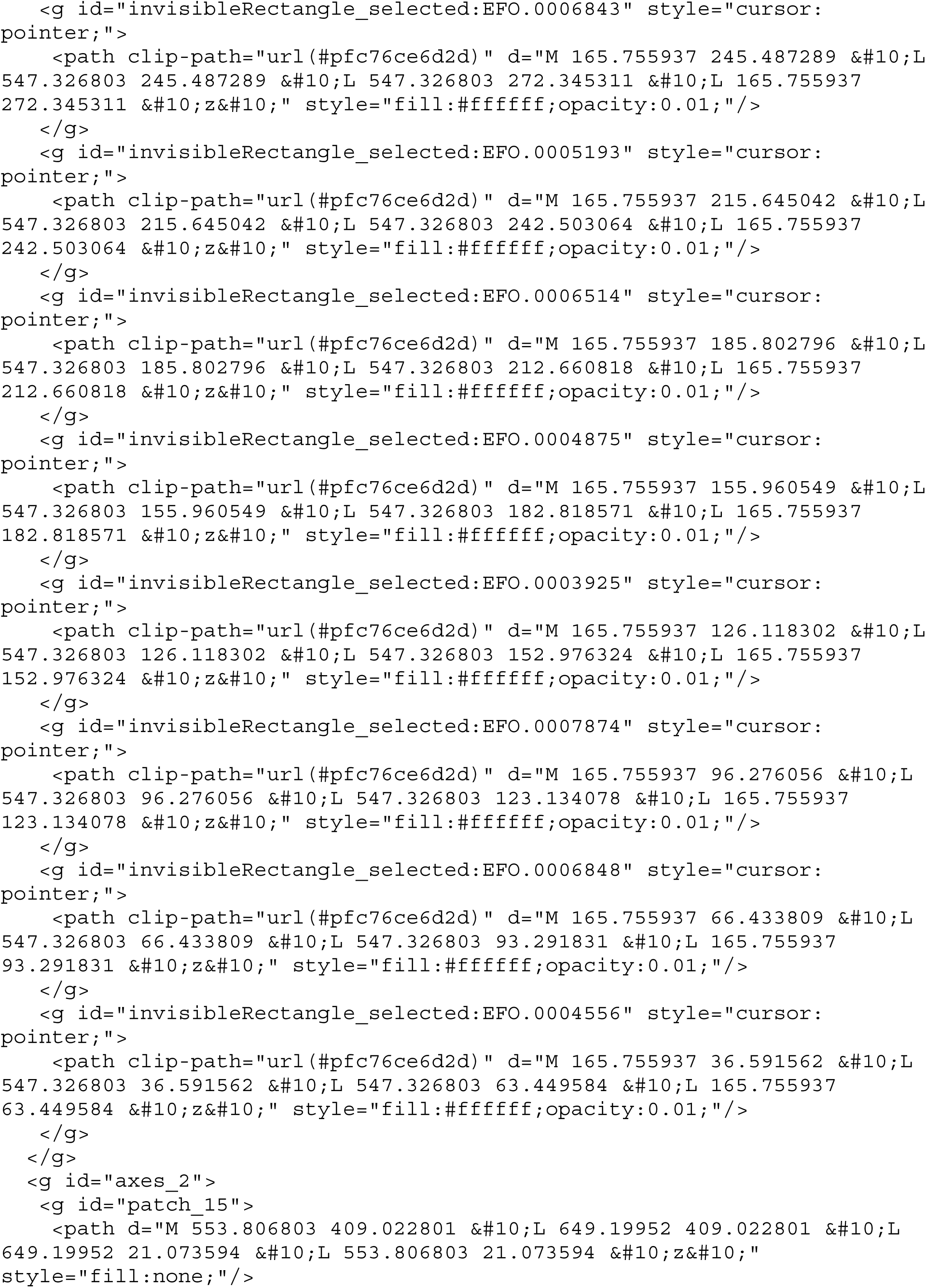

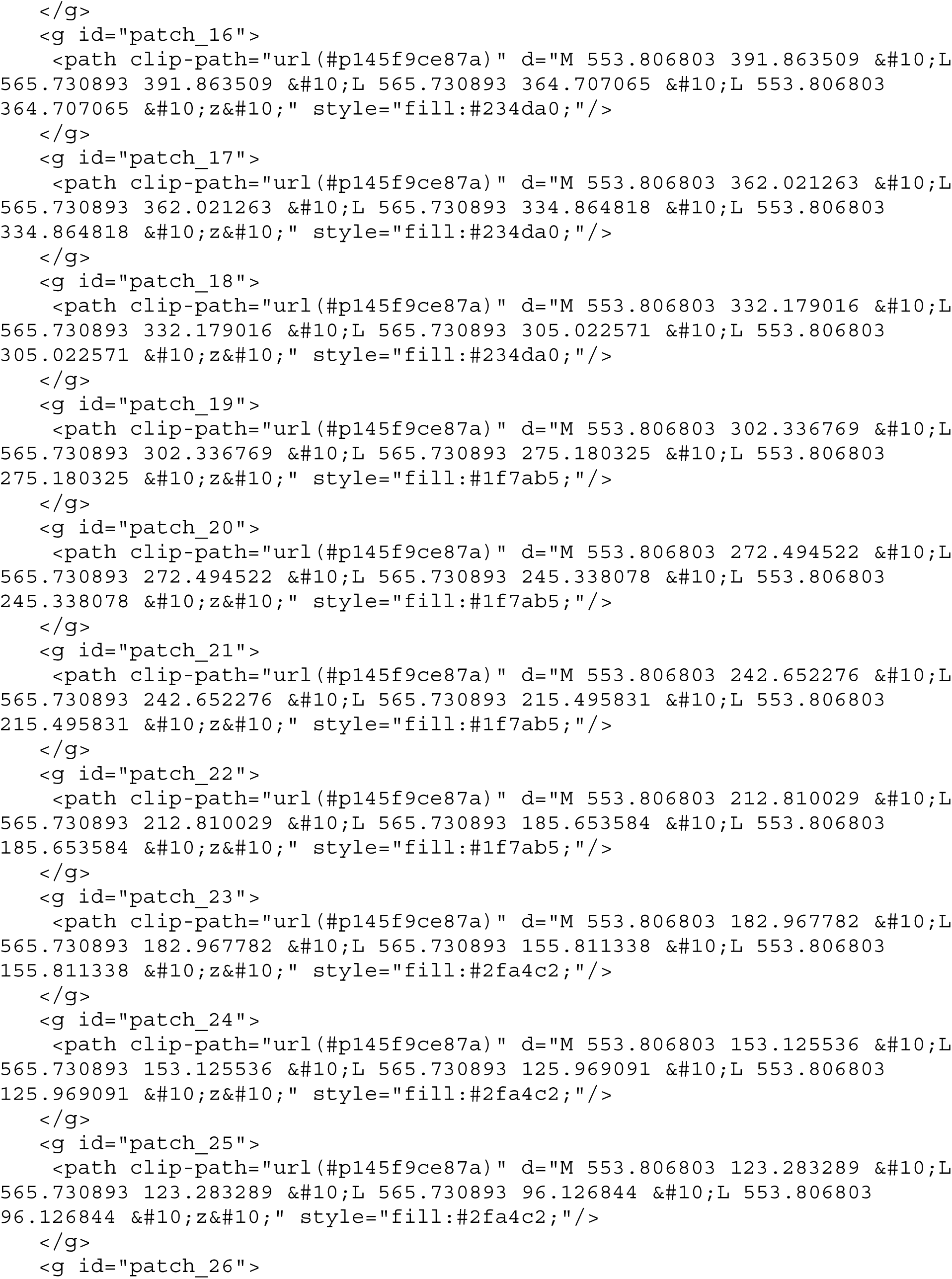

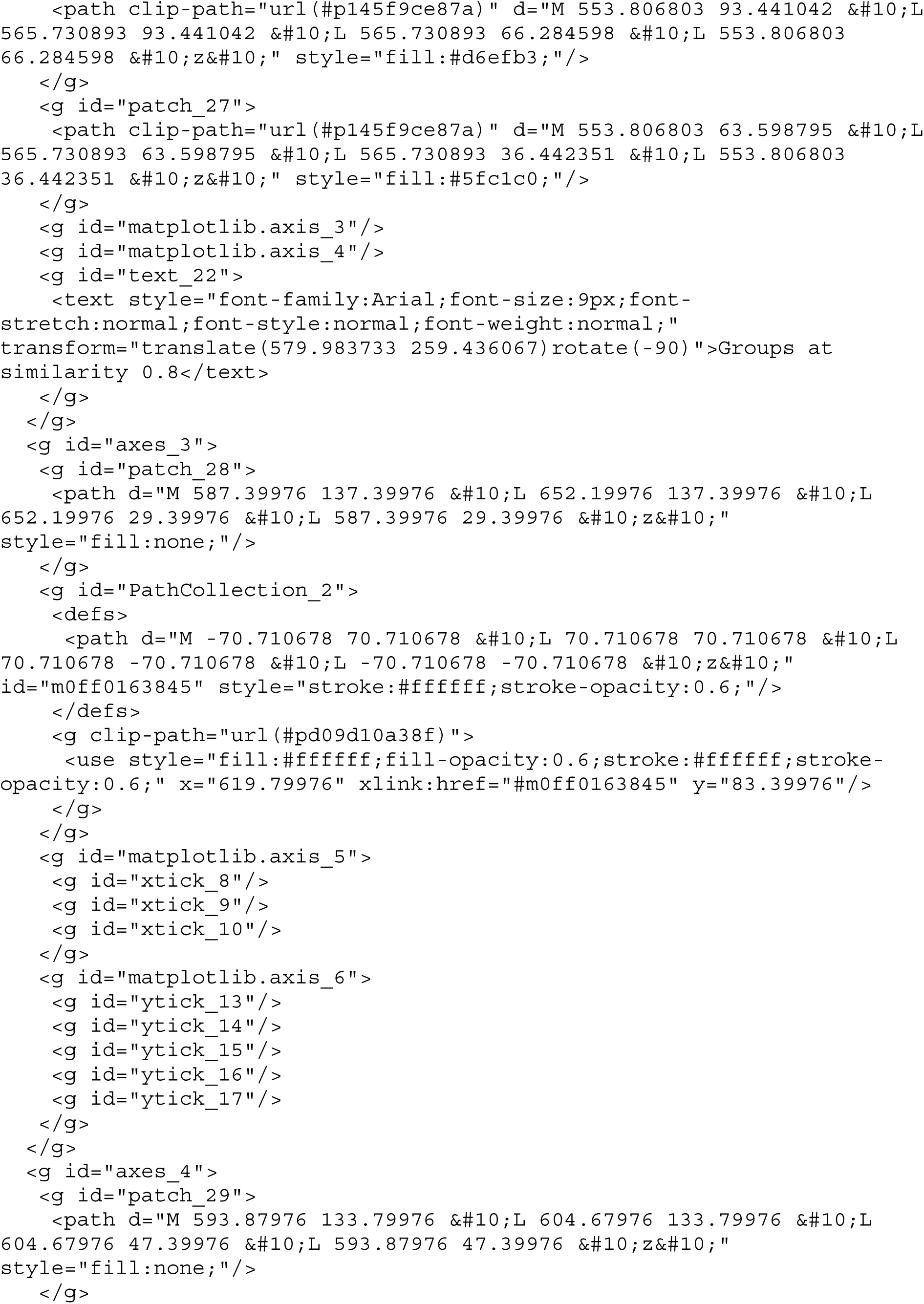

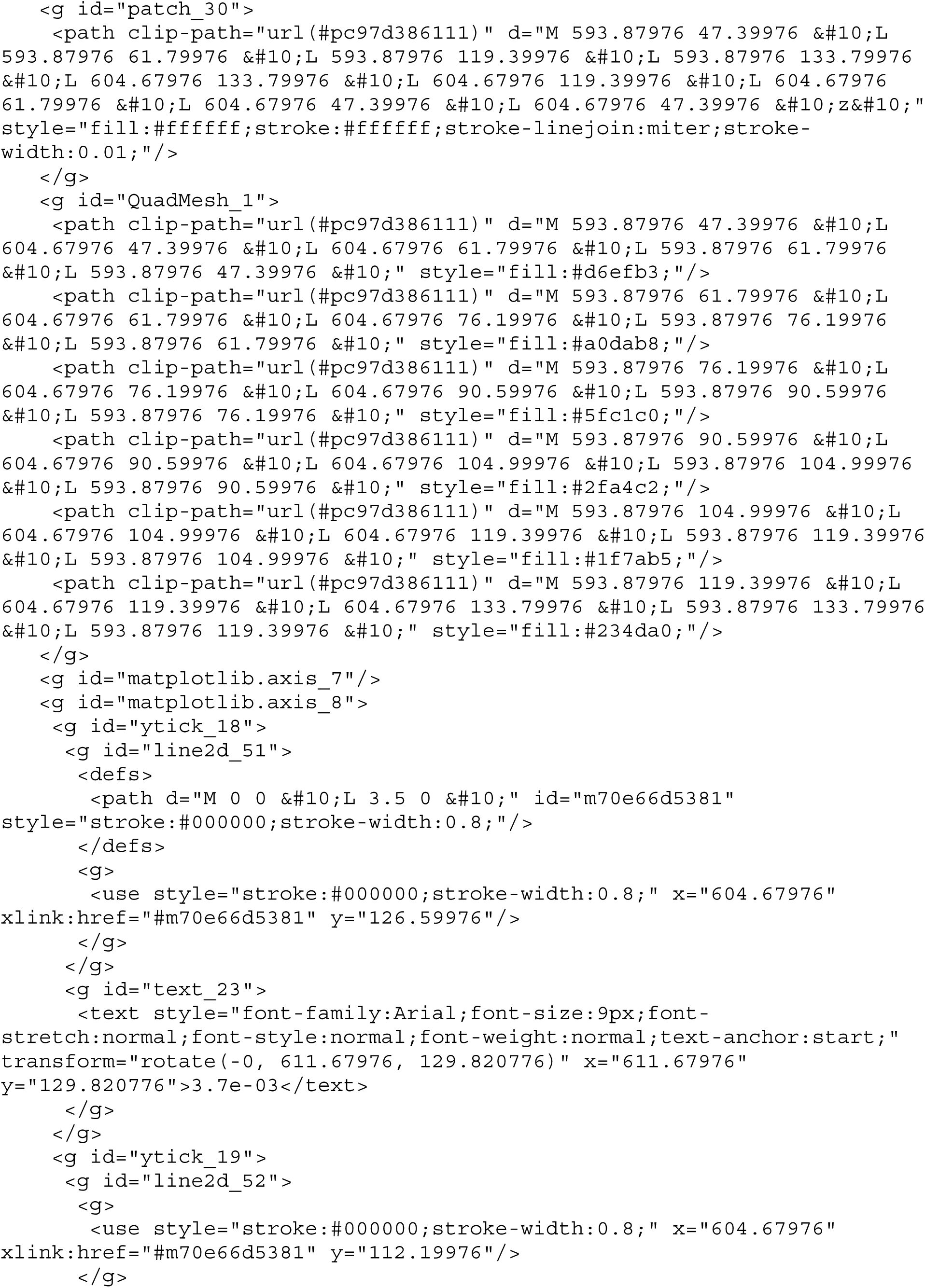

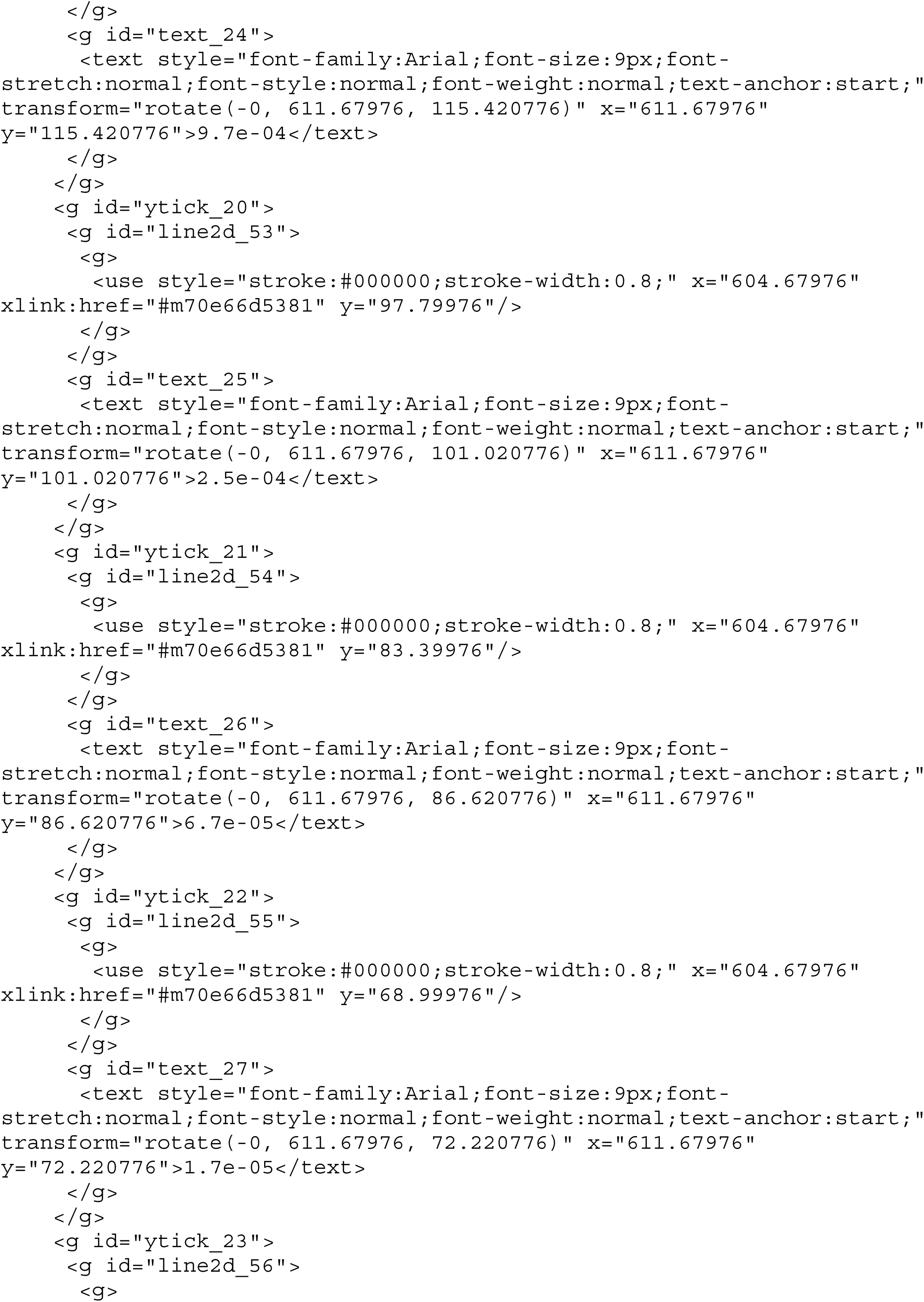

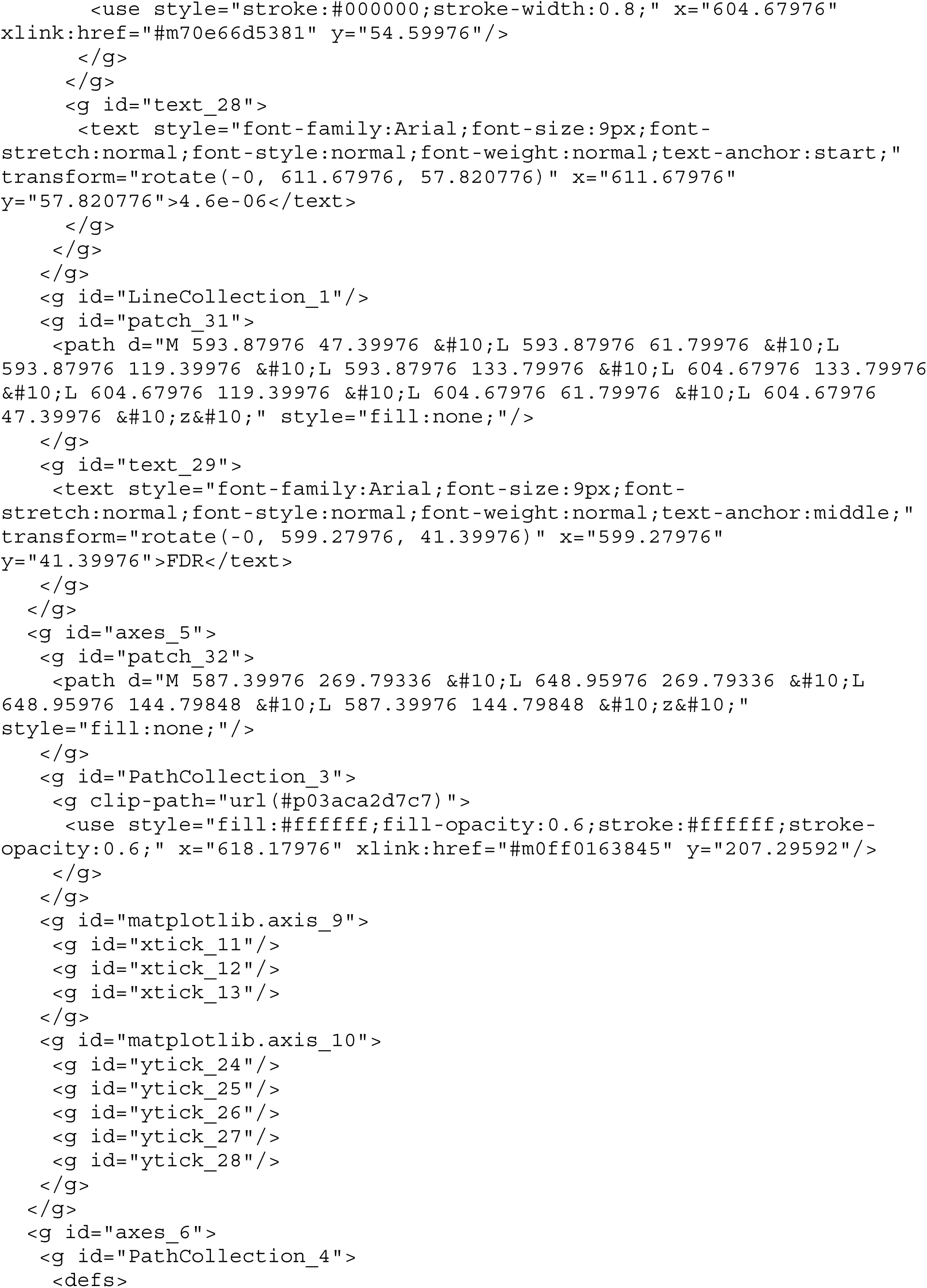

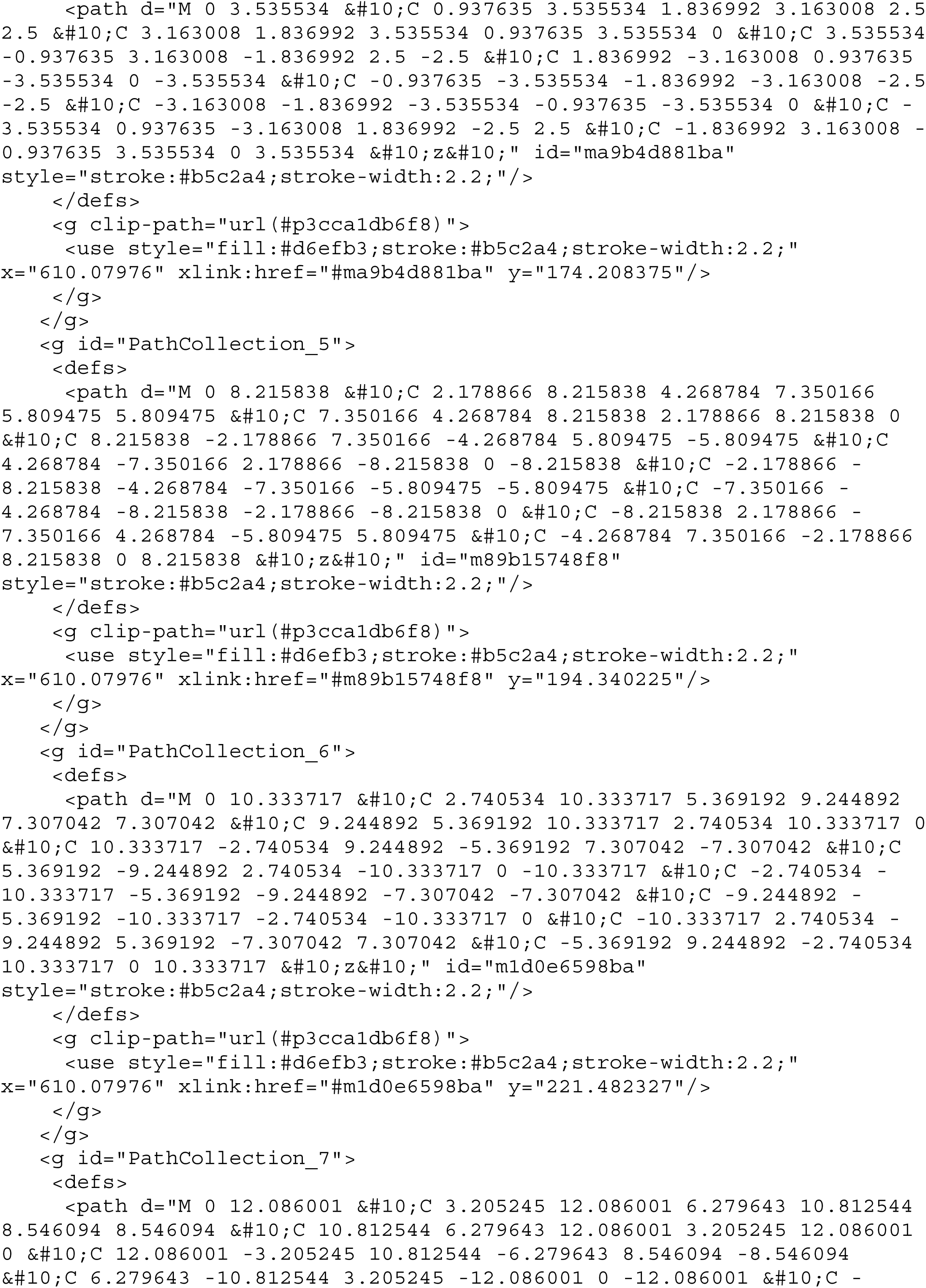

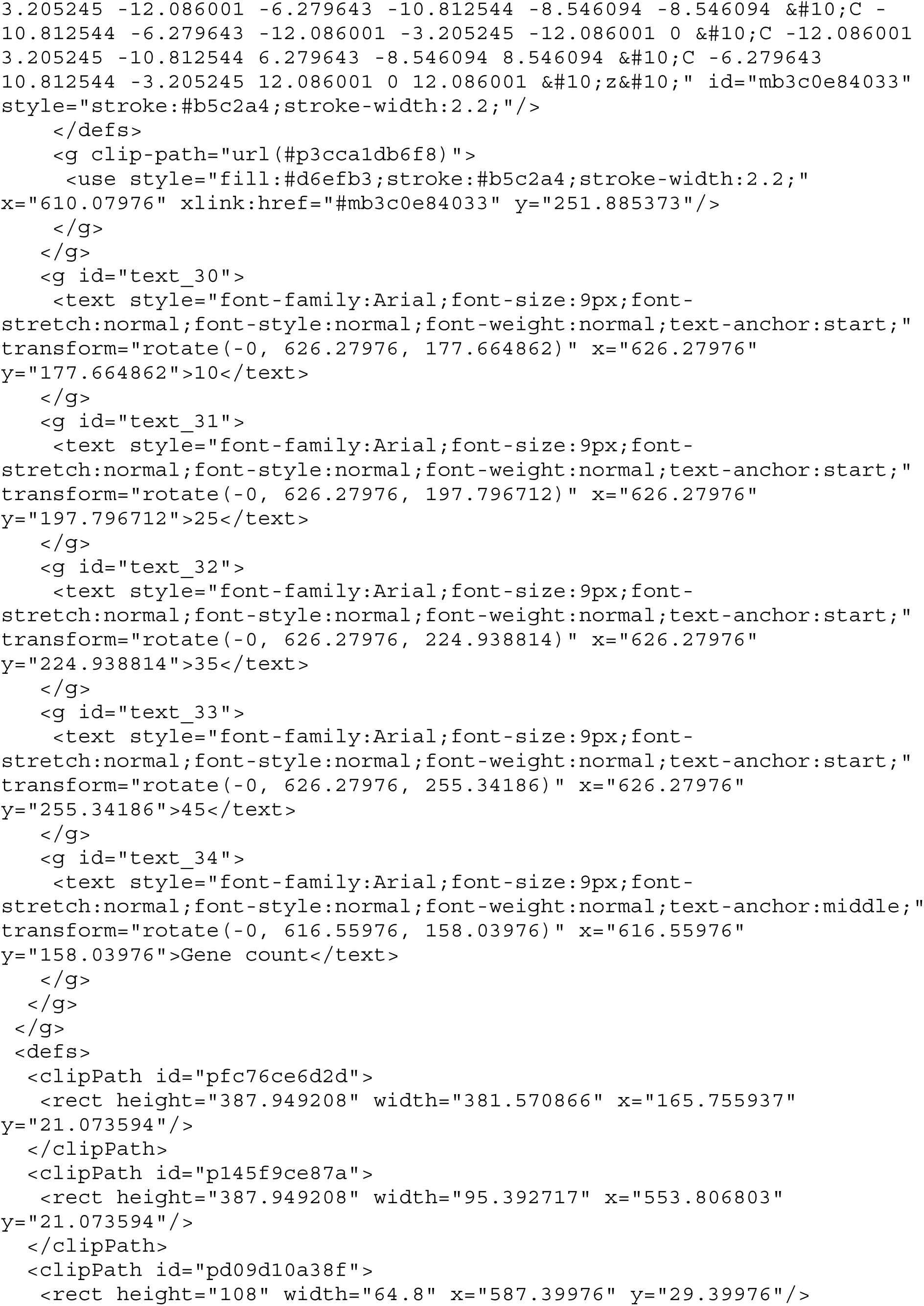

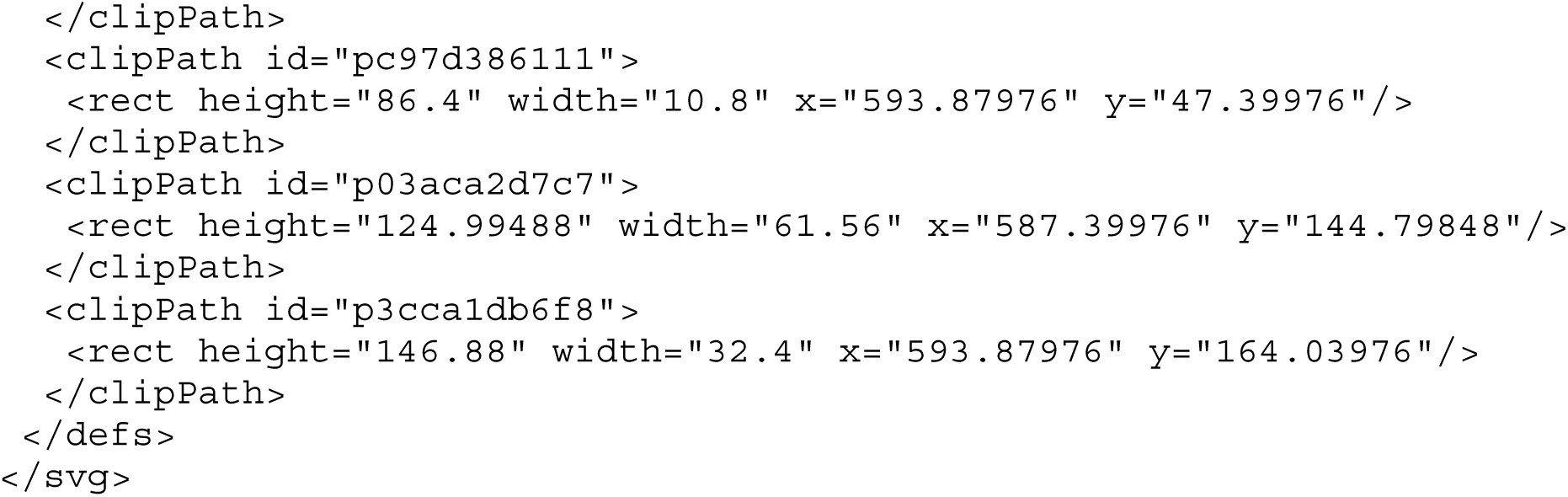

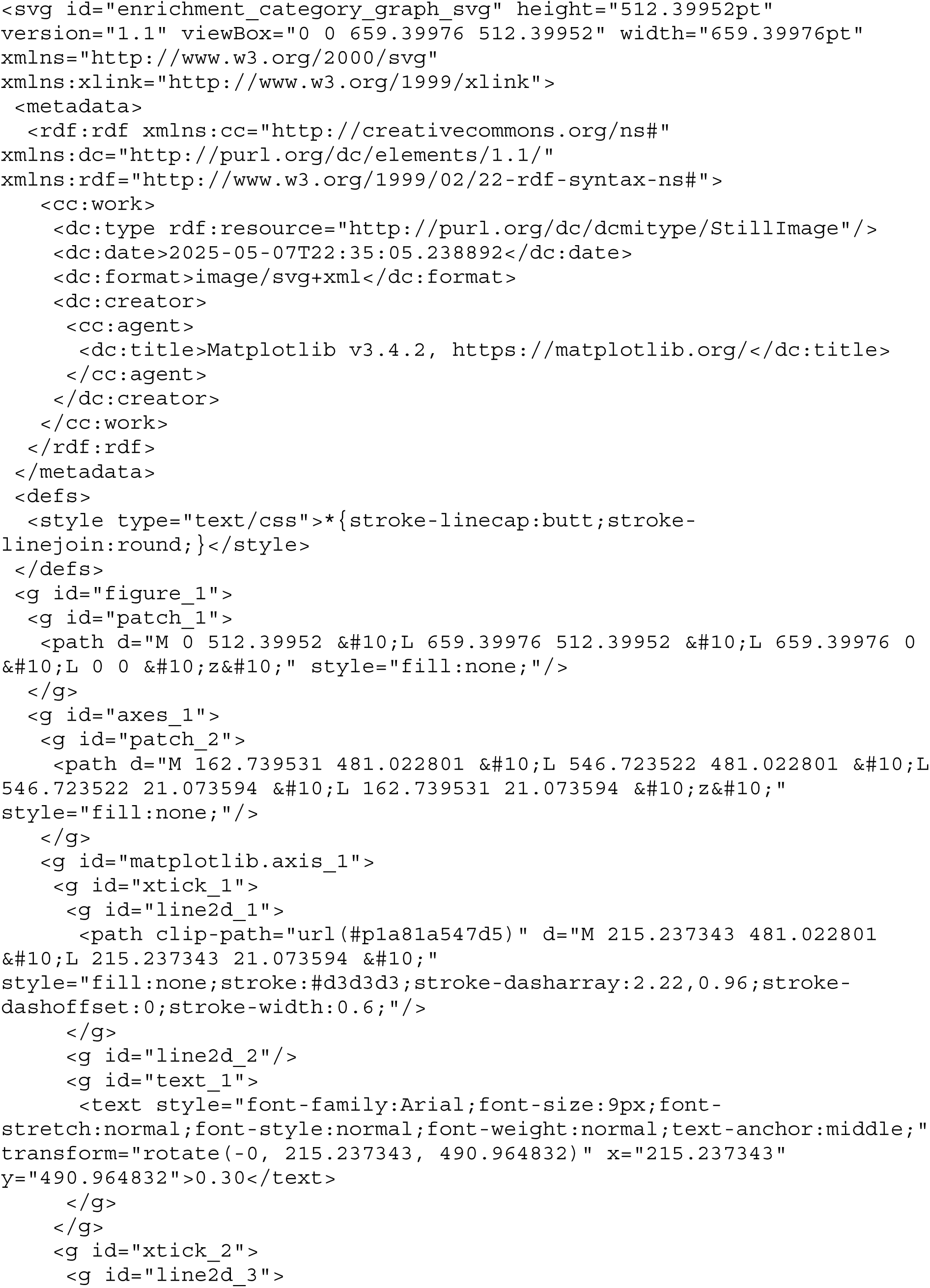

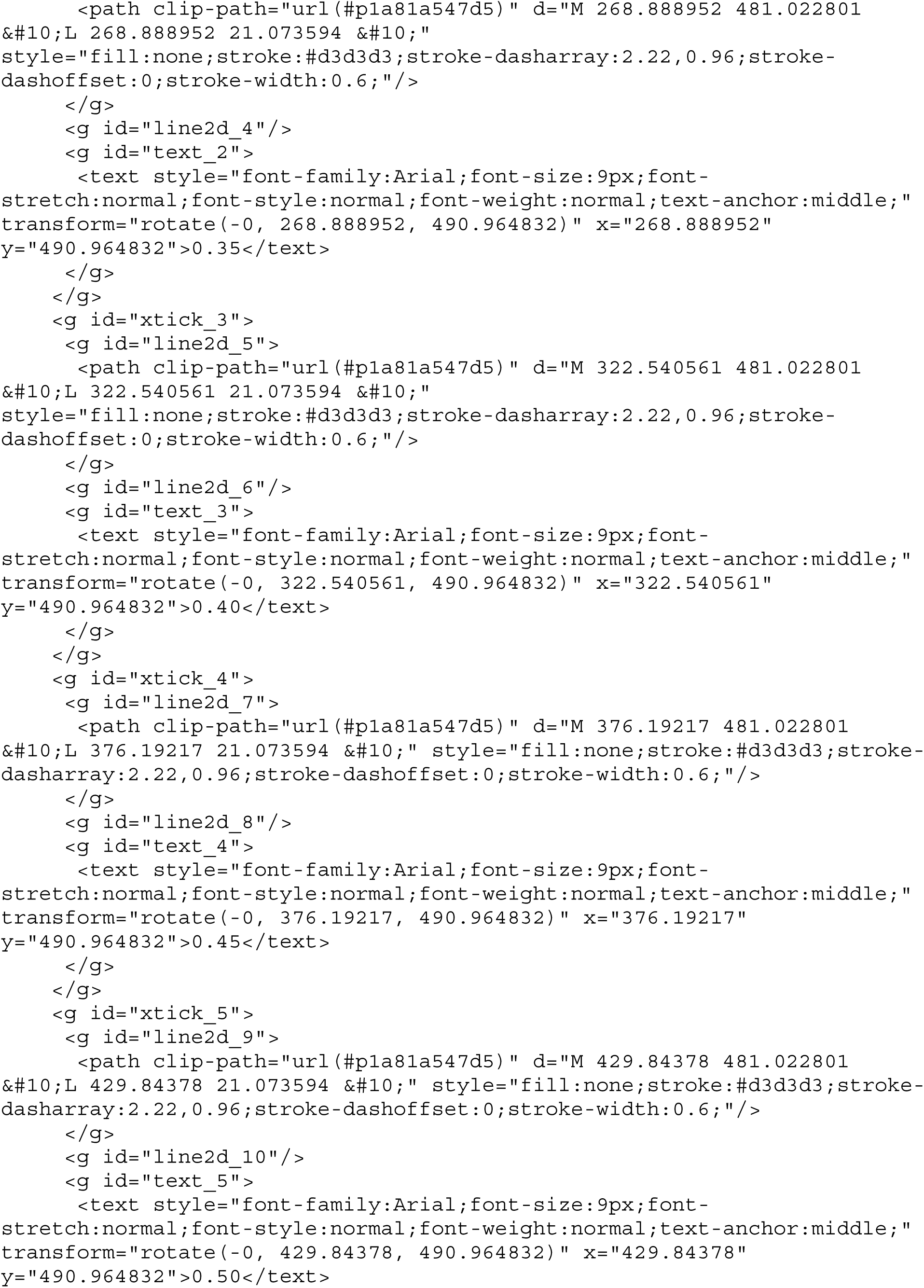

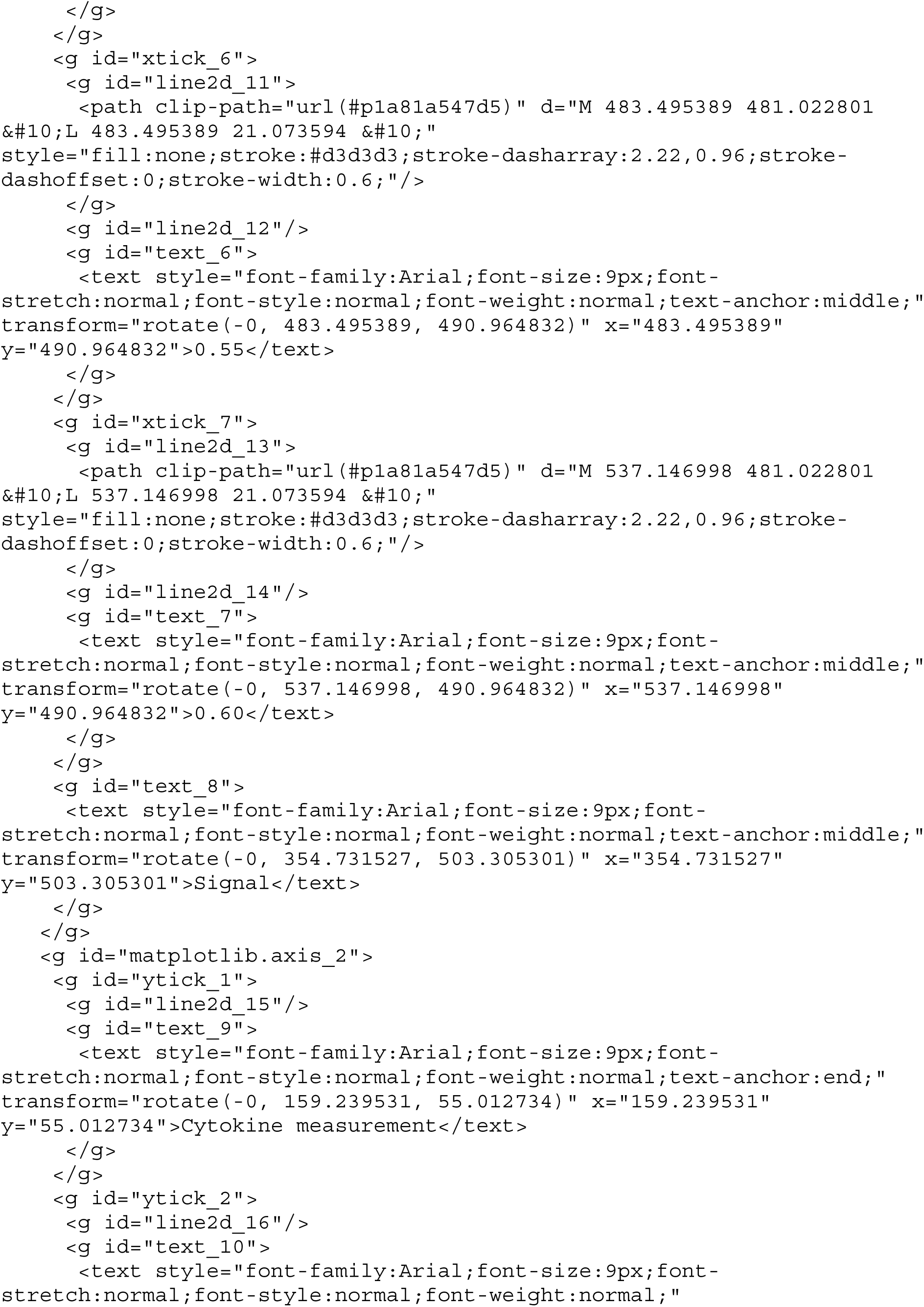

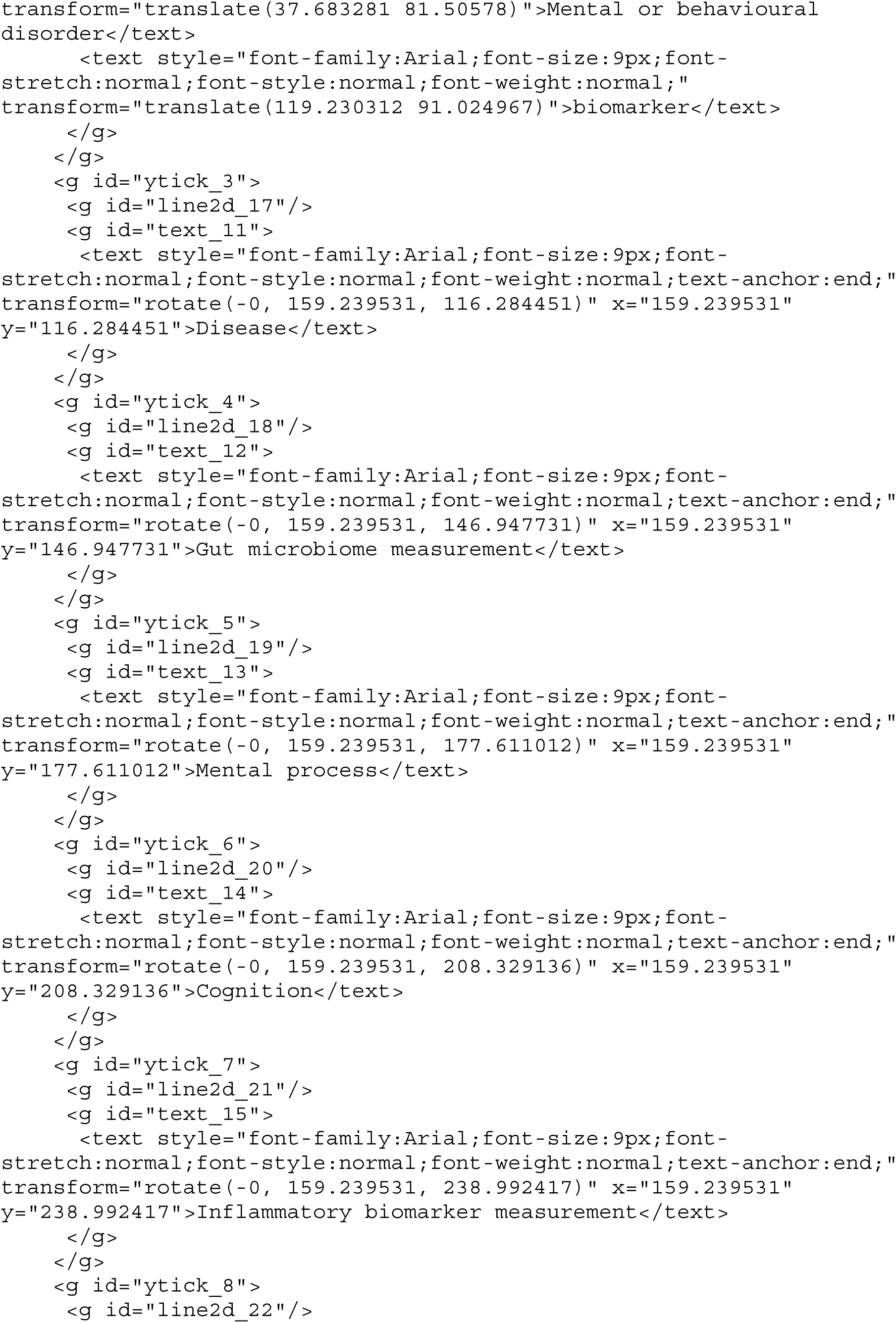

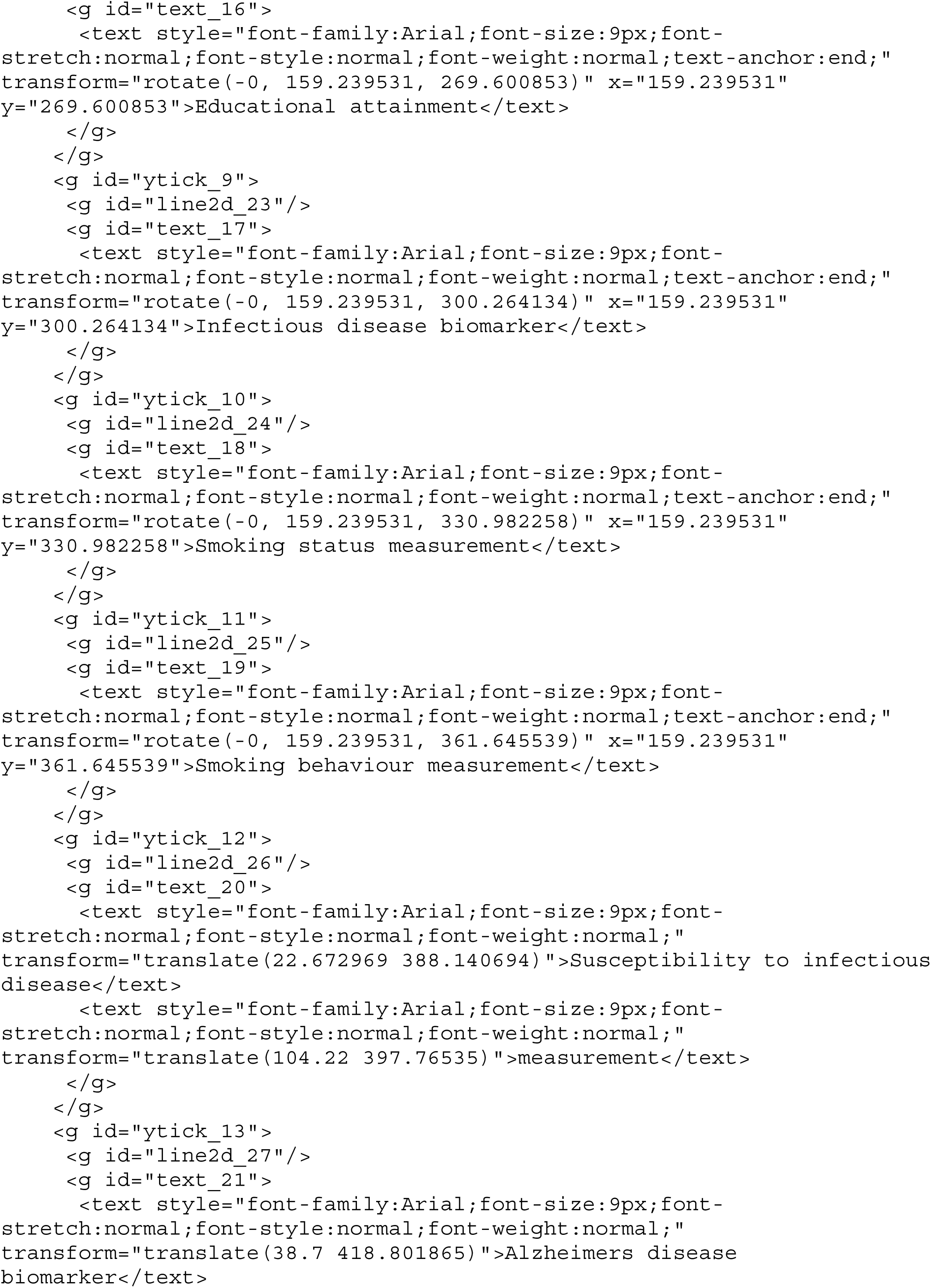

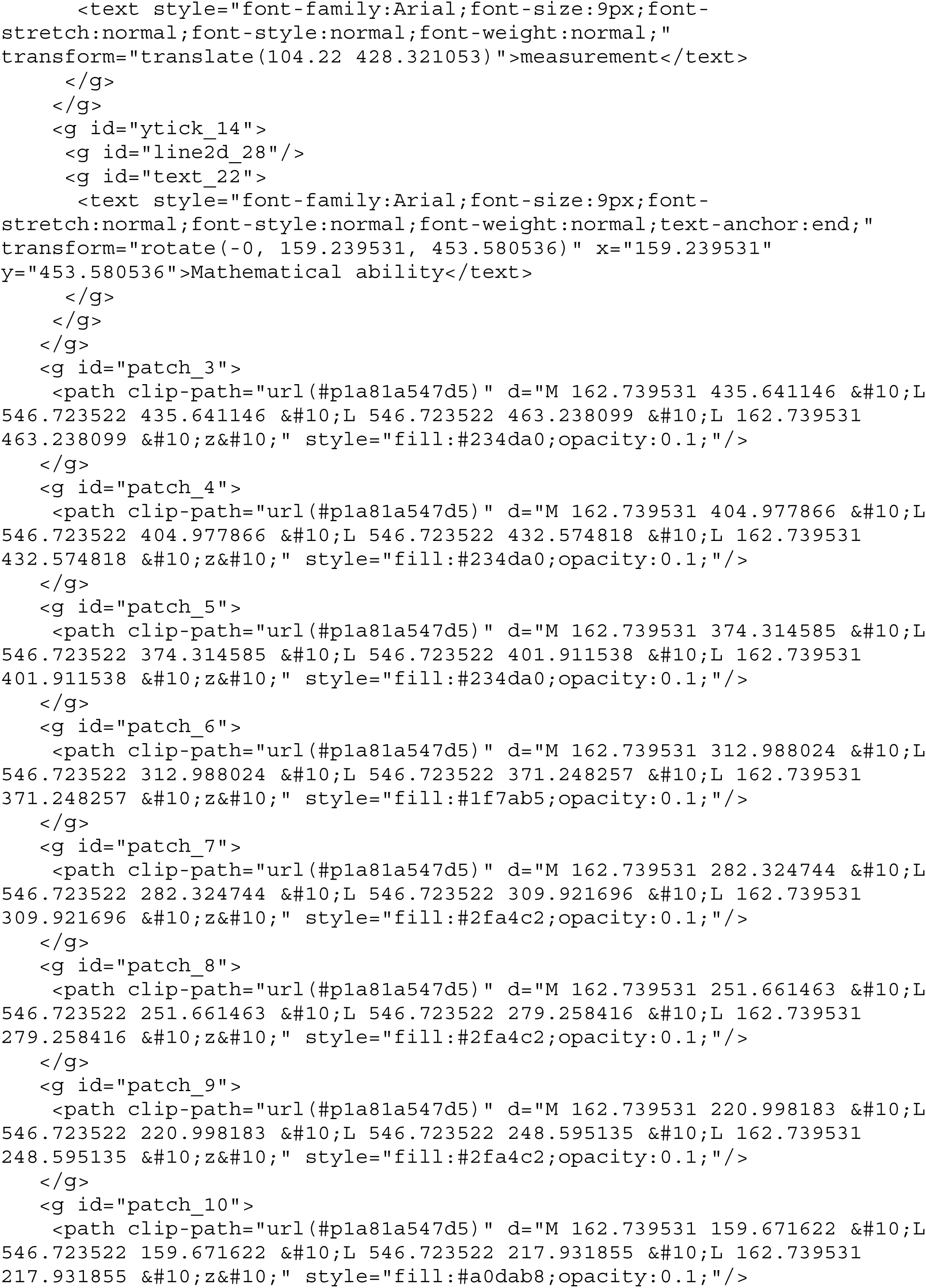

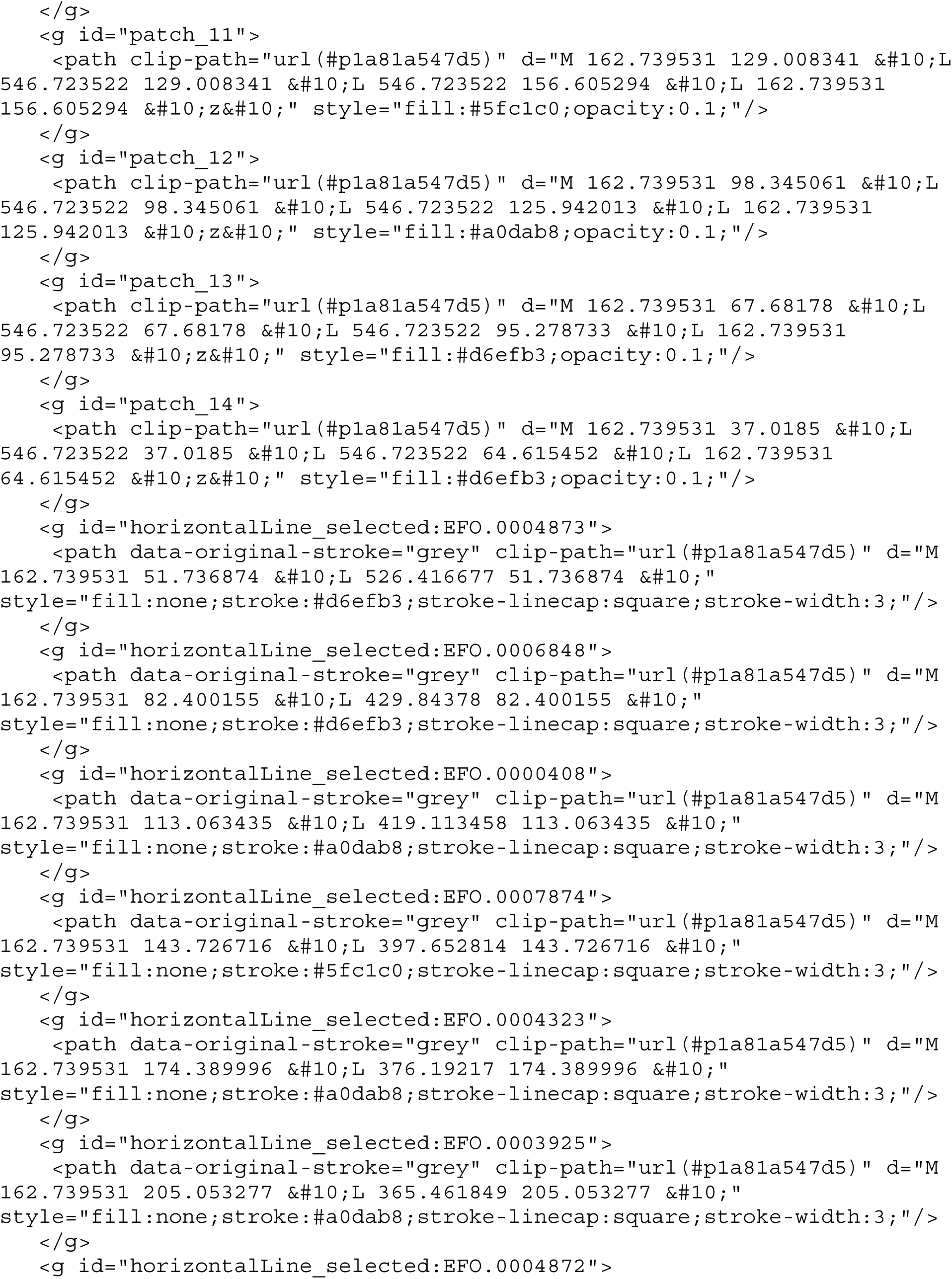

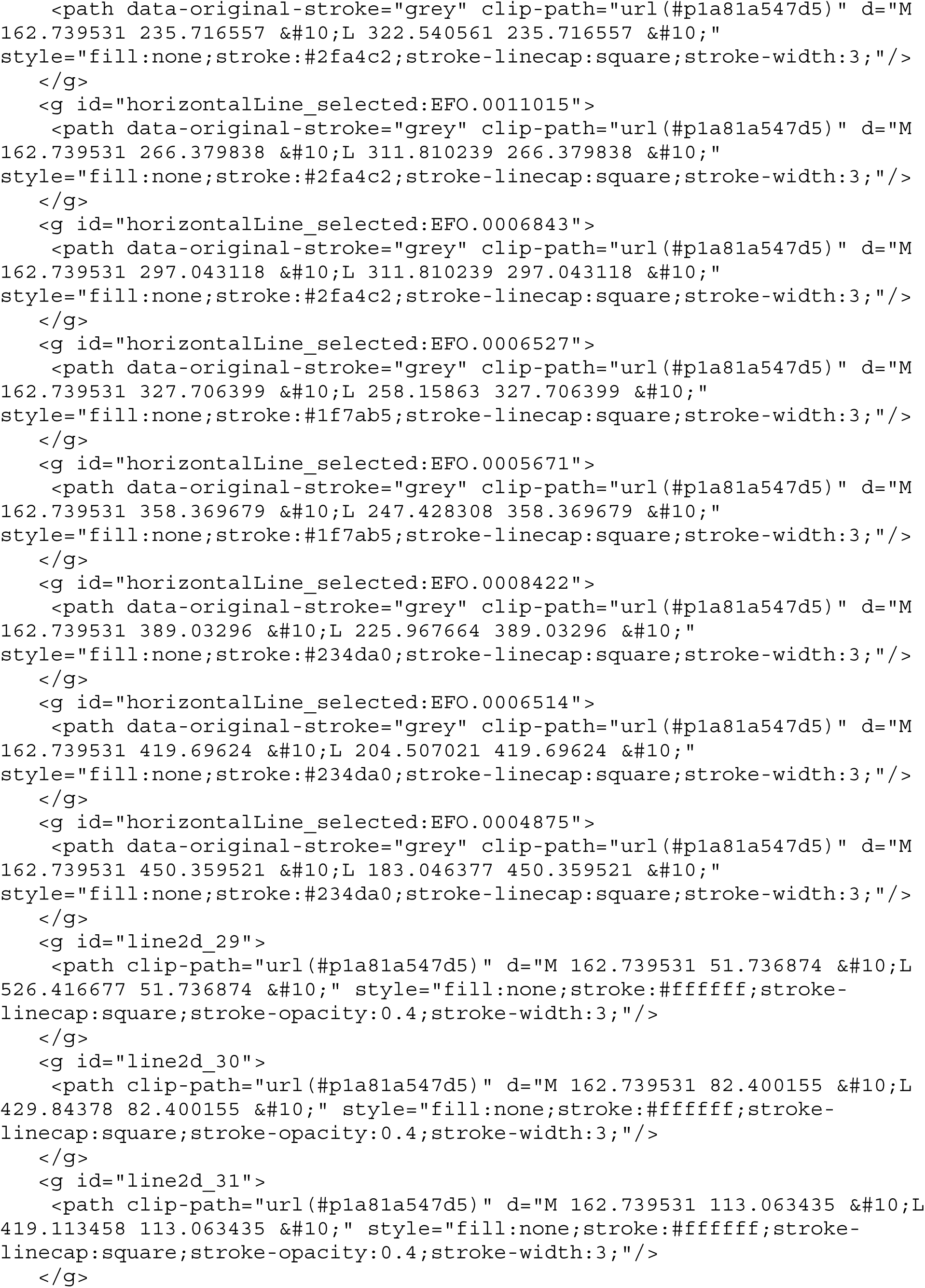

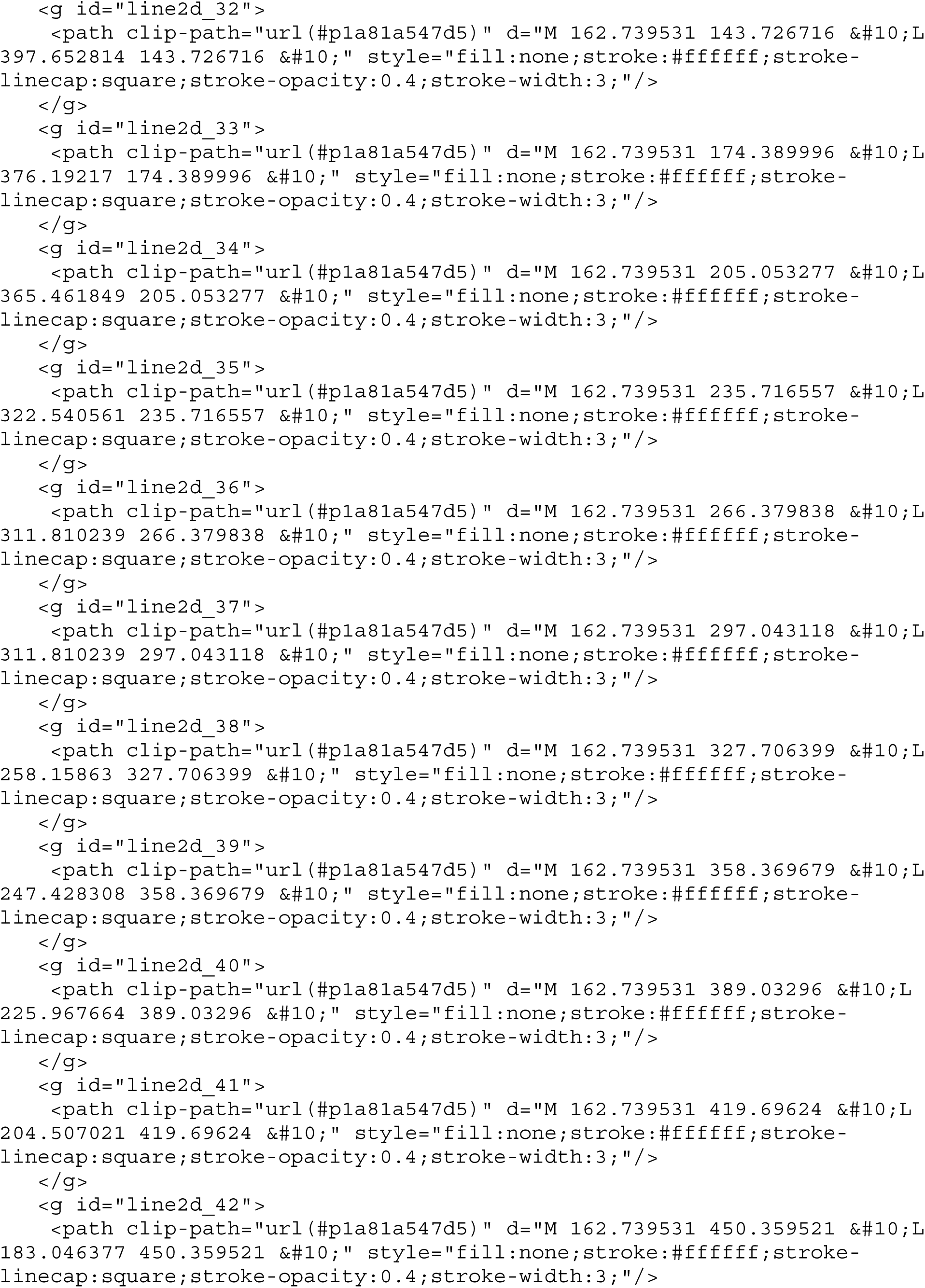

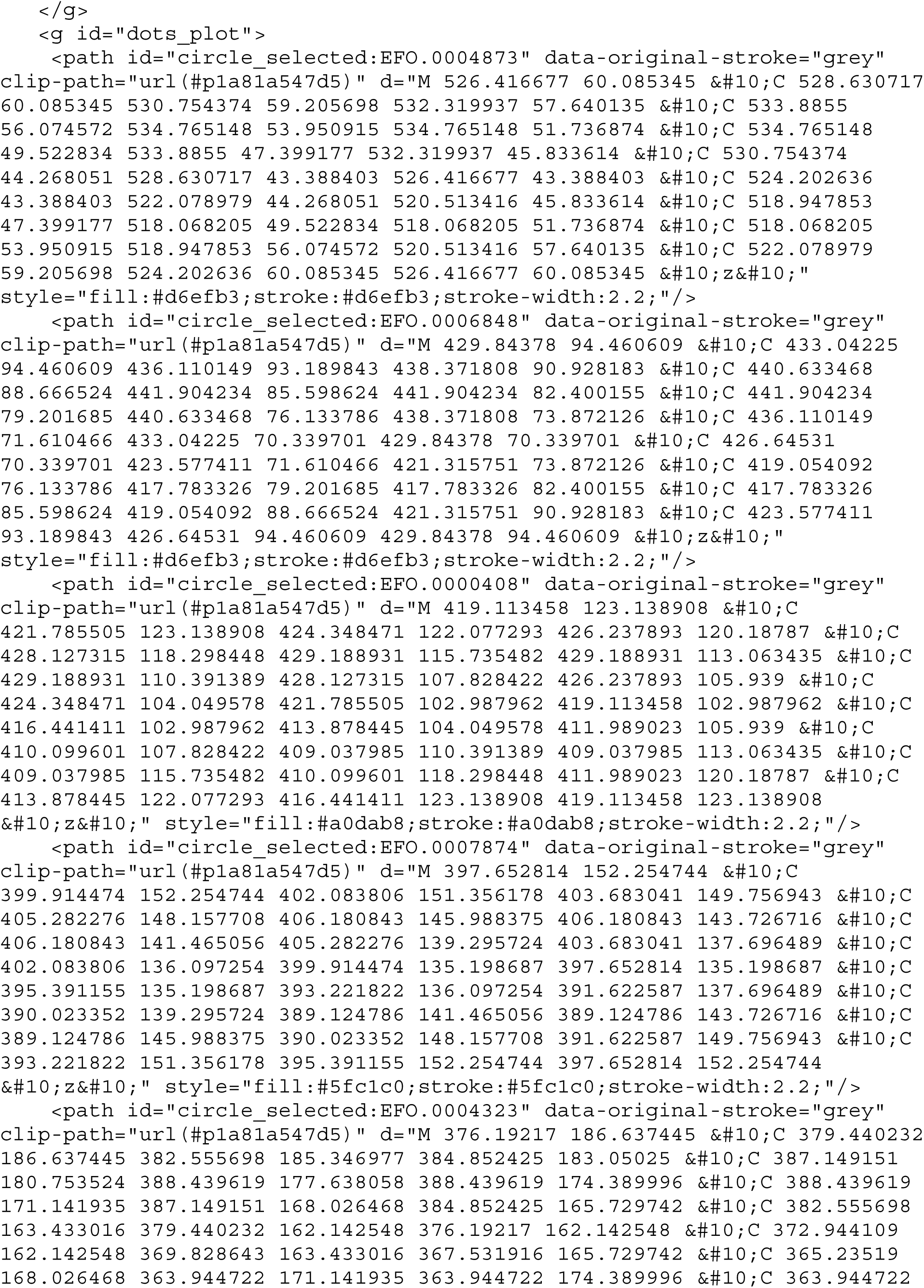

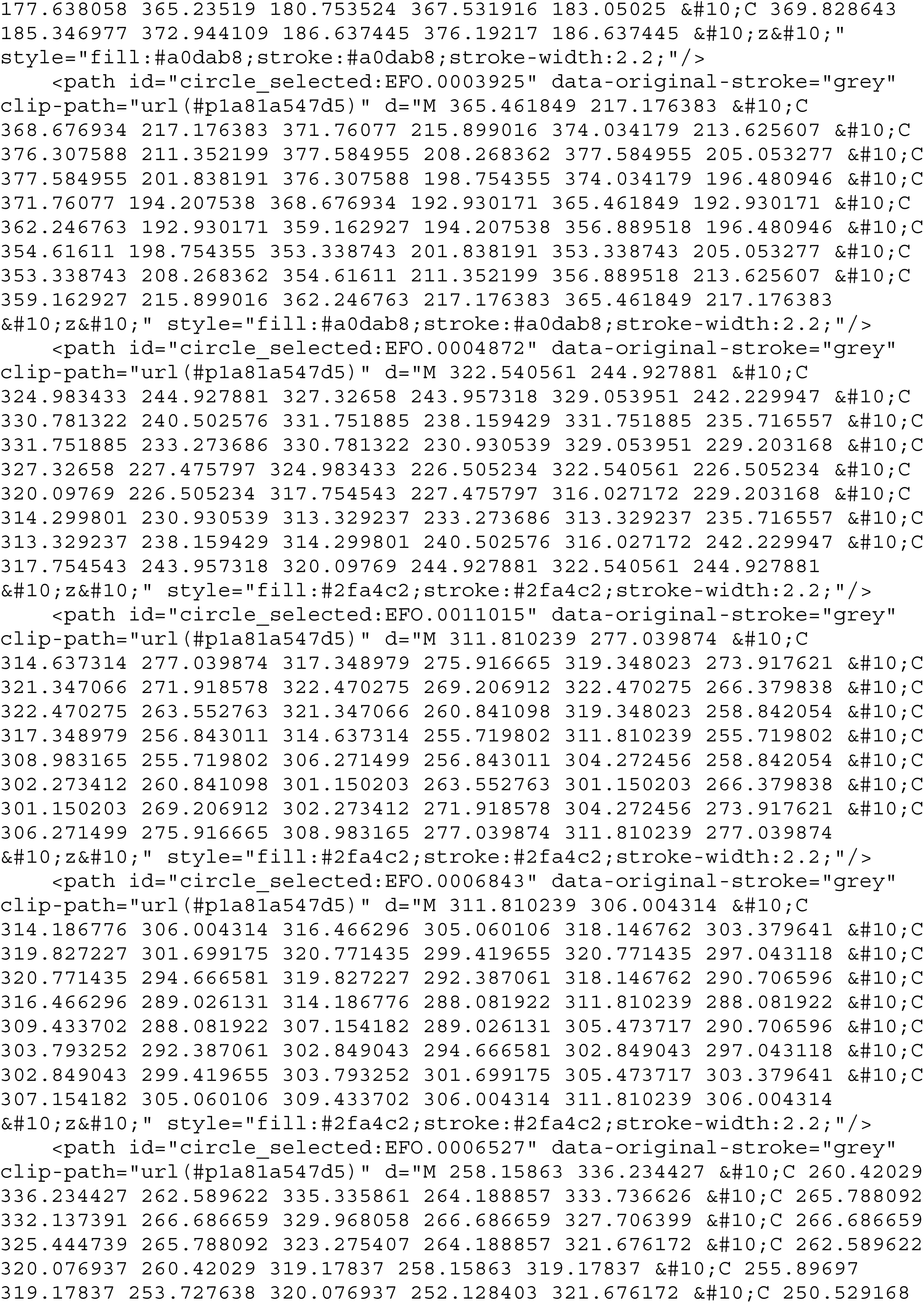

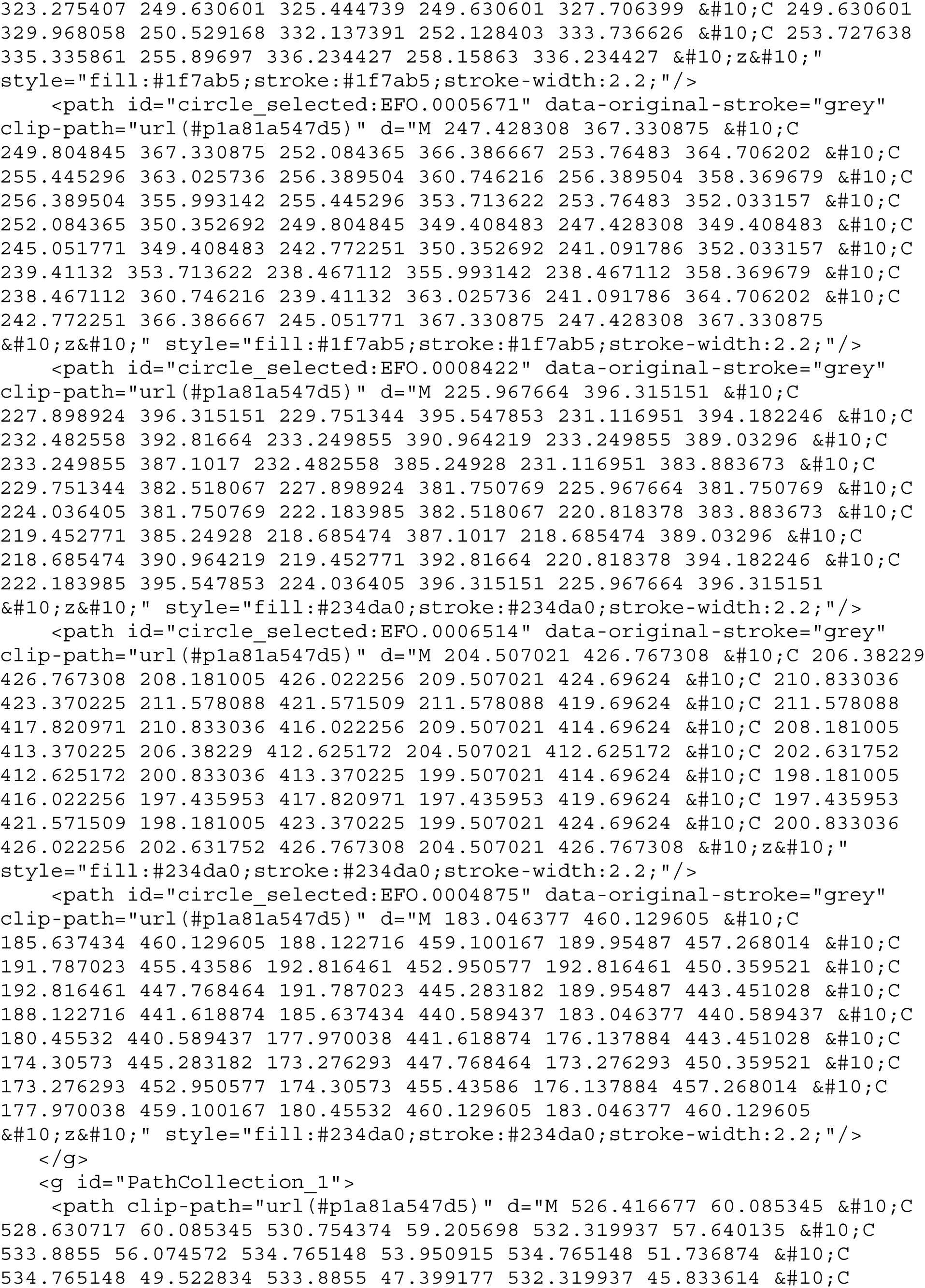

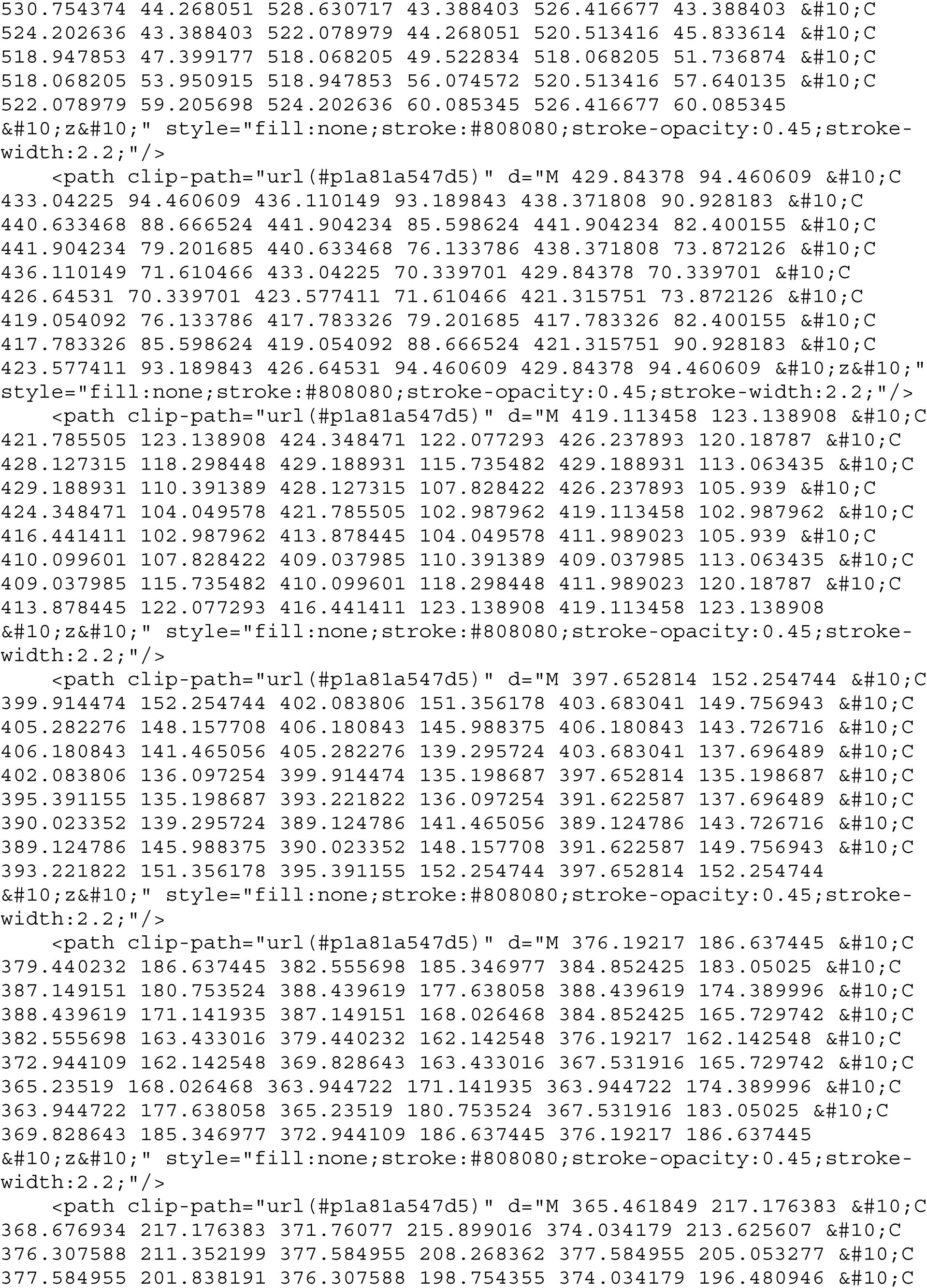

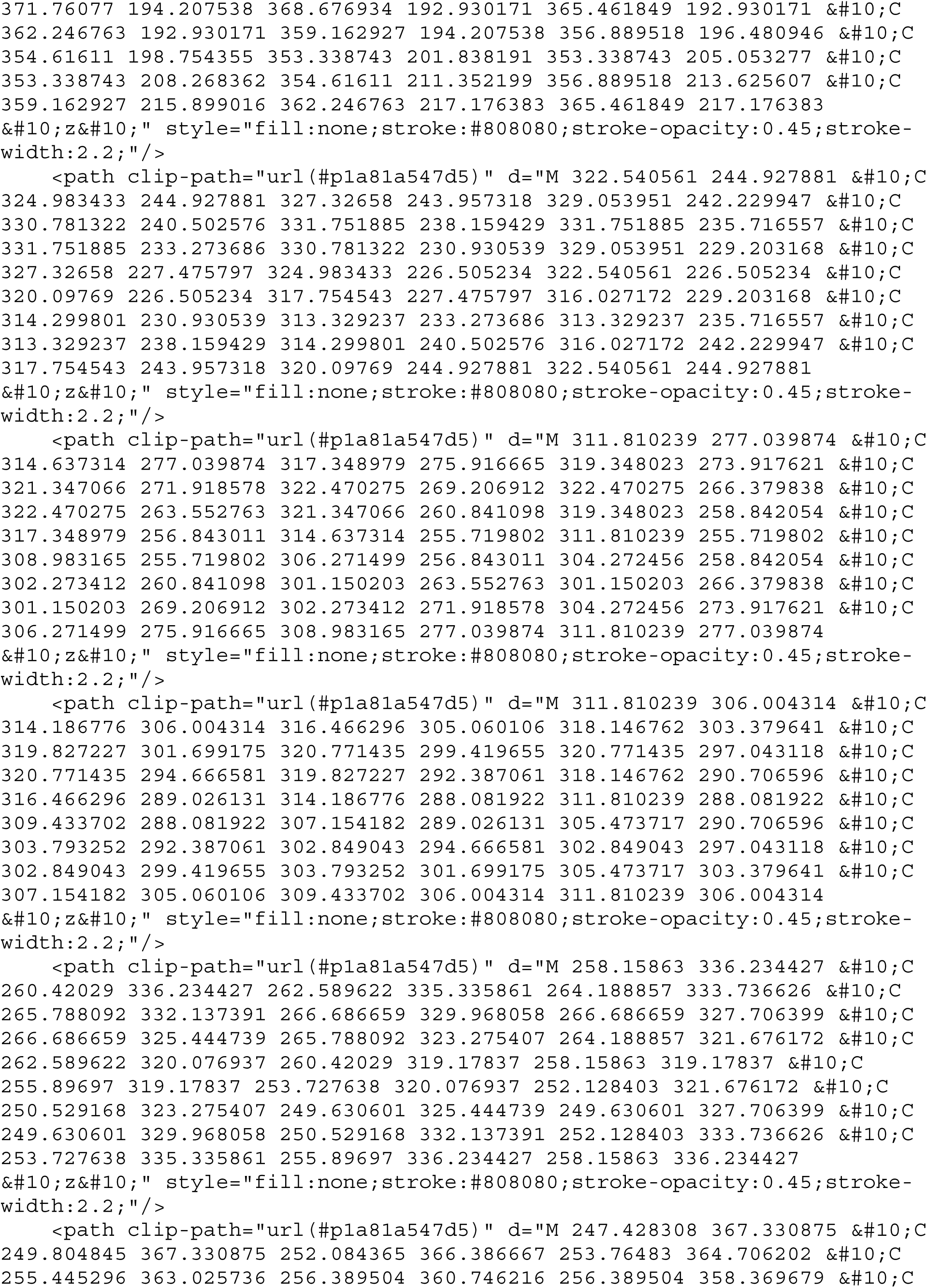

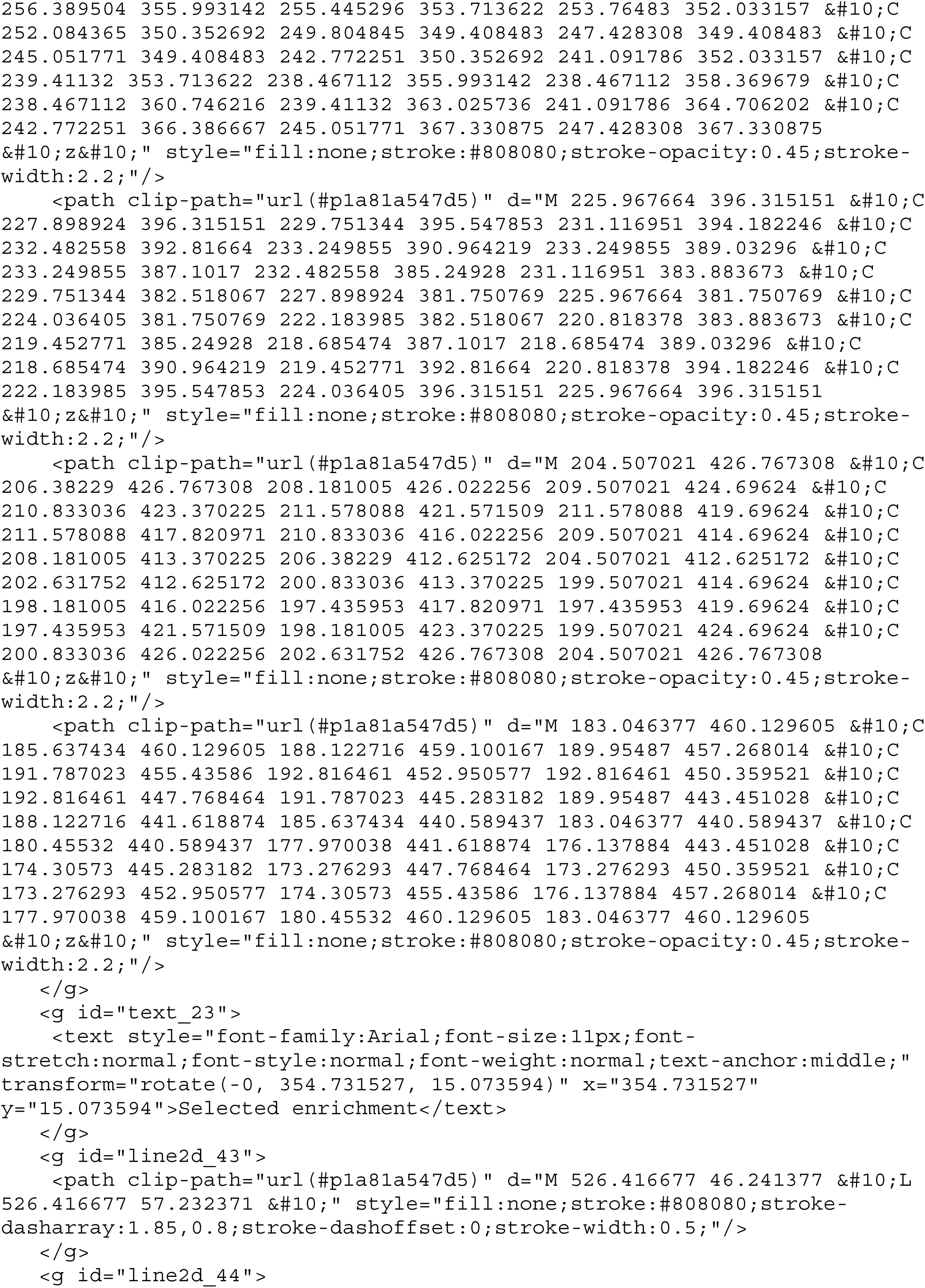

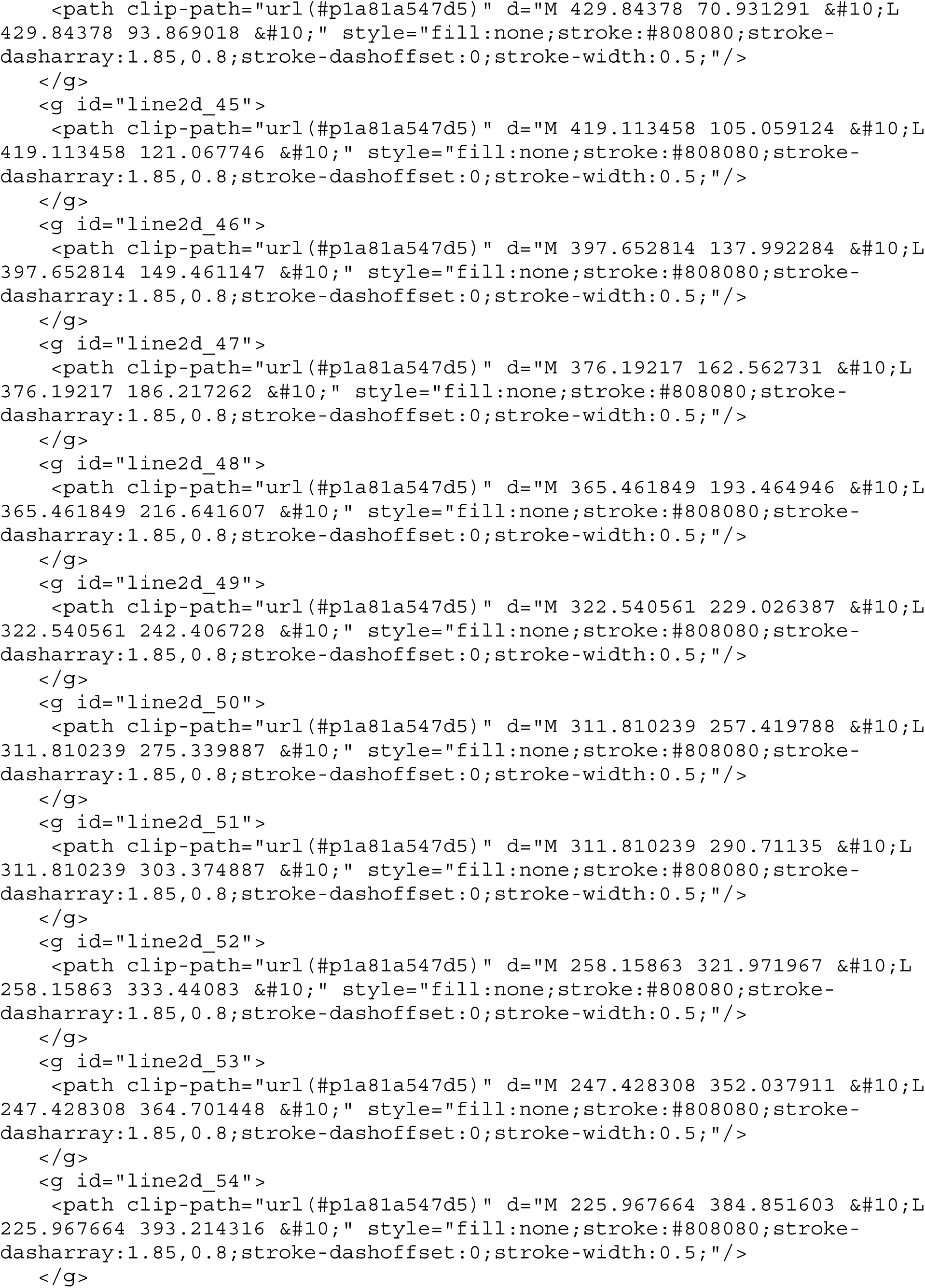

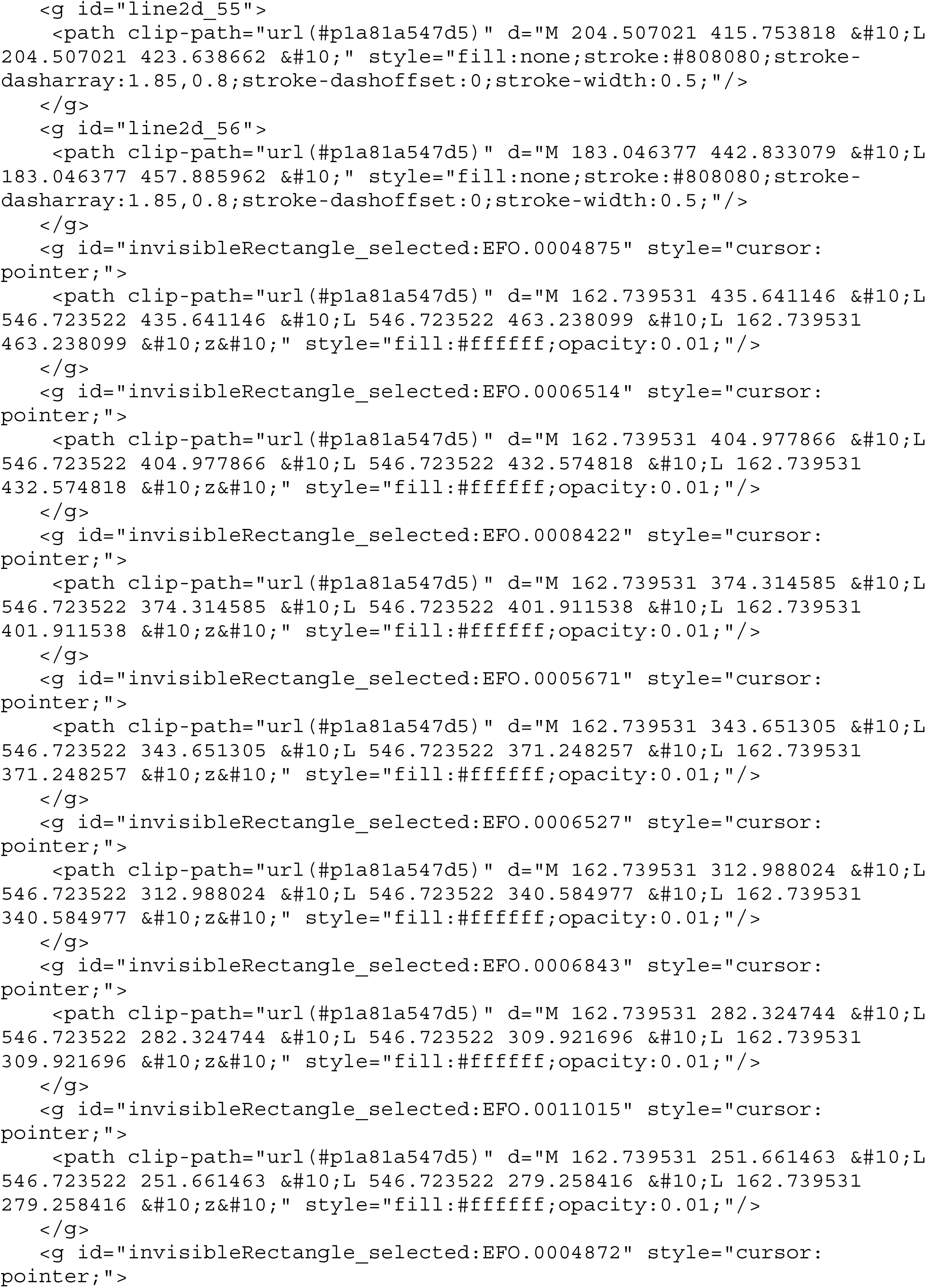

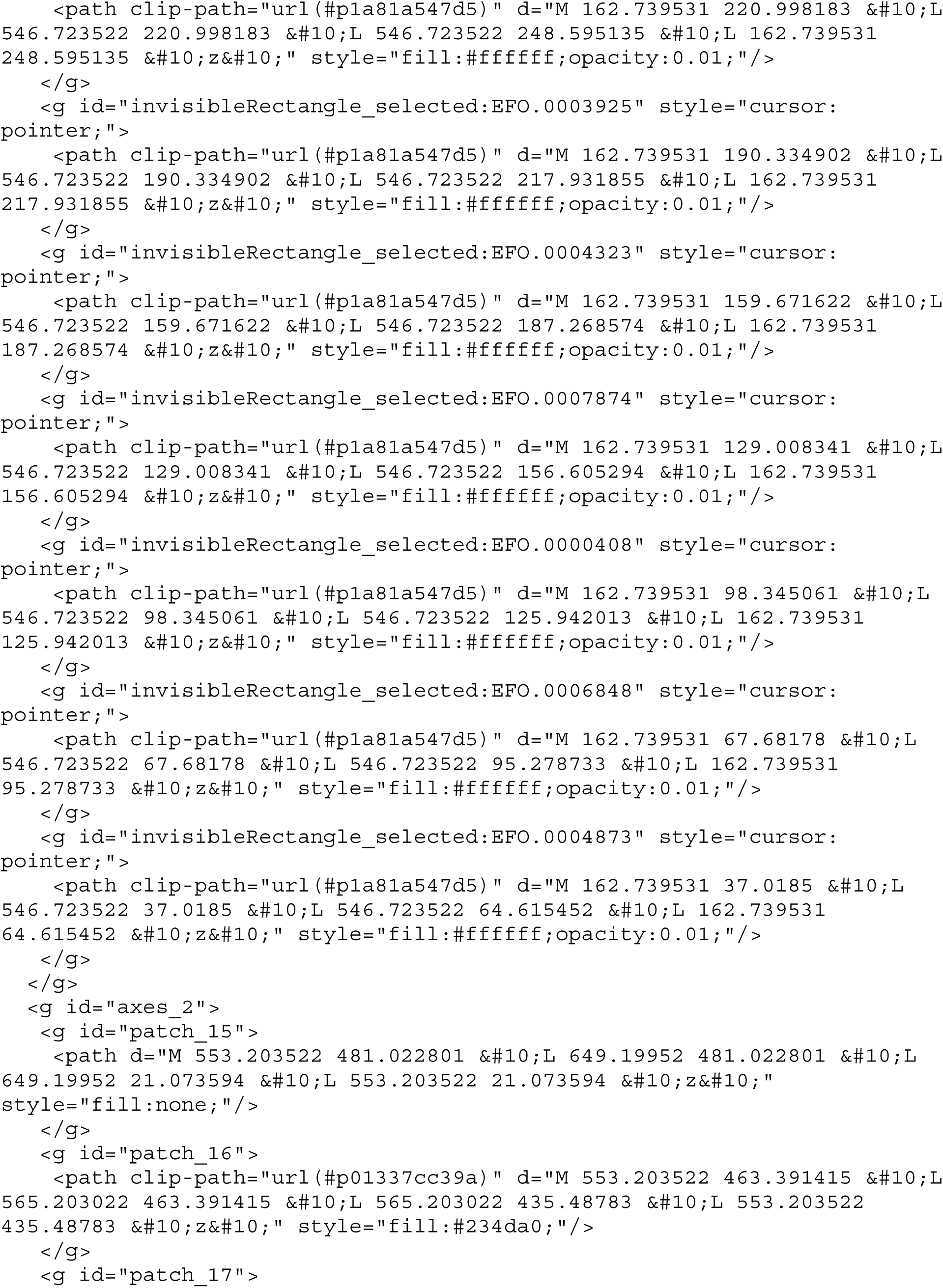

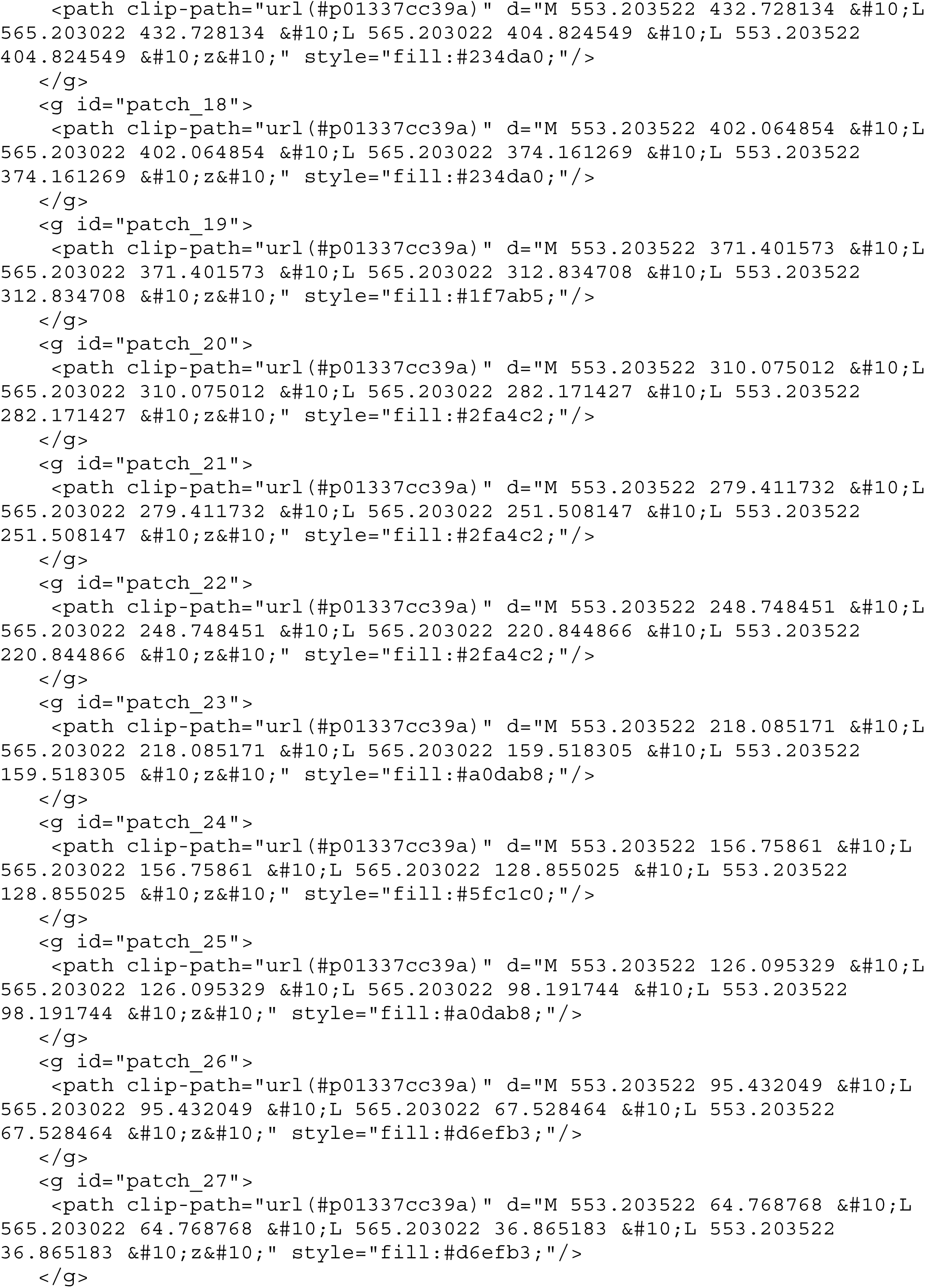

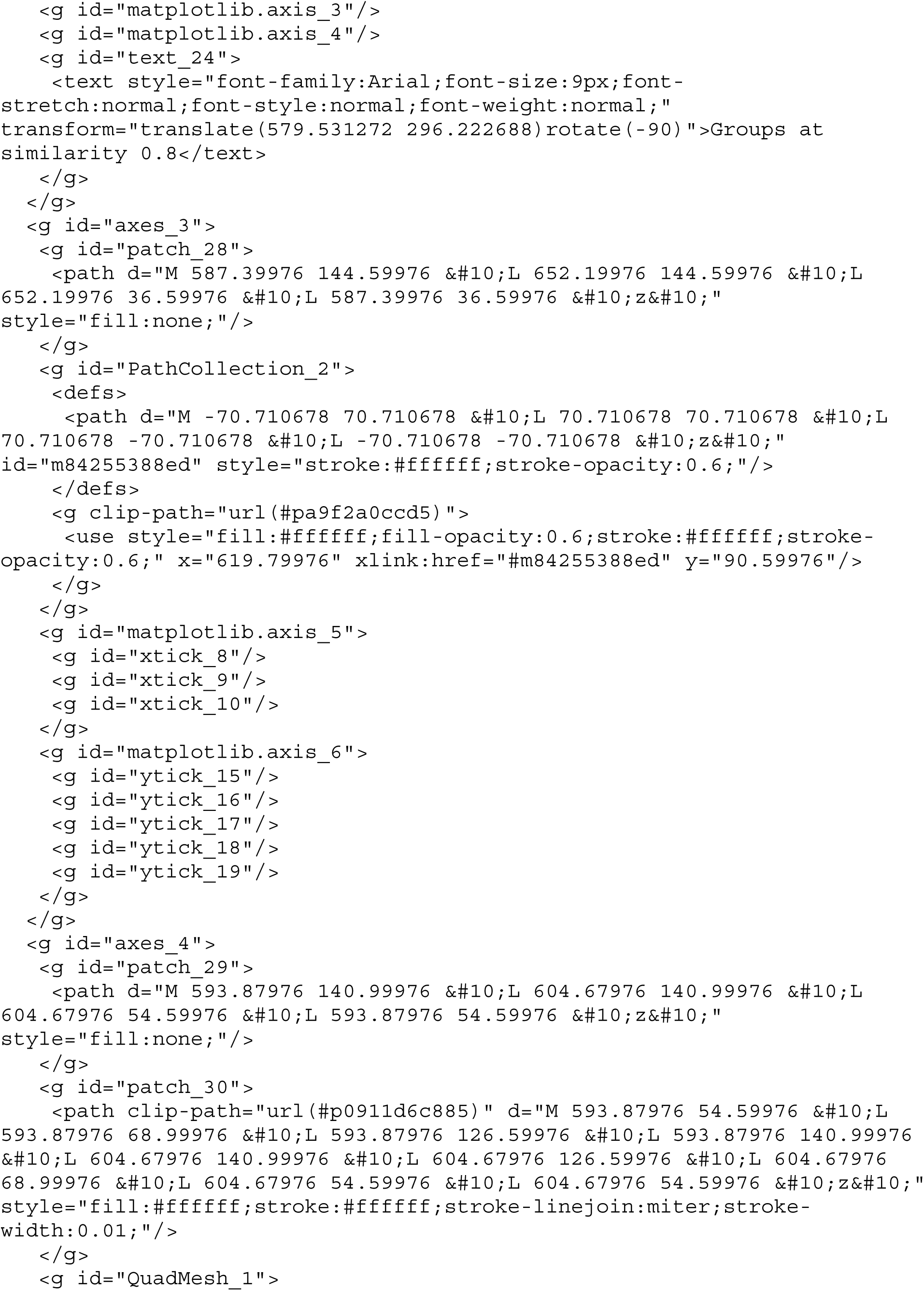

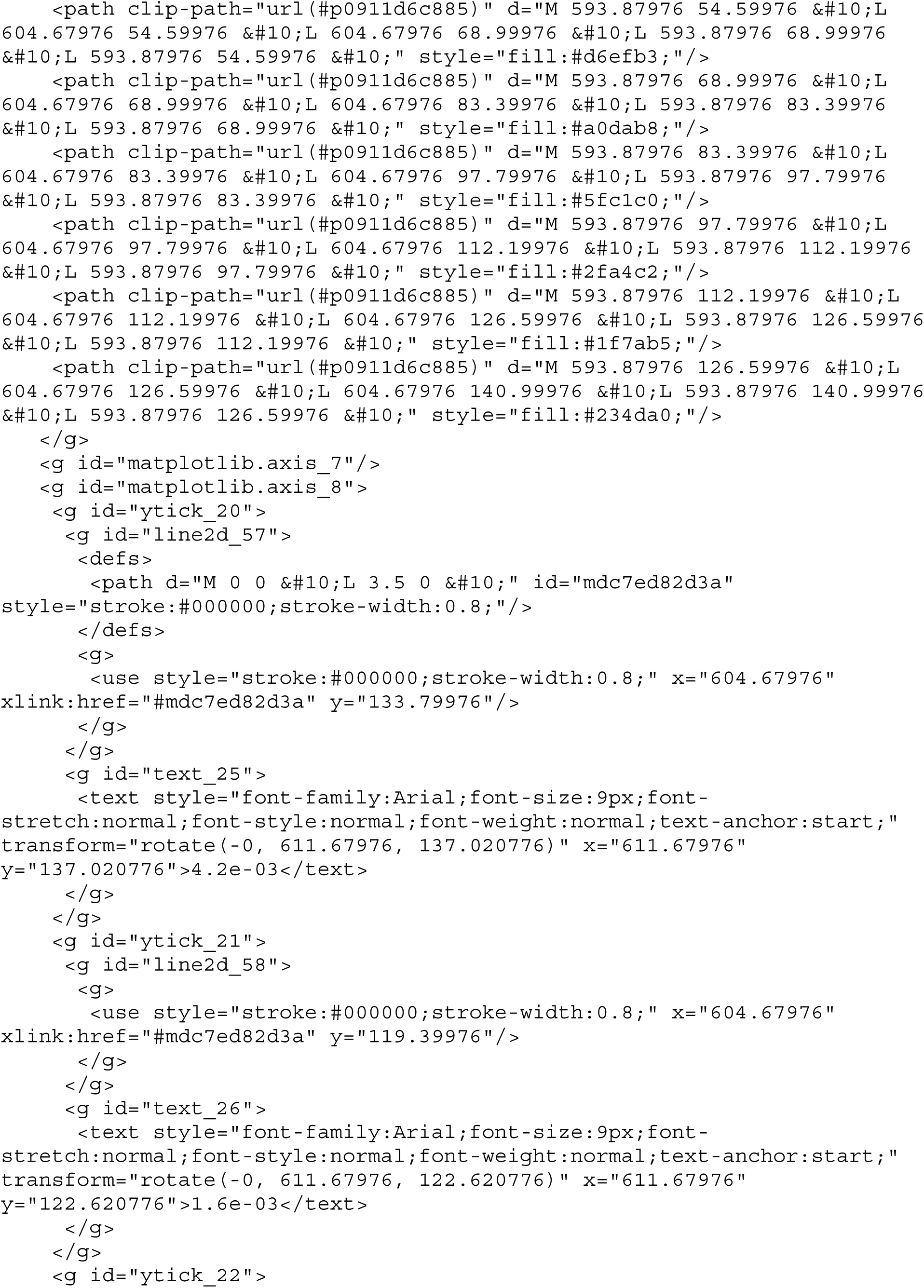

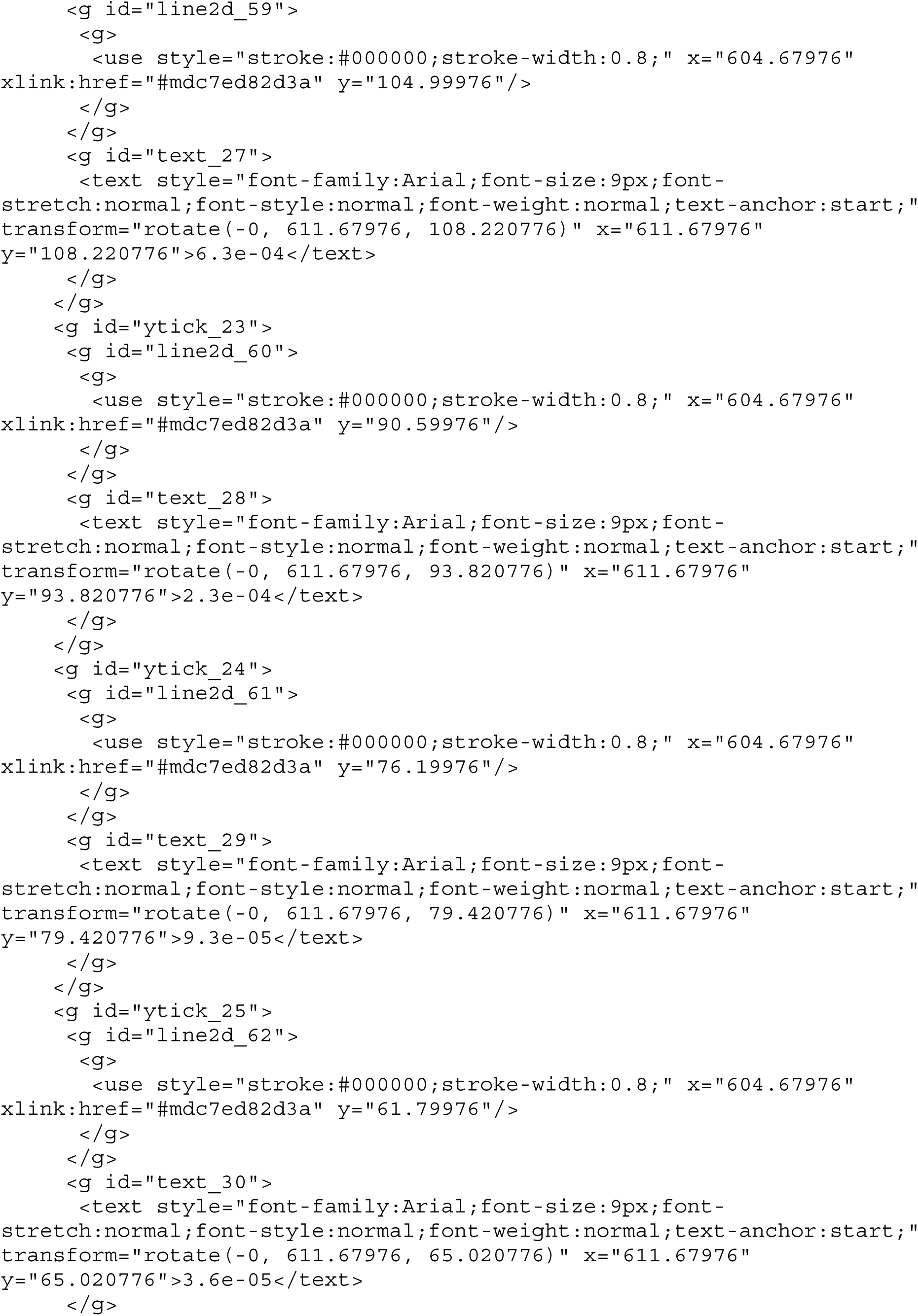

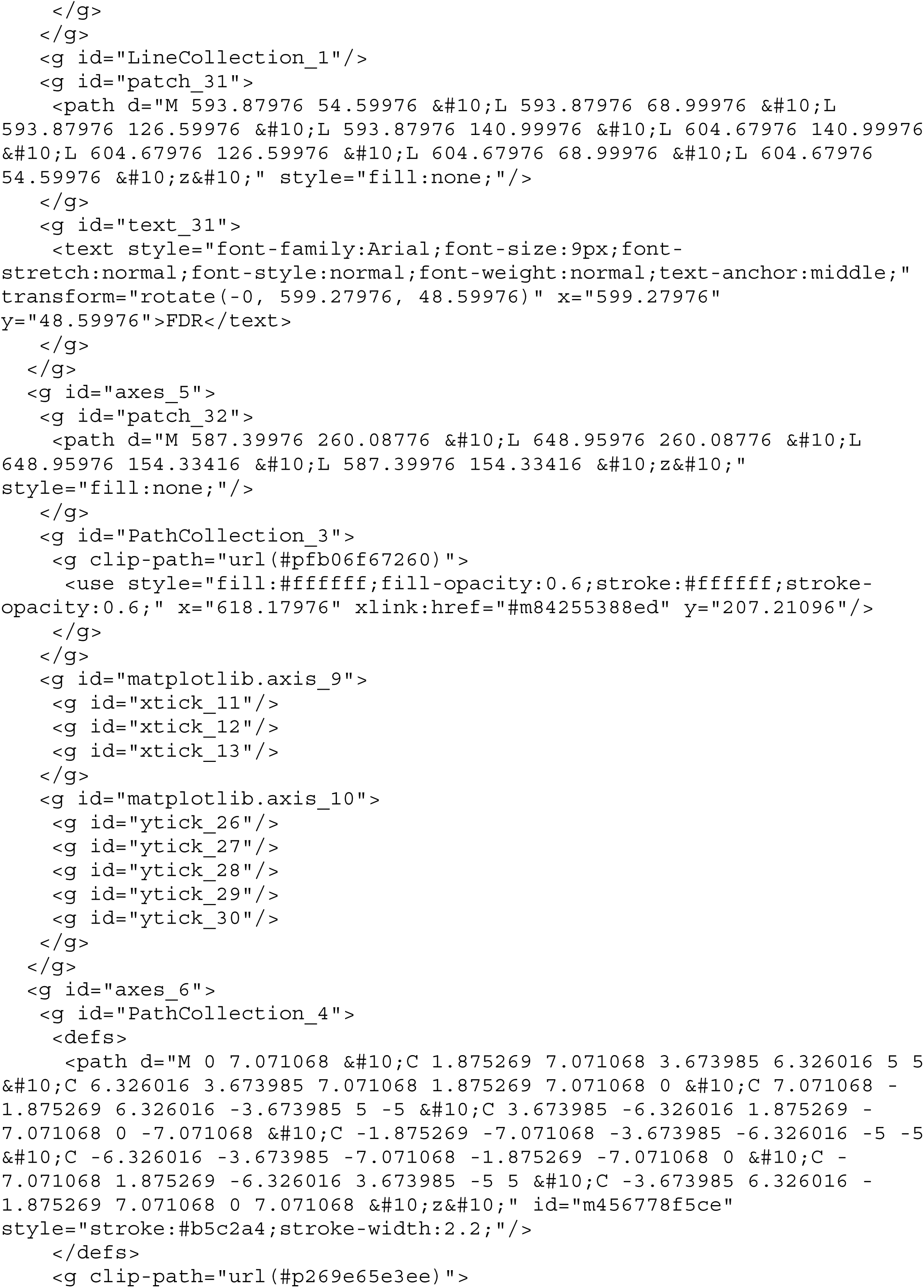

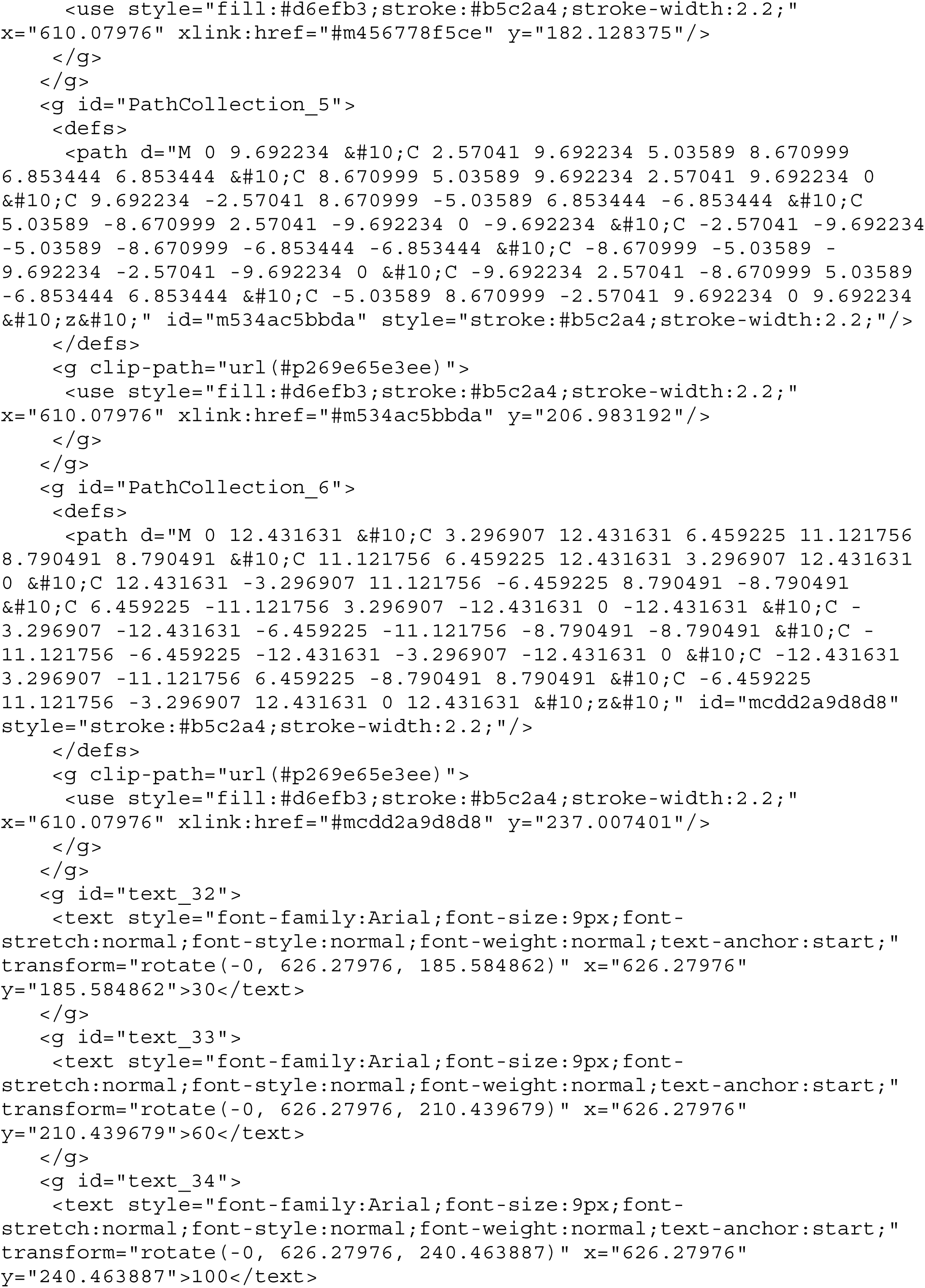

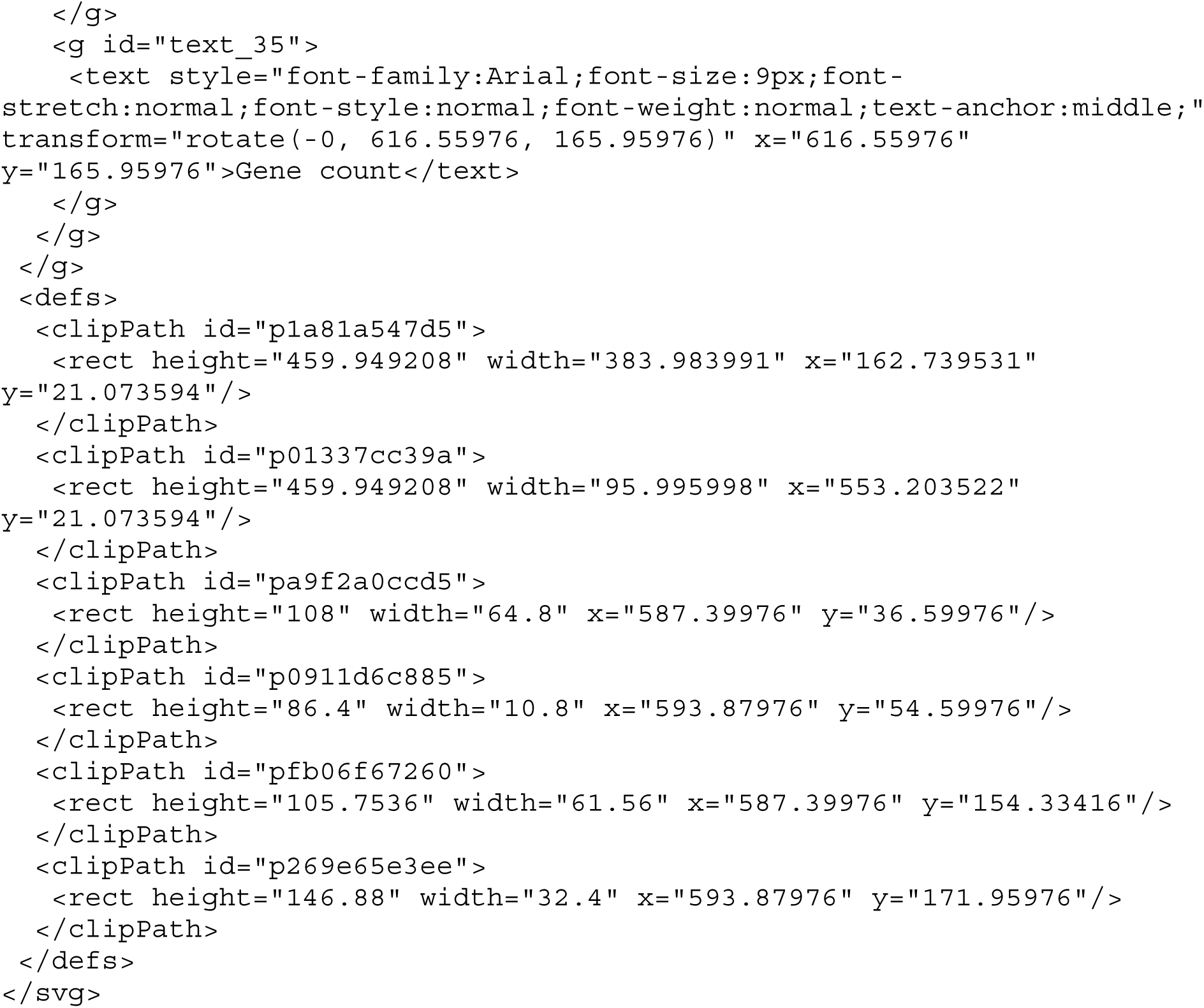

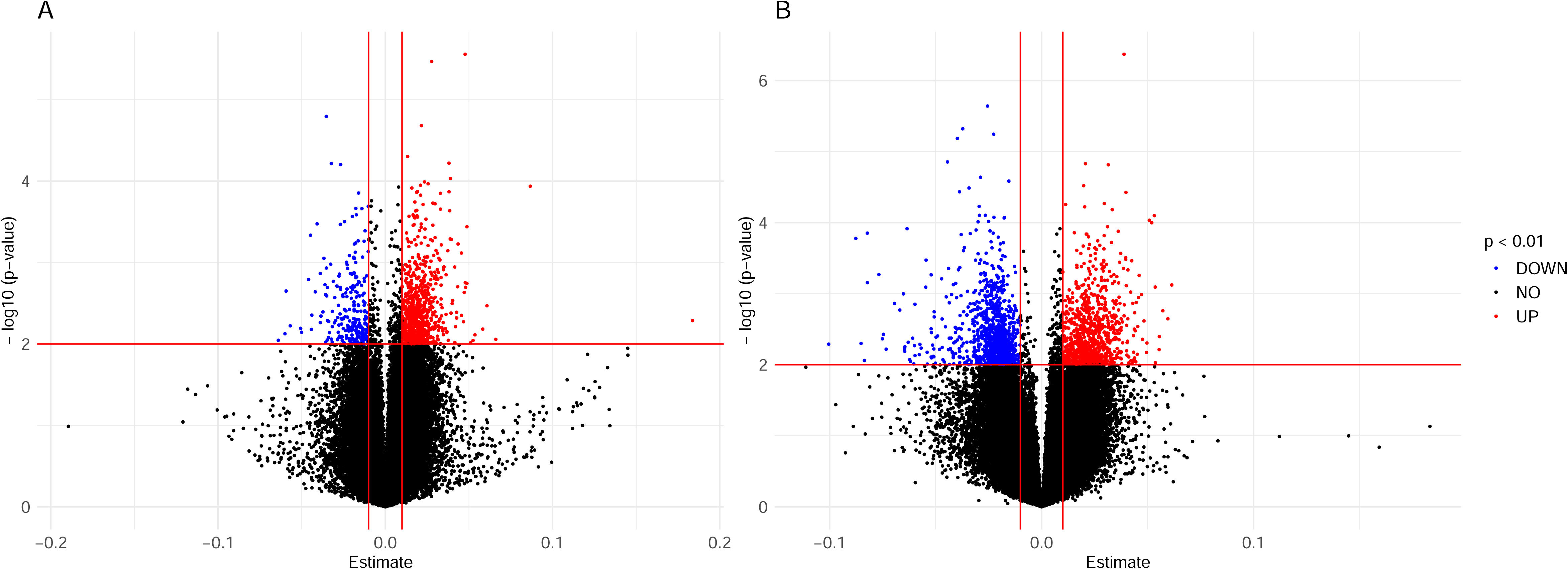

